# Quorum-Sensing Synchronization of Synthetic Toggle Switches: A Design based on Monotone Dynamical Systems Theory

**DOI:** 10.1101/024810

**Authors:** Evgeni V. Nikolaev, Eduardo D. Sontag

## Abstract

Synthetic constructs in biotechnology, biocomputing, and modern gene therapy interventions are often based on plasmids or transfected circuits which implement some form of “on-off” switch. For example, the expression of a protein used for therapeutic purposes might be triggered by the recognition of a specific combination of inducers (e.g., antigens), and memory of this event should be maintained across a cell population until a specific stimulus commands a coordinated shut-off. The robustness of such a design is hampered by molecular (“intrinsic”) or environmental (“extrinsic”) noise, which may lead to spontaneous changes of state in a subset of the population and is reflected in the bimodality of protein expression, as measured for example using flow cytometry. In this context, a “majority-vote” correction circuit, which brings deviant cells back into the desired state, is highly desirable, and quorum-sensing has been suggested as a way for cells to broadcast their states to the population as a whole so as to facilitate consensus.

In this paper, we propose what we believe is the first such design that has mathematically guaranteed properties of stability and auto-correction. Our approach is guided by concepts and theory from the field of “monotone” dynamical systems developed by M. Hirsch, H. Smith, and others. We benchmark our design by comparing it to an existing design which has been the subject of experimental and theoretical studies, illustrating its superiority in stability and self-correction of synchronization errors. Our stability analysis, based on dynamical systems theory, guarantees global convergence to steady states, ruling out unpredictable (“chaotic”) behaviors and even sustained oscillations. These results are valid no matter what are the values of parameters, and are based only on the wiring diagram. The theory is complemented by extensive computational bifurcation analysis, performed for a biochemically-detailed and biologically-relevant model that we developed. Another novel feature of our approach is that our theorems on stability of steady states for homogeneous or mixed populations are valid independently of the number of cells in the population, and depend only on the relative proportions of each type of state.

While monotone systems theory has been used previously for systems biology analysis, the current work illustrates its power for synthetic biology design, and thus has wider significance well beyond the application to the important problem of coordination of toggle switches.

**Author Summary:** For the last decade, outstanding progress has been made, and considerable practical experience has accumulated, in the construction of elementary genetic circuits that perform various tasks, such as memory storage and logical operations, in response to both exogenous and endogenous stimuli. Using modern molecular “plug-and-play” technologies, various (re-)programmable cellular populations can be engineered in a routine manner, and they can be combined into more complex cellular systems.

Among all engineered synthetic circuits, a *toggle*, a robust bistable switch leading to a binary response dynamics, is the simplest basic synthetic biology device, analogous to the “flip-flop” or latch in electronic design, and it plays a key role in biotechnology, biocomputing, and proposed gene therapies. However, despite many remarkable properties of the existing toggle designs, they must be tightly controlled in order to avoid spontaneous switching between different expression states (loss of long-term memory) or even the breakdown of stability through the generation of stable oscillations.

To address this concrete challenge, we have developed a new design for quorum-sensing synthetic toggles, based on monotone dynamical systems theory. Our design is endowed with strong theoretical guarantees that completely exclude unpredictable chaotic behaviors, as well as undesired stable oscillations, and leads to robust consensus states.

## Introduction

In the short period since the pioneering milestones in synthetic biology [1,2], outstanding progress has been made, and considerable practical experience has accumulated, in the construction of genetic circuits that perform various tasks, such as memory storage and logical operations, as well as support biomedical interventions and biotechnological manipulations in response to both exogenous and endogenous stimuli [3–6]. These circuits often include plasmids or transfected circuits which implement some form of “on-off” binary device, generically referred to as a *toggle switch*. For example, the expression of a protein used for gene therapy could be triggered by the recognition of some combination of inducers such as antigens, and memory of this event should be maintained across a cell population until a specific stimulus commands a coordinated shut-off [1,3,4]. In this context, as well as in many others, it is desirable for populations of cells to achieve coordinated static and/or dynamic functionalities. However, this coordination is hampered by molecular (“intrinsic”) or environmental (“extrinsic”) noise, which may lead to spontaneous changes of state in a subset of the population and is reflected in the bimodality of protein expression, as measured for example using flow cytometry.

To achieve robustness across a population, one may implement a “majority-vote” correction circuit that brings deviant cells back into the desired state. Much synthetic biology research focuses on single-cell microorganisms, often bacteria [4,6]. Bacterial populations are relatively simple, and their sociality in all its complexity can be rationally understood [7], providing a foundation for building more complex cellular systems. For bacteria, quorum-sensing (QS) has been suggested as a way for cells to broadcast their states to the population as a whole so as to facilitate consensus. QS signaling pathways [8] can, for example, regulate bacterial gene expression in response to fluctuations in cell-population density. Bacteria produce and release various signaling molecules called autoinducers (AIs) [8–11]. The detection of a minimal threshold stimulatory concentration of an AI leads to an alteration in the host’s gene expression. Both Gram-positive and Gram-negative bacteria use QS communication to regulate a diverse array of physiological activities. Synthetic biology design has adopted QS communication in its toolbox [12], because natural and artificially engineered QS modules can be used to interface synthetic circuits with exogenous and endogenous cues [4], and a systematic modular approach to standardize engineering toggle genetic circuits that would allow programmed cells to be designed for various specific purposes and to communicate their states to other cells was suggested as a bioengineering “plug-and-play” modular approach [4]. The design of such QS-toggle combinations is the focus of this paper.

### A known design and its drawbacks

Kobayashi et al [4] consider a genetic toggle switch, interfaced with a QS signaling pathway. Their *E. coli* strain “B2” detects as well as produces (through the synthetase encoded by the expressed gene luxI, which converts common precursor metabolites) acyl-homoserine lactone (AHL) signaling molecules. AHL is a QS signaling pathway from *Vibrio fischeri*. Toggle B2 (Fig 1) enables an *E. coli* population to measure population density through AHL, because AHL signaling can be reversibly transported to the medium via diffusion, contributing to the AHL density in the culture [4].

To achieve an in-depth understanding of dynamic properties of coupled QS and toggle constructs, Kuznetsov et al. [13] developed and studied a mechanistic mathematical model of a population (or, equivalently, an ensemble) comprising *N* toggles, see Fig. 2 (bottom panel), corresponding to Toggle B2 (Fig. 1), *N* > 1. Their study revealed important multiple functions, namely bistability as well as stable oscillations, that an ensemble of Toggles B2 was capable of exhibiting. Analytical conditions for bistability were found, and a time separation was introduced to obtain a stable limit cycle for a population of interacting cells.

**Figure 1.**
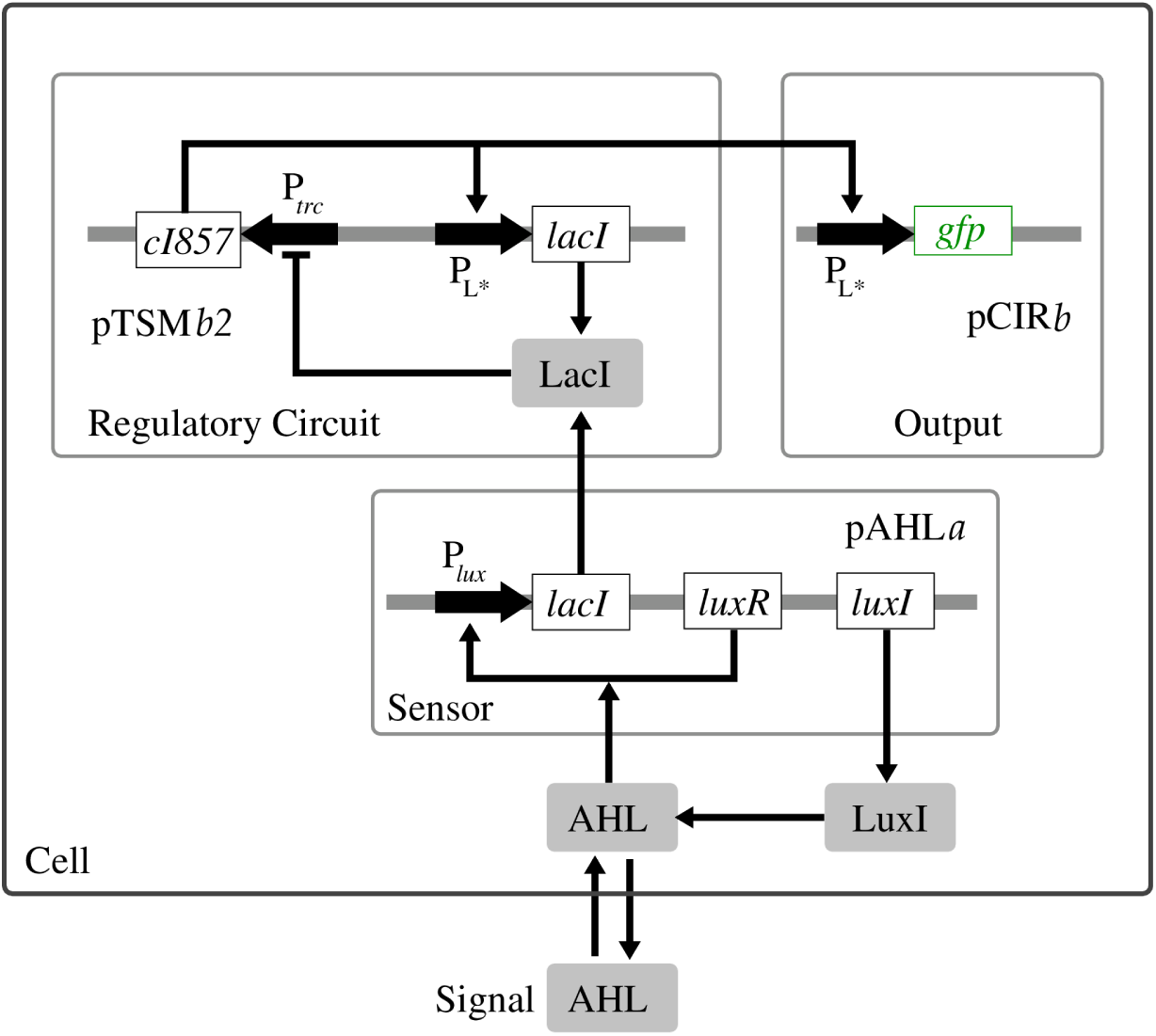
Toggle B2: Density-Dependent Gene Activation. *Notations and abbreviations*. Plasmids: pTSM*b2*, pCIR*b* and pAHL*a*; genes: *gfp*, *cI857*, *lacI*, *luxR*, and *luxI* ; promoters: P_*trc*_, P_L_*\**, and P_*lux*_; AHL, acyl-homoserine lactone. Figure adapted from Fig. 6(A) in [4].

In bistable circuits (toggles), transitions such as those caused by fluctuations due to low copy numbers of species per cell, or due to local environmental “noise” can force individual cells to change expression state at random [4]. This noise effect can spontaneously lead to the emergence of heterogeneous (mixed) populations consisting of cells in different expression states, which appear as bimodal population distributions when the corresponding protein levels are measured [4]. To investigate the effect of a spontaneous toggle switching in single and coupled cellular systems, leading to bimodal population distributions, Wang et al. [14] developed models for a single cell and a multi-cellular toggle system comprising *N* cells, respectively. In their models, the dynamics of the repressor proteins LacI and *λ CI* is described by the two ODE equations developed in [1]. The AI-interfacing employed in the population model [14] corresponds to a signaling pathway which is slightly different from the signaling pathway in Toggle B2 (Fig. 1) as suggested in [4] and described earlier. The main difference is in the description of the expression of the gene that encodes LuxI, see [14] for more details.

Despite the remarkable properties of design B2 and its modifications, observed experimentally in controllable experimental settings [4], and studied theoretically [13,14], the fact that their functional repertoire includes not only a bistable long-term memory but also the generation of stable oscillations suggests that the environment-toggle system must be tightly controlled in order to avoid spontaneous switching, not merely between different expression states but even between different functions.

To address this challenge, we propose a novel design, which is endowed with mathematically guaranteed properties of stability and auto-correction. Our approach is closely guided by concepts and theory from the powerful framework of *monotone dynamical systems* pioneered by Hirsch and H. Smith [15–21].

We benchmark our design by comparing it to Toggle B2, which, as discussed, has been subject of experimental and theoretical studies, illustrating its superiority in stability and self-correction of synchronization errors. We employ monotone theory to provide guarantees of global convergence to steady states, thus ruling out unpredictable (“chaotic”) behaviors and sustained oscillations. These theorems are valid no matter for all values of parameters and are based only on the network structure. We also provide an extensive computational bifurcation analysis for a biochemically-detailed and biologically-relevant. Our results for homogeneous or mixed populations are valid independently of the number of cells in the population, and depend only on the relative proportions of each type of state.

### The components

As a basic design, we chose a genetic toggle switch consisting of two mutually repressing genes, *lacI* and *tetR* [1]. *We use two acylated homoserine lactones (Acyl-HSLs), (i) N* -butanoyl-l-homoserine lactone (C4-HSL) secreted by *Pseudomonas aeruginosa* [22], *and (ii) N* -(3-hydroxy-7-*cis*-tetradecenoyl)-L-homoserine lactone (3-OH-C14-HSL) produced by *Rhizobium leguminosarum* [10] as a means of coordinating toggle-host activity. Our design has two QS arms built-in the toggle in such a way that each promoter-repressor pair is controlled by its own QS signaling pathway symmetrically. Because of this “mirror-like” toggle symmetry, we call our design a symmetric toggle or an “S” design.

To benchmark the new S toggle design and the monotone systems approach, we compare the S design to the well-studied asymmetric B2-strain (Fig. 1) which has one QS arm only [4,13]. In this work, we call the asymmetric B2-strain the “A” design. Our S design cannot be reduced to the A design by removing one QS arm, and, thus, the S design cannot be viewed as a straightforward extension of the A design. From a theoretical standpoint, it is worth remarking that the A design is non-monotone.

The S vs. A toggle comparative results obtained in this work can be summarized as follows:

- The monotone S toggle design completely excludes any unpredictable chaotic behaviors, as well as stable oscillations. Typical trajectories converge globally to stable equilibria. This conclusion is valid for all parameter values, and provides a strong theoretical guarantee missing from other synthetic biology designs.
- We refer to mixed states leading to bimodal distributions as *spontaneous synchronization* errors. We find that the S toggle design is able to self-correct (or, auto-correct) synchronization errors, while the non-monotone A toggle design is not.
- We show how monotone systems theory can predict not only the dynamics of an S toggle population but it also explains certain monotonically increasing or decreasing parametric dependencies of population steady states. We show that some of these predictions can facilitate self-synchronization and, thereby, reduce any chance for synchronization errors to emerge spontaneously.

### Organization of paper

In Models and Methods, the S toggle and A toggle mathematical models are introduced. The basic formalism and fundamental mathematical results of monotone systems theory, including Strong Monotonicity and Hirsch’s Theorem [15–19,21] are reviewed there as well. Balanced graphs, relation to graph partitions, and order presentation by flows are explained there. Additionally, reference values of dimensionless parameters, a scaling procedure, and the selection and interpretation of bifurcation parameters are discussed, and the mirror symmetry of the S toggle is formalized. The symmetry will be used to explain symmetry breaking (pitchfork) bifurcations in the S toggle. We also formalize a concept of spontaneous synchronization errors. Here, three types of equilibrium populations are considered: One homogeneous population, and two heterogeneous (mixed) populations (bimodal distributions) with both equally (1:1)-mixed and not-equally (*N*_1_:*N*_2_)-mixed transcriptional signatures, *N*_2_ ≪ *N*_1_, the latter giving rise to spontaneous synchronization errors, where *N* = *N*_1_ + *N*_2_, and *N* is the number of cells in the given population.

In Results and Discussion, we proceed to a comparative theoretical and computational analysis of the S toggle and A toggle design models. We begin this section with results on the application of monotone systems theory to the S design, as these results constitute the main conceptual and practical subject motivating this work (Application of Monotone Systems Theory to the S design). We start by explaining how monotone systems theory allows one to predict, based on qualitative knowledge only, that generically all solutions converge to equilibria, with no possible oscillations [13] nor chaotic behavior [23], no matter what the kinetic parameters are. This is in contrast to the A design, which may admit oscillations [13]. Next, we analyze single S and A toggles decoupled from the environment (Bistability in Single S-Design and A-Design Toggles Isolated from the Environment), and observe that the S toggle remains bistable even if “redundant” repressor genes are removed from the corresponding plasmids. To show how the S design is more robust than the A design, we carry out a comparative bifurcation analysis of populations consisting of coupled S or A toggles. We select a free (bifurcation) dimensionless parameter which can be interpreted in terms of experimental interventions [6] leading to *(a)* changes in the membrane permeability, or *(b)* changes in the half-lives of repressor proteins, *(c)* or changes in the specific growth rate of the host cell. We additionally test the toggle design capabilities to self-correct spontaneous synchronization errors by sampling the basin of attractor of the corresponding equilibrium solutions. We find that the S toggle design is able to self-correct synchronization errors far better than the A toggle design.

The paper also has three Supplemental Information (SI) materials. In S1 Model Derivation, we derive mathematical models and carry out a nondimensionalization (scaling) procedure, the conclusions of which are used in the main text (Scaling). In S2 Estimation of Parameter Values, we discuss ranges of biologically meaningful parameter values based on data available in the existing literature. Values of biologically meaningful parameters depend upon experimental conditions and other factors controlled by an experimenter, as reviewed in [6]. Therefore, we provide an example of a concrete estimation of values of dimensionless parameters, which we interpret in terms of interventions reviewed in [6]. Since bistability only holds for ranges of these dimensionless parameters, to increase the chances that both S and A-synthetic toggles will function in the bistable regime, we have tuned the estimated values of reference dimensionless parameters to pin a region in the parameter space from which we choose appropriate parameters values for subsequent modeling studies. In S3 Exponential Stability of Cellular Populations, we prove a number of general theorems to analyze exponential stability [24] of both homogeneous and heterogeneous (mixed) population equilibrium states, independently of the number *N* of cells in the given population, which (*i.e.,* the value of *N* ≥ 2) can be *a priori* unknown.

## Models and Methods

Although our main objective in this paper is to present a conceptual and general organizing principle for the construction of self-correcting “majority-vote” multistable synthetic systems, we instantiate our ideas through a very concrete set of genes and protein products, all being standard molecular parts in synthetic biology [1,2,4,25–29]. We do that in order to emphasize the fact that our constructs can be realistically implemented with currently available molecular components. However, replacing these components with others does not change the basic mathematical framework.

To facilitate a conceptual and quantitative comparison of the S and A toggle designs, the corresponding genetic circuits are assumed to be built from the same tightly controlled *lac-tet* transcription entities [30–35], which have been intensively used in a number of experimental and theoretic-modeling studies in the context of synthetic biology [1,2,4,26–29]. Below, we briefly characterize relevant molecular details and then form the corresponding mathematical models.

### Toggle Designs

For the sake of completeness of our description, we begin our discussion of the S toggle and A toggle designs (Fig. 2) with two classical orthogonal repressors (Table 1):

I. LacI from *E. coli* which inhibits the transcription of the second repressor gene, tetR from the tetracycline-resistance transposon Tn10;
II. TetR which represses the transcription of the first repressor gene *lacI*.

Next, the communication network among all toggles (Fig. 2) is built upon two quorum-sensing (QS) signaling molecules (Table 1):

1. *N* -(3-hydroxy-7-*cis*-tetradecenoyl)-L-homoserine lactone (3-OH-C14-HSL);
2. *N* -butanoyl-l-homoserine lactone (C4-HSL).

For the sake of brevity, the QS signaling molecules are called autoinducers G (C14-HSL) and R (C4-HSL). Note that the Gand R-signals (acylated homoserine lactones) are natural biological signals secreted by Gram-negative bacteria, including *E. coli*, as a means of coordinating cellular activity [4,8].

Finally, to drive the autoinducer concentrations, two synthases are used (Table 1):

a. CinI, the gene product of *cinI*, driving the concentration of C14-HSL;
b. RhlI, the gene product of *rhlI*, driving the concentration of C4-HSL.

Using the above molecular species, we implement and study two different toggle designs called *symmetric* (S) and *asymmetric* (A) designs, respectively, (Fig. 2):

i. In the S design, each of the two autoinducers activates symmetrically the transcription of the corresponding repressor gene through a single promoter, that is, promoter P_cin_ (P_G_) for gene *lacI* (*x*) and promoter P_rhl_ (P_R_) for gene *tetR* (*y*);
ii. In the A design, the same repressor genes (as used in the S design) antagonistically repress one another directly, while there is only one autoinducer that asymmetrically facilitates communication between all toggles.

**Figure 2.**
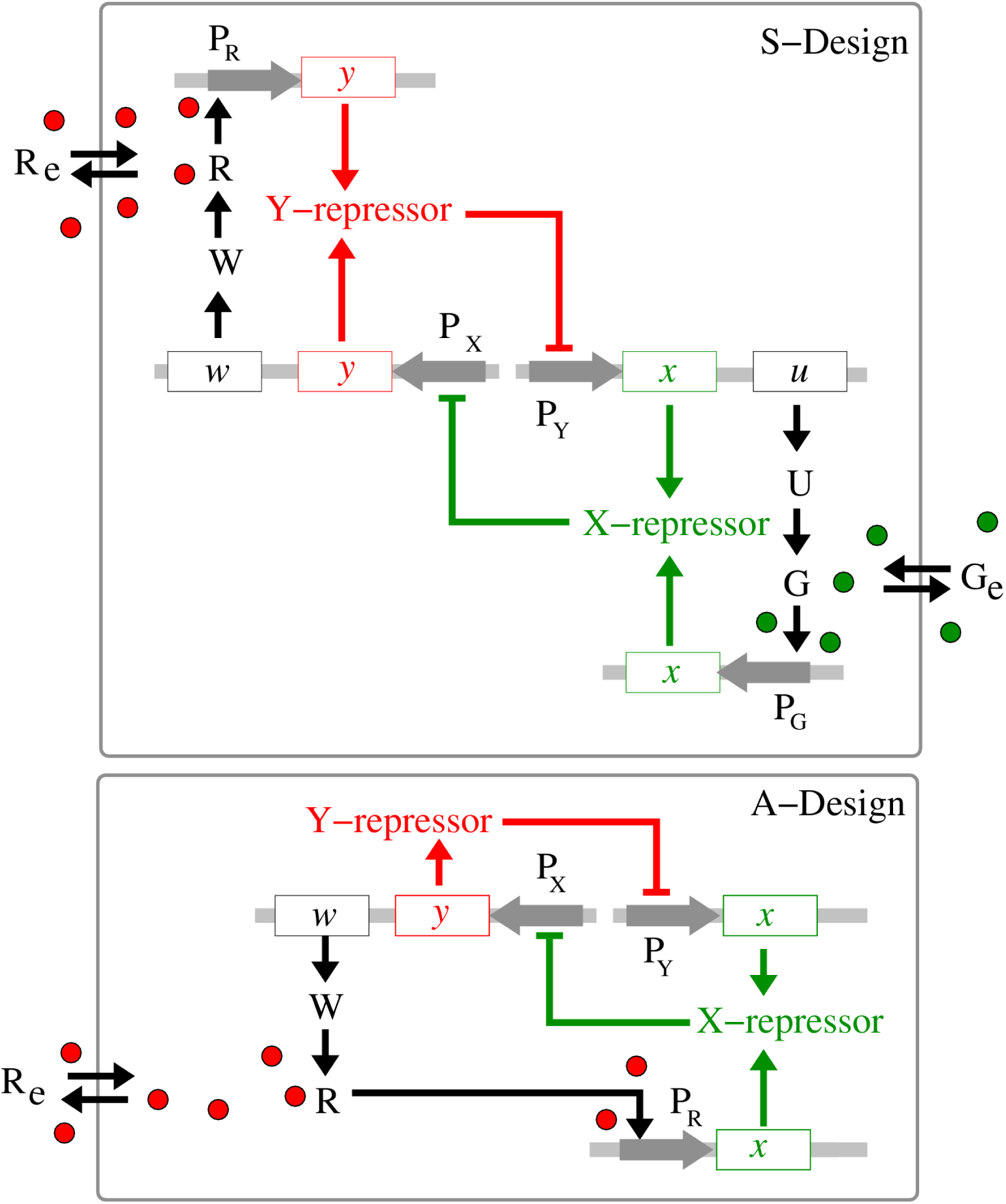
Monotone-symmetric and nonmonotone-asymmetric toggle designs. S design. (*top panel*): Activation of the expression of gene *x* (*lacI*) occurs by binding of autoinducer G (C14-HSL) to promoter PG (Pcin). Inhibition of the expression of both genes *x* (*lacI*) and *u* (*cinI*) occurs by binding of the gene product Y (TetR) of gene *y* (*tetR*) to a single promoter PY (Ptet). Symmetrically, activation of the expression of gene *y* (*tetR*) occurs by binding of autoinducer R (C4-HSL) to promoter P_R_ (P_rhl_), while inhibition of the transcription of both genes *y* (*tetR*) and *w* (*rhlI*) occurs by binding of X (LacI) to a single promoter P_X_ (P_lac_). **A design** (*bottom panel*): Activation of the expression of gene *x* (*lacI*) occurs by binding of autoinducer R (C4-HSL) to promoter P_R_ (P_rhl_). Expression of genes *y* (*tetR*) and *w* (*rhlI*) is driven by a common single promoter PX. Gene products U and W are synthases CinI and Rhil, respectively. Gray horizontal strips correspond to integration plasmids.

**Table 1.** A toggle molecular part catalog (explanations of variables are given in S Model).

| Name | variable | Function | Description | References |
| --- | --- | --- | --- | --- |
| *lacI* | — | repressor gene | lactose-inducible transcriptional repressor from *E. coli* | [1, 2, 6, 31] |
| *tetR* | — | repressor gene | from the tetracycline-resistance transposon Tn10 | [1, 2, 6, 31, 34] |
| *cinI* | — | autoinducer gene | encodes protein CinI which synthesizes C14-HSL | [8, 10, 11] |
| *rhII* | — | autoinducer gene | encodes protein RhII which synthesizes C4-HSL | [8, 9] |
| LacI | $x_i$ | lactose inhibitor | a DNA-binding protein encoded by *lacI* | [1, 2, 6, 30, 31, 33, 35] |
| TetR | $y_i$ | repressor protein | a basic element of tetracycline-controlled regulation | [1, 2, 6, 32, 34] |
| CinI |  | synthase | the gene product of gene *cinI* | [8, 10, 11] |
| RhII |  | synthase | the gene product of gene *rhII* | [8, 9] |
| C14-HSL | $g_i, g_e$ | autoinducer | *N*-(3-hydroxy-7-*cis*-tetradecenoyl)-L-Homoserine Lactone | [10] |
| C4-HSL | $r_i, r_e$ | autoinducer | *N*-butyryl-L-Homoserine Lactone | [9, 22] |

The genetic circuit topology used in the A design (Fig. 2) is taken from [13]. In order to keep making a fair comparison with the S design, we have replace the *luxA*-*luxI* system considered in [13] by the *lacI* -*tetR* system suggested in [1]. Note that both CinI and RhiI are homologous to LuxI [36].

To host the S and A toggles, we use *E. coli*, a bacterial cell which has been well-studied in a huge number of relevant experimental and modeling works [30,37–46], and which has been widely used to implement and test various synthetic circuits [1,2,4,12]. A practical modeling reason for this selection is narrowing-down our search for biologically-meaningful parameters to values known from the *E. coli* studies. However, our conclusions do not depend in any way on biological properties of the host.

As a readout of the toggle state in individual cells, we further assume that each *E. coli* cell contains a gene encoding a spectrally distinct fluorescent reporter, GFP for gene *lacI*, and RFP for gene *tetR*, driven by promoters that respond to the autoinducers C14-HSL and C4-HSL, respectively. We do not simulate the processes of bio-synthesis and degradation of the fluorescent proteins explicitly, using appropriate cascade models, for two reasons: *(i)* the “reporter” submodel does not affect the dynamics of the entire model, and *(ii)* the half-lives of the reporter proteins can be made similar to the half-lives of the repressor proteins [2].

Finally, because each toggle can either be in a state where *(a)* LacI protein is abundant, while TetR protein is scarce, or in a state where *(b)* TetR protein is abundant, while LacI protein is scarce, we call state *(a)* a green state or, simply, a G-state and state *(b)* a red state or, simply, an R-state, respectively.

### S Model

A mathematical model describing a population of identical S toggles is

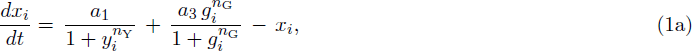

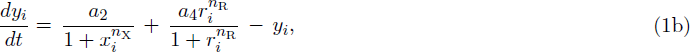

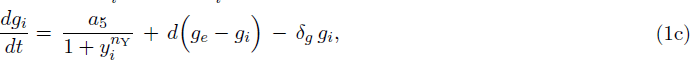

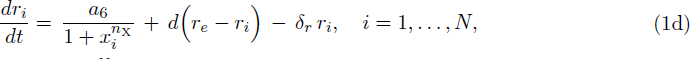

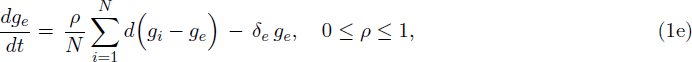

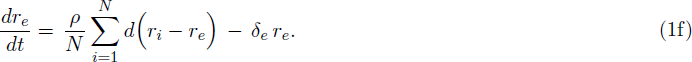

Here, all state variables and parameters are dimensionless, and are obtained from the corresponding biologically meaningful state variables and parameters describing the *lac-tet* system (Table 1) after an appropriate nondimensionalization carried out in S1 Model Derivation.

In the S model (1), *t* is dimensionless time; *x*_*i*_ and *y*_*i*_ are the dimensionless concentrations (levels) of intracellular repressor proteins LacI and TetR, respectively; *g*_*i*_ and *r*_*i*_ are the dimensionless concentrations of intracellular autoinducers C14-HSL and C4-HSL, respectively; *g*_*e*_ and *r*_*e*_ are the dimensionless concentrations of extracellular autoinducers C14-HSL and C4-HSL, respectively.

The dimensionless rate constants *a*_*i*_, *i* = 1, *…,* 6, depend on the copy numbers of the plasmids that bear the corresponding genes, see relationships (4) given in Scaling; *n*_X_, *n*_Y_, *n*_G_, and *n*_R_ are the corresponding Hill coefficients; *d* is the dimensionless diffusion coefficient; *δ_g_* and *δ_r_* are the dimensionless lumped dilution-degradation rates due to the exponential cell growth and degradation of the corresponding species.

The degradation rate constants for repressor species *x*_*i*_ and *y*_*i*_ are scaled out to unity, as it is done in [1,2, 13,14], see S2 Estimation of Parameter Values; *δ_e_* is the dilution rate due to flow in the medium; *ρ* is a population density; and *N* is the number of cells in the given population.

### A-Model

A dimensionless mathematical model describing a population of identical A toggles is

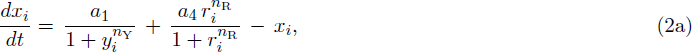

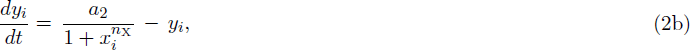

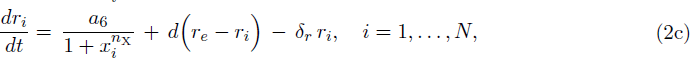

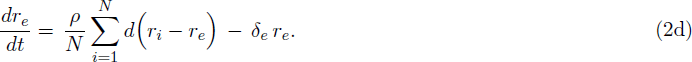

Here, all state variables and parameters are as defined for the S model (1). A similar model was developed in [13] for Toggle B2 shown in Fig. 1.

### Model Parameters

Uncertainty about the values of parameters characterizing molecular components of synthetic circuits always presents a significant difficulty in circuit design [2]. Here, we discuss reference values of dimensionless parameters obtained using an appropriate scaling procedure. We also explain how we select and interpret parameters for our bifurcation analysis.

### Reference Values of Model Parameters

Reference values of all parameters used in our modeling studies are estimated in S2 Estimation of Parameter Values, and these correspond to half-lives of all proteins in the range 4-10 min., which are close to a typical mRNA half-life in *E. coli* [2]. Also, to avoid competition for ribosomes [41], only a few plasmids bearing four promoters P_X_, P_Y_, P_G_, and P_R_ can be used, and we use 1-2 copies per cell, see S2 Estimation of Parameter Values. The E. *coli* replication period is assumed to be around 25 min.

Despite the fact that much is known about *E. coli* [37–39,43–46], it is not possible to model behavior in a quantitatively precise way, since not enough is yet known about molecular interactions between the toggle and the host cell to make such a description realistic [6]. Instead, we hope to identify classes of toggle designs and dynamic behaviors to determine which of the designs could lead to an improved self-synchronization reliability and an improved capability for self-correction of spontaneous synchronization errors, when a small fraction of cells flips to the opposite (undesirable) transcriptional signature state, see Spontaneous Synchronization Errors. We will also make some predictions that might help to facilitate engineering toggles with desired robust traits.

In our computational analysis, the following set of reference parameter values is used:

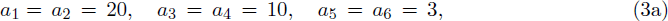

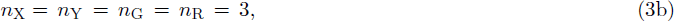

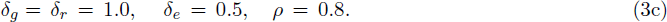

Groups of parameters with identically the same values are used to introduce the toggle mirror symmetry into the S model as discussed in Symmetry. We find that the working values of parameters estimated in (3) are within the range of equivalent parameters (rate constants, Hill coefficients, *etc*.) used earlier for genetic circuits built from similar (*e.g*., homologous) molecular entities [1,2,4,13,14,26–29,47].

Finally, we note that the above choices of the parameter values can also be viewed biologically meaningful because both the strengths of the four different promoters, P_X_, P_Y_, P_G_, and P_R_, and the degradation/dilution rates can be set accordingly by appropriate experimental interventions [1,2], reviewed in [6] and also discussed below, see Scaling and Selection and Interpretation of Bifurcation Parameters.

### Scaling

One of the goals of a model nondimensionalization and scaling is to reduce the number of (correlated) parameters by lumping original parameters into a smaller parameter set. In this case, interpretation of changes in the values of dimensionless parameters should be done carefully, as the set of non-dimensionless parameters is usually not in one-to-one correspondence with the set of original parameters. For example, mathematical models used for synthetically engineered systems often contain parameters representing multiple biological parts and, so, tuning a dimensionless parameter in the corresponding mathematical model can be implemented experimentally in a number of different ways [6].

The dimensional and dimensionless parameters used in the S and A toggle models are related to one another by the following relationships (see S1 Model Derivation):

1. For the dimensionless rate parameters, we obtain:

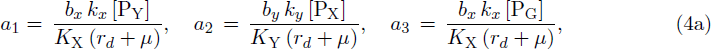

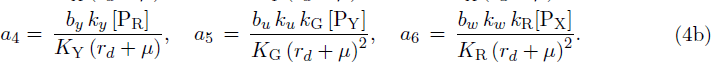

2. For the dimensionless diffusion and degradation parameters, we obtain:

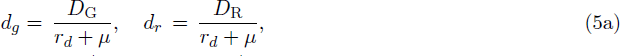

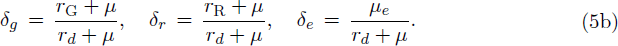

Let us briefly discuss (4) and (5). Here, the burst parameter *b*_*x*_ for the protein X or, equivalently, LacI, depends on the efficiency of translation, controlled by strength of ribosome-binding sites (RBS) [1,6], and the mRNA half-life time [48]; [P_X_] is the number of promoters per cell for gene *x*; *k*_*x*_ is an average transcription rate for gene *x* (*lacI*); *K*_X_ is the number of LacI proteins required to half-maximally repress P_lac_; *k*_G_ is the maximum production rate of C14-HSL by CinI, *D*_G_ is the export rate of C14-HSL; *μ* is the intracellular specific dilution rate due to the host cell growth, *μ* = ln 2*/T*, *T* is the division period. Parameters for other proteins and QS signaling molecules are defined similarly, see S1 Model Derivation and S2 Estimation of Parameter Values. Based on the fact that N-Acyl Homoserine Lactone Lactonase (AHL-lactonase) hydrolyzes C4-HSL effectively [49], we also assume that specific degradation rate constants for the signaling molecules, C14-HSL and C4-HSL, can be set experimentally [6], corresponding to the parameter values used in our models. We pick these specific promoters and autoinducers in order to be concrete and to justify biologically meaningful values of the model parameters. However, we wish to emphasize that our results are generic for the architectures shown in Fig. 2.

### Selection and Interpretation of Bifurcation Parameters

In our bifurcation analysis, we use almost all dimensionless parameters given in (3) as free parameters allowed to be varied to detect changes in stability of the corresponding solutions. Whenever a new bifurcation point is detected, we provide an appropriate interpretation in terms of interventions reviewed in [6], which can potentially lead to the corresponding effect.

For example, suppose that *d* is the free parameter used in bifurcation analysis. Due to the relationships (5), changes in the values of *d* may correspond to different and independent experimental interventions [6] leading to: *(a)* changes in the membrane permeability (*i.e*., *D*_G_ and *D*_R_), or *(b)* changes in the half-lives of repressor proteins, or *(c)* changes in the specific growth rate of the host cell. As such, both higher values of protein half-lives and diffusion permeability as well as lower values of the specific growth rate (longer replication periods) correspond to higher values of the parameter *d*. Recall that the value of the parameter *d* characterizes the strength of the interaction between cells in the given population, which facilitates self-synchronization [12–14,23,50].

More broadly, we can rely upon the fact that all dimensionless parameters are defined via appropriate combinations of the original dimensional parameters (4) and (5) in our interpretation of results obtained from bifurcation analysis as follows.

The values of dimensionless rate parameters (*i.e*., *a*-parameters) can be changed by decreasing or increasing translational efficiency, which depends on the nucleotide sequence of the ribosome binding sites (RBS) located within the upstream noncoding part of the mRNA [46]. The RBS is encoded by the DNA sequence immediately upstream of the start codon of the gene and is an independent regulatory element that can be manipulated experimentally [1]. The values of dimensionless rate parameters can also be changed by decreasing or increasing the lifetime values of appropriate proteins. Indeed, a carboxy-terminal tag, based on ClpX, the ATP-dependent unfoldase/translocase of ClpXP recognizing specific protein substrates bearing *ssrA* tags [51], can be inserted at the 3W end of each repressor gene [2]. Proteases in *E. coli* recognize this tag and target the attached protein for destruction. Such tags are used to reduce the half-life of the proteins from more than 60 min to around 4 min, which makes it possible and (also convenient) to set the half-life times for all toggle proteins (approximately) equal to one another and close to the half-lives of mRNAs [2,12]. To this end, both RBS and carboxy-terminal tags are the principal tools by which the parameters of an engineered gene network can be adjusted experimentally [1,2,6].

### Symmetry

In this section, we formalize the symmetry of

- a single S toggle embedded into an environment (*N* = 1),
- a population (network) of *N* -identical S toggles interacting via a common environment (*N* ≥ 2).

As is routine in physics and engineering, symmetry-based simplifications often lead to important insights into complex phenomena, and we also use symmetry to discuss bifurcations in S toggle populations. We observe that under a special condition imposed on the parameter values,

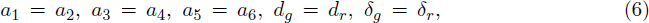

a single S toggle embedded into an external environment is described by the S model (1) with *N* = 1 which has a ℤ_2_-symmetry group generated by involution *𝒤* [52,53],

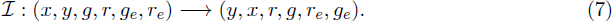

Consider the fixed-point subspace Fix (ℤ_2_) ∈ ℝ^6^ of the group ℤ_2_, see [52],

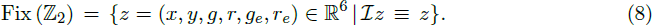

We ignore the trivial equilibria that belong to Fix (ℤ_2_), that is, equilibria of the S model (1) for which the following equalities hold, *x* = *y*, *g* = *r*, and *g*_*e*_ = *r*_*e*_, corresponding to identically the same levels of LacI and TetR, and C14-HSL and C4-HSL, respectively. Let us denote the equilibrium of the S model (1) by *z*_0_, and let us assume that *z*_0_ ∉ Fix (ℤ_2_). Now, because the S model is invariant with respect to the involution (7), and because *z*_0_ ∉ Fix (ℤ_2_), we obtain that both *z*_0_ and *𝒤 z*_0_, *𝒤 z*_0_ ≠ *z*_0_, are different equilibria of the S model (1), see [52,53]. The equilibria *z*_0_ and *𝒤 z*_0_ are called relative equilibria [54]. All bifurcations for the relative equilibria occur simultaneously at the same values of free parameters. We generalize Gand R-homogeneous populations states as relative equilibria, which means that as soon as the S toggle has a G-state, it will also have the corresponding R-state, implying bistability.

The general case of *N* ≥ 2 is slightly more complicated as a population of identical S toggles has a symmetry group obtained after combinations of permutations among all cells in the given population and the toggle involution (7), which we denote *𝒢* = ℤ_2_ *×* **S**_*N*_ for brevity. Here, **S**_*N*_ is a symmetric group of order *N* ≥ 1, and ℤ_2_ is the toggle involution (7) applied to all toggles simultaneously. For example, for *N* = 2, we will have one permutation,

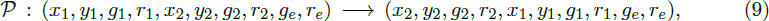

and the involution,

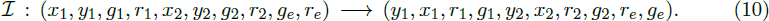

### Spontaneous Synchronization Errors

Capabilities of toggles to fail and recover from spontaneous synchronization errors can be formalized in terms of a *multistability* concept, that is, as a co-existence of bistable *homogeneous* populations and various *heterogeneous* populations (Fig. 3), also called *mixed* states, under the same conditions. Recall that mixed states are known to lead to bistable distributions [4].

Following [4], we call a population heterogeneous or, equivalently, mixed if it comprises toggles with *different* transcription signatures for the *same* genes: *(i)* the repressor gene *lacI* is active (G-state), while *tetR* is repressed, and *(ii) lacI* is repressed, while the repressor gene *tetR* is active (R-state), see Toggle Designs. In other words, a homogeneous population is fully characterized by either transcription signature *(i)* or *(ii)*, while a heterogeneous population is characterized by mixed signatures *(i)* or *(ii)* simultaneously present in the population (Fig. 3).

Different heterogeneous populations can be characterized by transcription signature “mixtures” with ratio (*p*:*q*), *p* + *q* = 1, describing the fraction of toggles in the G-state versus the fraction of toggles in the R-state within the same population. For homogeneous populations, we, therefore, have either (1:0) or (0:1) transcriptional signature fulfilled by all toggles in the population (Fig. 3).

With these concepts, we can formulate more precisely our objective: to find conditions under which heterogeneous (mixed) population equilibrium solutions can loose their stability or can even be eliminated completely.

As a proof of concept, an example of an (9:1)-heterogeneous population (Fig. 3) will be used, where the number of toggles in the first, Green-subpopulation (G) (*tetR* is suppressed) is 9 times bigger that the number of toggles in the second, Red-subpopulation (R) (*lacI* is suppressed). In this simplest case, the G-subpopulation comprises 9 cells (*p* = 0.9 or 90%-fraction of all cells), while the R-subpopulation comprises one cell (*q* = 0.1 or 10%-fraction of all cells).

**Figure 3.**
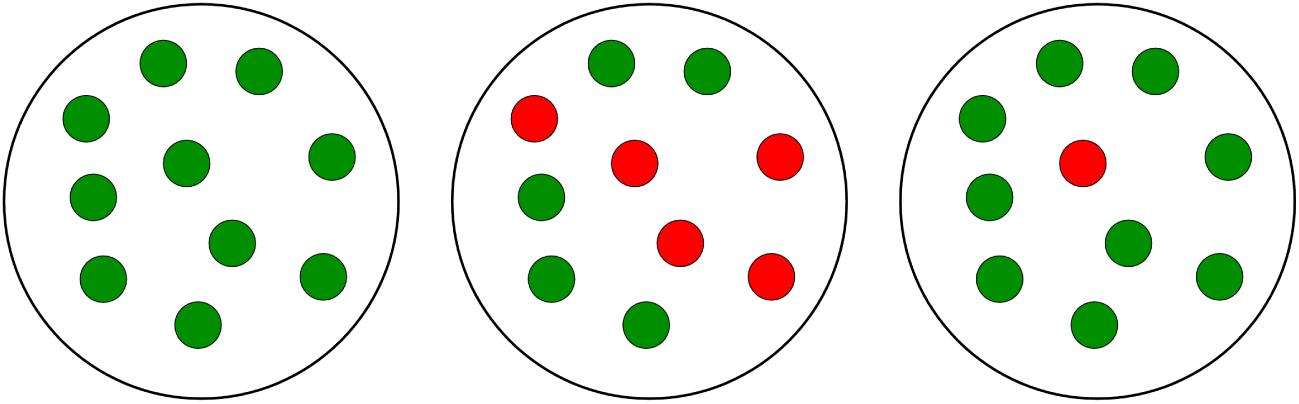
Homogeneous and heterogeneous (mixed) populations. An example of a population consisting of 10 cells is shown. The left panel demonstrates a homogeneous G-population. The center panel demonstrates a heterogeneous (1:1)-population, where the homogeneous Gand R-subpopulations have equal number of cells. The right panel demonstrates a heterogeneous (9:1)-population formed of two unequal subpopulations which represent a *spontaneous synchronization error*, when one or a few toggles spontaneously flip from green (G) to red (R) states. Therefore, our modeling goal is to find conditions under which the population can recover from the heterogeneous (right panel) case to the homogeneous (left panel) case. We will also try to eliminate cases similar to the one shown in the center panel which we, however, do not interpret as a synchronization error.

Note that our analysis of (9:1)-mixed states does not depend on the number of cells *N* in the entire population, which is usually unknown in experiments. In other words, our results hold for any integers *N*, *N*_1_, and *N*_2_, such that *N* = *N*_1_ + *N*_2_, and *N*_1_ : *N*_2_ = 9 : 1, where the fractions of cells with different transcription signatures are defined by the numbers *p* = *N*_1_*/N* and *q* = *N*_2_*/N*, respectively, see S3 Exponential Stability of Cellular Populations.

### Monotone Systems Formalism

The systems considered here are described by the evolution of states, which are time-dependent vectors *x*(*t*) = (*x*_1_(*t*), *…, x_n_*(*t*)). The components *x*_*i*_ represent concentrations of chemical species (such as proteins, mRNA, metabolites, and so forth), the dynamics of which are given by a system of ODE’s:

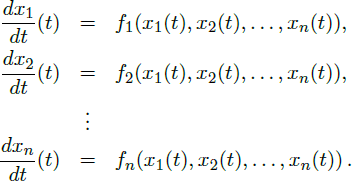

We also write simply *dx/dt* = *f* (*x*), where *f* is a differentiable vector function with components *f*_*i*_. The coordinates *x*_*i*_(*t*) are non-negative numbers. We write *φ*(*t, x*_0_) for the solution of the initial value problem 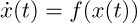 with *x*(0) = *x*_0_, or just *x*(*t*) if *x*_0_ is clear from the context, and assume that this solution *x*(*t*) exists and remains bounded for all *t ≥* 0.

#### Definition of Monotone Systems

A system is said to be *monotone* if there exists a partition of the set of indices of state variables:

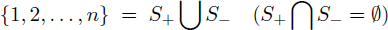

with the following properties:

(1) for each pair of indices {*i, j*} *∈ S*_+_ (*i ≠ j*) and each pair of indices {*i, j*} ∈ *S_−_* (*i ≠ j*),

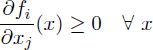

(2) and for each pair of indices {*i, j*} such that *i ∈ S*_+_ and *j ∈ S_−_* and each pair of indices {*i, j*} such that *i ∈ S_−_* and *j ∈ S*_+_,

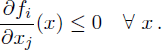

Observe that the definition does not impose any constrains on diagonal entries 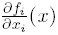. These may have arbitrary signs, even depending on *x*.

Monotone systems [55–57] were introduced by Hirsch, and constitute a class of dynamical systems for which a rich theory exists. (To be precise, we have only defined the subclass of systems that are “monotone with respect to some orthant order” but the notion of monotone dynamics can be defined with respect to more general orders.)

We assume from now on that our system satisfies the following property: for each pair of distinct nodes *i* and *j*, one of these holds:

1. 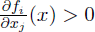 for all states *x*

2. 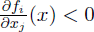 for all states *x*

3. 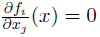 for all states *x*.

Of course, there are many models for which partial derivatives may change sign depending on the particular point *x*. With assumptions (1-3), however, the main results that we need from monotone dynamical systems theory will be particularly easy to state.

Monotone systems cannot admit any stable oscillations [16,58,59]. Under a stronger property, described next, only convergence to steady states is generically possible.

#### Strong Monotonicity

The directed *species influence graph G* associated to a system with *n* state variables is defined as follows. The graph *G* has *n* nodes (or “vertices”), which we denote by *v*_1_, *…, v_n_*, one node for each species. If

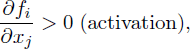

we introduce an edge labeled “1” from *v*_*j*_ into *v*_*i*_. If, instead,

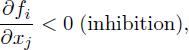

we introduce an edge labeled “*-*1” (or just “*-*”) from *v*_*j*_ into *v*_*i*_. Finally, no edge is drawn from node *v*_*j*_ into node *v*_*i*_ if the partial derivative 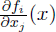 vanishes identically (no direct effect of the *j*th species upon the *i*th species). An alternative is to write a normal arrow “*→*” or a blunted arrow “┤” (or an arrow labeled “*-*”) respectively for the first two cases. The graph *G* is an example of a *signed graph* [60], meaning that its edges are labeled by signs.

No self-edges (edges from a node *v*_*i*_ to itself) are included in the graph *G*, whatever the sign of the diagonal entry *∂f_i_/∂x_i_* of the Jacobian. The sign of this derivative may be positive, negative, or even be state-dependent. Results will not depend on signs of diagonals of the Jacobian of *f*.

The graph *G* is said to be *strongly connected* if, given an arbitrary pair of different indices {*i, j*}, there is a some, possibly indirect, effect of *i* on *j*. Formally, we ask that there is a sequence of indices *i* = *k*_0_, *k*_1_, *…, k_r_* = *j* such that

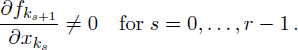

A system is said to be *strongly monotone* if it is monotone and, in addition, its species influence graph *G* is strongly connected. (As with the definition of monotonicity, one can extend strong monotonicity to far more general classes of systems, but we use a more restrictive notion that makes results less technical to state.) Even when there are multiple steady-states, the Hirsch Generic Convergence Theorem [55–57,59] is a fundamental result.

#### Hirsch’s Theorem

Even though they may have arbitrarily large dimensionality, monotone systems behave in many ways like one-dimensional systems: Hirsch’s Theorem asserts that generic bounded solutions of strongly monotone differential equation systems must converge to the set of (stable) steady states. “Generic” means here “for every solution except for a measure-zero set of initial conditions.” In particular, no nontrivial attractors arise. The genericity qualifier is needed in order to exclude the unstable manifolds of saddles as well as behavior on lower-dimensional sets [15].

### Order Preservation by Flows of Monotone Systems

We defined monotonicity using graph partitions because this is the easiest way to present the concept. However, the usual definition found in textbooks is not phrased in that form. We explain here how our definition is equivalent to the usual one as well as to another property. For further remarks on these equivalences, see [20]. A signed graph (such as the species influence graph obtained by looking at signs of Jacobain entries) *G* is said to be *balanced* (see Harary [61]) if every undirected closed loop in the graph *G* has a net positive sign, that is to say, an even number, possibly zero, of negative arrows. Equivalently, any two (undirected) paths between two nodes must have the same net sign. By undirected loops or paths, we mean that one is allowed to transverse an edge either forward or backward. A *spin assignment* Σ for the graph *G* is an assignment, to each node *v*_*i*_, of a number *σ_i_* equal to “+1” or “*−*1” (a “spin,” to borrow from statistical mechanics terminology). If there is an edge from node *v*_*j*_ to node *v*_*i*_, with label *J*_*ij*_ ∈ {*±*1}, we say that this edge is *consistent with the spin assignment* Σ provided that:

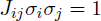

which is the same as saying that *J*_*ij*_ = *σ_i_σ_j_*, or that *σ_i_* = *J*_*ij*_σ_*j*_. An equivalent formalism is that in which edges are labeled by “0” or “1,” instead of 1 and *-*1 respectively, and edge labels *J*_*ij*_ belong to the set {0, 1}, in which case consistency is the property that *J*_*ij*_ ⊕ σ_*i*_ ⊕ σ_*j*_ = 0 (sum modulo two). One says that Σ *is a consistent spin assignment for the graph G* (or simply that *G* is consistent) if every edge of *G* is consistent with Σ. In other words, for any pair of vertices *v*_*i*_ and *v*_*j*_, if there is a positive edge from node *v*_*j*_ to node *v*_*i*_, then *v*_*j*_ and *v*_*i*_ must have the same spin, and if there is a negative edge connecting *v*_*j*_ to *v*_*i*_, then *v*_*j*_ and *v*_*i*_ must have opposite spins. (If there is no edge from *v*_*j*_ to *v*_*i*_, this requirement imposes no restriction on their spins.) It is easy to see that if there is a consistent spin assignment for *G*, then the graph is balanced. Conversely, if *G* is balanced then there is a consistent spin assignment for *G*: to see this, simply label one node arbitrarily, and follow paths to label other nodes consistently. (If the graph is not connected, repeat the procedure in each connected component.)

For any spin assignment Σ, let *A*_1_ be the subset of nodes labeled +1, and let *A*_−1_ be the subset of nodes labeled *-*1. The set of all nodes is partitioned into *A*_1_ and *A*_−1_. Conversely, any partition of the set of nodes into two subsets can be thought of as a spin assignment. With this interpretation, a consistent spin assignment is the same as a partition of the node set into two subsets *A*_1_ and *A*_−1_ in such a manner that all edges between elements of *A*_1_ are positive, all edges between elements of *A*_−1_ are positive, and all edges between a node in *A*_1_ and a node in *A*_−1_ are negative. In summary, our definition of monotonicity, given in terms of partitions of state variables, amounts to the same as the requirement that there exist at least one consistent spin assignment for its associated graph *G*, or equivalently, that its graph *G* is balanced.

Supposing that a system is monotone, with a consistent spin assignment Σ = {*σ_i_, i* = 1, *…, n*}, we introduce following the relation among vectors 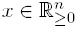:

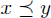

means that

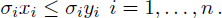

This is a componentwise inequality that requires *x*_*i*_ ≤ *y*_*i*_ if node *i* has been assigned a positive spin, and *x*_*i*_ ≥ *y*_*i*_ if instead node *i* has been assigned a negative spin. Let *y*(*t*) and *z*(*t*) be any two solutions of the system *dx/dt* = *f* (*x*), and suppose that *σ_i_y_i_*(0) *≤ σ_i_z_i_*(0) for each *i* = 1, *…, n*. Then, *Kamke’s Theorem* states that *σ_i_y_i_*(*t*) *≤ σ_i_z_i_*(*t*) for all *t ≥* 0 and coordinate *i* = 1, *…, n*. This is the usual definition of monotonicity: if states start at time zero in a certain order, then they must remain forever in the same order. Conversely, a flow that preserves an order of this type must be monotone in the sense that we have defined the concept. See the textbook [57] for a proof, and [18] for extensions with systems with external inputs.

## Results and Discussion

To carry out computational bifurcation analysis, MatCont [62,63] has been used. A technical description of bifurcation points can be found in [53,62–64].

### Application of Monotone Systems Theory to the S design

To apply monotone systems theory to the S toggle model (1), we first rewrite the model in the following convenient general form with 4 *N* + 2 variables:

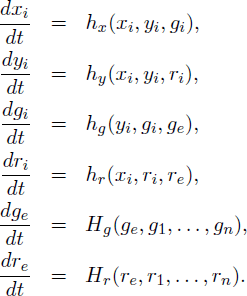

Here, *i* = 1, *…, n*, all the functions in the right-hand side are differentiable, and the following signs hold for partial derivatives, everywhere in the state space:

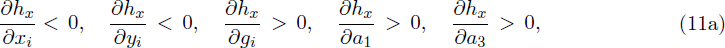

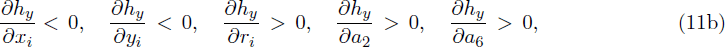

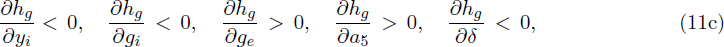

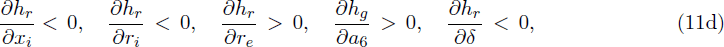

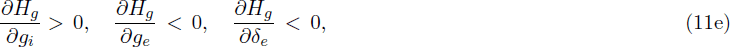

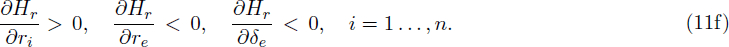

Next we observe that the S system is monotone, because we may partition its state variables as follows. One set consists of

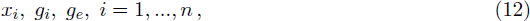

and another set consists of

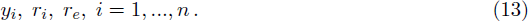

Moreover, the corresponding graph is strongly connected, as we have the following paths, for each two indices *i, j*:

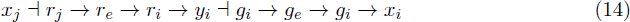

which shows that one can reach any node from any other node by means of a directed path. Thus, the S model (1) is strongly monotone. We conclude as follows.

#### Theorem 1.

*Typical solutions of the S model* (1) *converge to steady states*.

This fundamental result is robust to parameters as well as to the functional form of the equations. It insures that our proposed deign has theoretically guaranteed global stability properties. No stable oscillations [13] can exist, nor can other (for, example, “chaotic” [23]) solution regimes arise. In addition to these global properties, it is also possible to use the theory of monotone systems in order to make qualitative predictions about bifurcation diagrams as discussed in the next section.

The monotonicity property of the S system has important consequences regarding its transient as well as asymptotic behavior. We discuss in an appendix how Kamke’s Theorem characterizes order relations for monotone systems. We explain now what these mean, explicitly, for the S system. Let *z*_*i*_(*t*) characterize the state of the *i*-th S toggle at time *t ≥* 0, that is, *z*_*i*_(*t*) = (*x*_*i*_(*t*), *y*_*i*_(*t*), *g*_*i*_(*t*), *r*_*i*_(*t*)), *i* = 1, *…, N*. Let *Z*(*t*) characterize the state of the population of cells, *Z*(*t*) = (*z*_1_(*t*), *…, z_N_* (*t*), *g*_*e*_(*t*), *r*_*e*_(*t*)). Suppose that we have two initial sets, *Z*(0) and 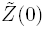, of values for the various expression levels of the repressor proteins, LacI and TerR, and we consider the behavior of *Z*(*t*) and 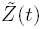 for *t* > 0.

Now suppose that we wish to understand what is the effect of a perturbation in one of the components of the initial state *z*_*i*_(0) for S toggle *i* with some fixed *i*, 1 *≤ i ≤ N*. (A similar argument can be applied to perturbations in other components of the initial state, or even simultaneous perturbations in all the components.) Suppose, for example, that we are interested in understanding the behavior starting from a state in which 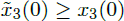 in the 3rd toggle *z*_3_. This gives rise to a new population-wide solution 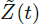, and we use a tilde to denote its coordinates, that is, 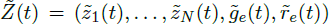, where 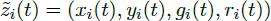, *i* = 1, *…, N*. Then, using the information provided by the partition shown in (12) and (13), we can predict that, for all 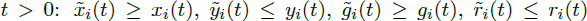, 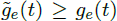 and 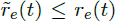 for all *i* = 1, *…, N*. As we will see shortly below, a similar conclusion can also be made with respect to perturbations in parameters, not merely initial states.

### Monotone Parametric Dependencies in the S design

As a first step, we can include the nine parameters, *a*_*i*_ (*i* = 1, *… ,* 6), *δ_g_*, *δ_r_*, and *δ_e_*, as constant state variables by formally adding the corresponding equations *da*_*i*_/*dt* = 0 (*i* = 1, *… ,* 6), and *dδ_g_/dt* = *δ_r_/dt* = *dδ_e_/dt* = 0 to the S-model (1). The extended S-model is a monotone system. Dependencies between the S-model state variables and parameters (11) as shown in Fig. 4.

The extended S system has no strong monotonicity property, because the nodes corresponding to the parameters cannot be reached from other nodes, as the parametric extension violates the strong connectivity relationships (14). However, this is not of any consequence, as the global stability properties of the S system are determined by constant values of the parameters. We only introduced the extended system in the context of bifurcation analysis.

One might add additional constant variables to represent other parameters, such as the *d*’s. These other parameters do not lead to monotonicity, and this lack of monotonicity will have important consequences in bifurcation analysis, as we discuss later.

The monotonicity of the extended system implies that stable loci in bifurcation diagrams depend monotonically on parameter variations. They will increase when the parameter being varied belongs to the component as the variable being analyzed, and will decrease if they are in different components. This property is a consequence of the general order preserving properties of monotone systems, as we explain now.

**Figure 4.**
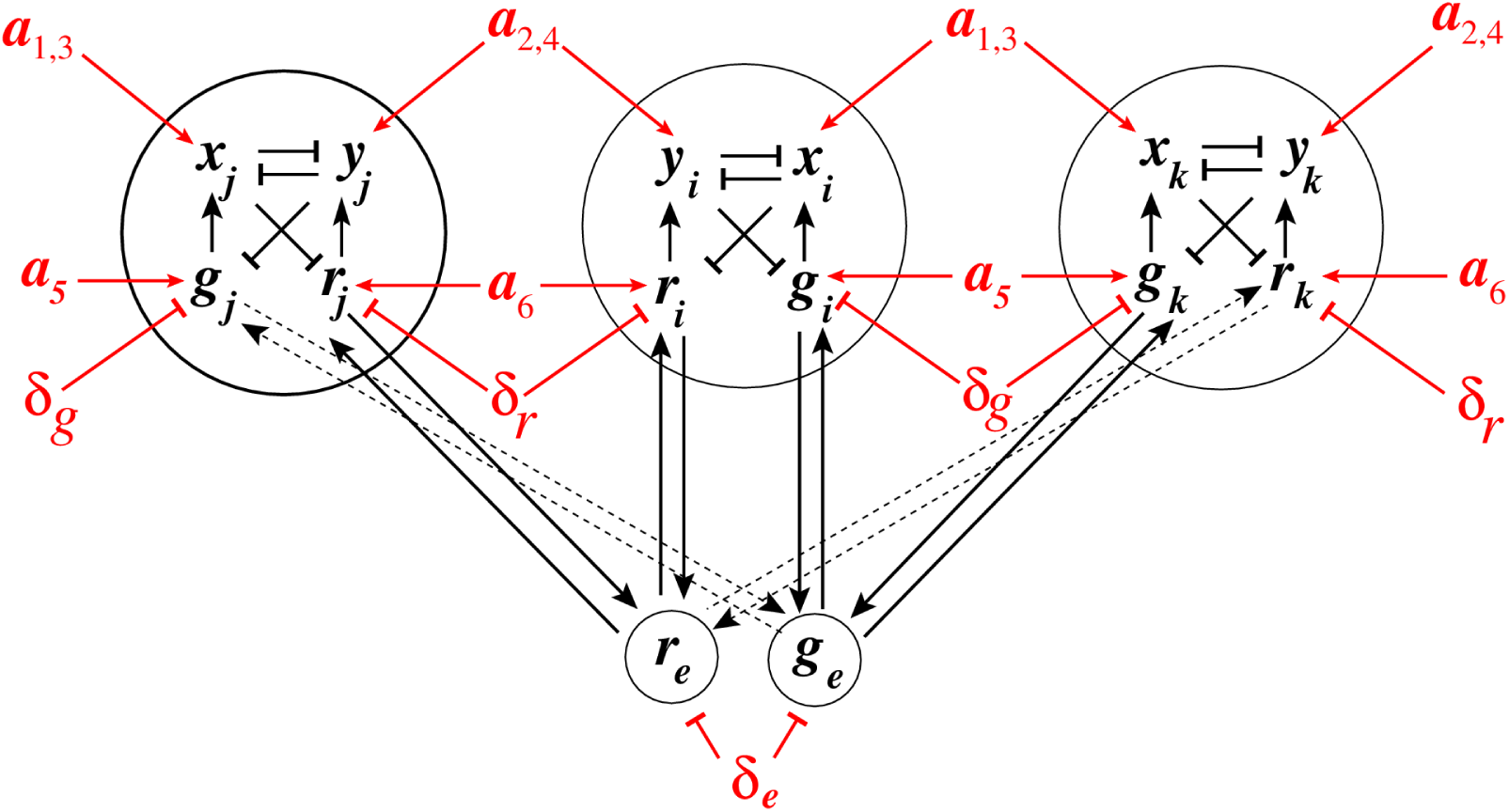
An example of three identical S toggles interacting via common autoinducers. Solid arrows and lines highlighted in red correspond to monotone parameter dependencies.

Suppose that 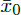 is a steady state corresponding to a real parameter value *p*_0_, that is to say, 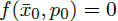. Suppose that we now consider a real parameter *p*_1_ that is very close to *p*_0_ and larger than *p*_0_, *p*_1_ *> p*_0_. Suppose in addition that 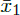 is a steady state for the parameter value 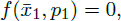, and that 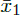 is stable. Now pick the solution *x*_1_(*t*) of 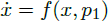 that has initial condition 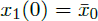. Suppose that the extended system 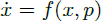 and 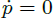 is monotone. Now, we may consider the following two initial states for the extended system: 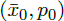 and 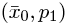. Since the second state is larger (in the sense of Kamke’s Theorem as earlier explained) in the monotone order, it follows that the solutions satisfy 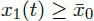 for all *t* > 0, and therefore, taking limits, we conclude that 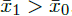, as desired.

Using Fig. 4 in conjunction with the dimension analysis in terms of the relationships (4) and (5), certain qualitative predictions can be made about the parametric dependencies based on monotone systems theory. To benchmark the approach, we have selected, as an example, a subset of dependencies shown in Fig. 4, presented in Fig. 5.

To preserve the S toggle symmetry, we change the values of the corresponding parameters for all toggles simultaneously. Using the S-model (1) with the values of fixed parameters given in (3), we find that Fig. 4 predicts monotonically increasing dependencies of the *x*-state variable (describing levels of LacI) on the values of the dimensionless parameters *a*_1_ and *a*_3_, see Fig. 5 (A) and (B), respectively.

The loss of stability and disappearance of the mixed states shown in Fig. 5 (C) as *a*_5_ increases can be interpreted intuitively by the fact that an increase in *a*_5_ leads to an increase in the intracellular levels of the corresponding QS signaling molecules, which, in turn, lead to an increase of extracellular levels of the QS molecules via diffusion, thereby facilitating self-synchronization of the given population of all toggles under conditions corresponding to a stronger interaction among all toggles. In particular, the strong interaction and coupling condition eliminates spontaneous synchronization errors in terms of suppressing the emergence of undesired (9:1)-mixed states.

This result is similar to a well-known fact for oscillators coupled via a common medium that a transition from an unsynchronized to a synchronized regime emerges as the strength of coupling increases [12–14,23,50]. Indeed, many microbial species accomplish this via quorum sensing, which entails the secretion and detection of diffusible molecules (autoinducers), whose concentration serves as a proxy for population density [7].

**Figure 5.**
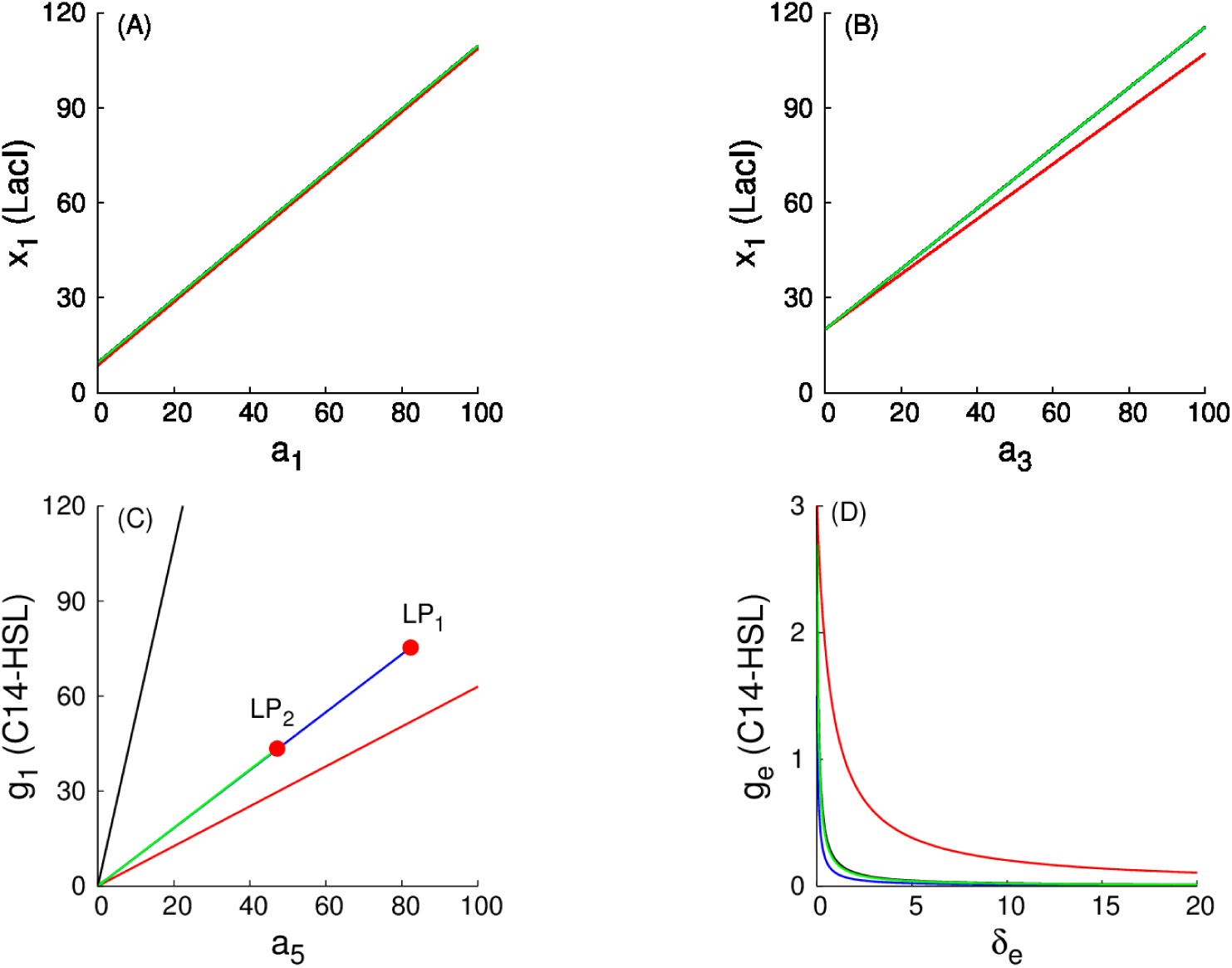
Examples of monotone parametric dependencies. Three different stable populations are chosen for the analysis of monotone parametric dependencies: (1) an G-homogeneous population; (2) an (1:1)-mixed population; here, levels of LacI and C14-HSL from one subpopulation (within which LacI is over-expressed) are shown; and (3) a (9:1)-mixed population (a spontaneous synchronization error); here, again, levels of LacI and C14-HSL from the largest subpopulation (within which LacI is over-expressed) are shown. The reference set of fixed parameter values is given by (3). The values of free parameters (*a*_1_, *a*_3_, *a*_5_, and *δe*) are unique for the entire population of identical cells and are changed simultaneously for all toggles (subpopulations). A color-coding schema for all curves is described at the bottom of the caption. **Panel (A)** shows the monotone dependence of the values in the state variable *x*_1_ (LacI) on the parameter *a*_1_. **Panel (B)** shows the dependence of *x*_1_ (LacI) on *a*_3_. **Panel (C)** shows the dependence of *g*_1_ (intracellular C14-HSL) on *a*_5_; and panel (D) shows the dependence of *ge* (extracellular C14-HSL) on *δe*. All detailed explanations, including biological interpretations, are given in the main text. Because the stable mixed populations do not exist for large values of the parameter *d* (see A (9:1)-Mixed Population Consisting of S-Toggles), we use both *d* = 0.1 (weak coupling) for all populations and, additionally, we use *d* = 10 (strong coupling) for the G-homogeneous population only. To distinguish between all the cases considered, we follow a color coding schema described below. However, due to mathematical properties of the models, projections of the corresponding plots on the 2D-planes often overlap, mixing different colors, which should not lead to any difficulty in recognizing similar monotone (“overlapping”) dependencies. **The color-coding schema for all curves includes:** *(i)* black plots are used for G-homogeneous solutions at *d* = 0.1; *(ii)* red plots are used for G-homogeneous solutions at *d* = 10; *(iii)* blue plots are used for (1:1)-mixed states at *d* = 0.1; and (*iv*) green plots are used for (9:1)-mixed states at *d* = 0.1. Red filled circles in panel (C), labeled with LP_1_ and LP_2_, correspond to Limit Point (LP) (or, equivalently, Saddle-Node) bifurcation points [62]. Here, the blue curve connecting the origin (0, 0) and the LP_1_-point corresponds to the stable branch of the (1:1)-mixed state. The green curve connecting the origin (0, 0) with the LP_2_-point corresponds to the stable branch of the (9:1)-mixed state. Because the green curve was plotted after plotting the blue curve, a part of the blue curve is hidden beneath the green curve.

Using the expression for the dimensionless parameter *a*_5_ given in (4), see Scaling, we can conclude that the increase in the values of the parameter *a*_5_ leading to the bifurcation point LP_2_ (Fig. 5) can be achieved by the following experimental interventions:

- stabilization of cell division with lower values of the specific growth rate *μ* (or, equivalently, higher division periods *T*);
- stabilization of relevant proteins, using lower values of *r*_*d*_ (or, equivalently, higher half-lives);
- an increase in the maximum production rate (*k*_G_) of C14-HSL by enzyme CinI, see S2 Estimation of Parameter Values;
- an increase in the sensitivity (*K*_G_) of promoter P_cin_ with respect to the number of molecules C14-HSL to half-activate P_cin_, see Table S2.2 given in S2 Estimation of Parameter Values.

We have used bifurcation analysis with respect to changes in the values of the parameter *a*_5_ as a way to illustrate predictions from monotone systems theory, and in the process we obtained conclusions regarding improvements of S toggle self-synchronization properties by eliminating the (9:1)-mixed state. To this end, we note that there is no need to further increase values of *a*_5_ to move the system to the bifurcation point LP_1_ at which the (1:1)-mixed state loses it stability and disappears, because we do not interpret the (1:1)-mixed state as a spontaneous synchronization error, see Spontaneous Synchronization Errors.

We next consider parametric dependencies with respect to changes in the parameters *δ_g_* and *δ_r_* (Fig. 6). As predicted by the diagram shown in Fig. 4, an increase in the values of *δ_g_* should lead to suppressed levels in *x*_1_ (LacI) as well as to elevated levels in *y*_1_ (TetR). This is illustrated in Fig. 6. The almost constant dependencies in Fig. 6 (C) and (D) can be explained by suppressed levels of TetR and C4-HSL in the G-population.

Analogously (Fig. 6), an increase in the values of *δ_g_* should also lead to suppressed levels in *g*_1_ (C14-HSL) as well as to elevated levels in *r*_1_ (C4-HSL), while an increase in the values of *δ_r_* should lead to elevated levels in *g*_1_ (C14-HSL) and, simultaneously, to decreased levels in *r*_1_ (C4-HSL). This is illustrated in Fig. 7. Constant dependencies in Fig. 7 (C) can be explained by suppressed levels of TetR and C4-HSL in the G-population.

We then repeat the analysis of the same parametric dependencies for a (1:1)-mixed state, illustrated in Fig. 8 and Fig. 9. Like in the previous case, we observe that all dependencies are in line with the predictions suggested by Fig. 4.

Additionally, the LP-bifurcation point (Fig. 8) can be interpreted as follows. Decreasing values of both parameters *δ_g_* and *δ_r_* leads to an increase in the intracellular and extracellular levels of the corresponding QS signaling molecules, which, in turn, leads to stronger interactions among all toggles. Indeed, it follows from (5) (see Scaling) that the described changes in the values of dimensionless parameters *δ_g_* and *δ_r_* can be achieved by increasing half-lives of the corresponding QS signaling molecules.

To this end and similarly to the interpretation provided earlier, as the values of the parameters *δ_g_* and *δ_r_* decrease, the (1:1)-mixed state loses its stability and disappear via an LP-bifurcation (Fig. 8), the effect which is similar to the well-known fact that oscillators coupled via common medium synchronize as the strength of coupling increases [12,23,50].

**Figure 6.**
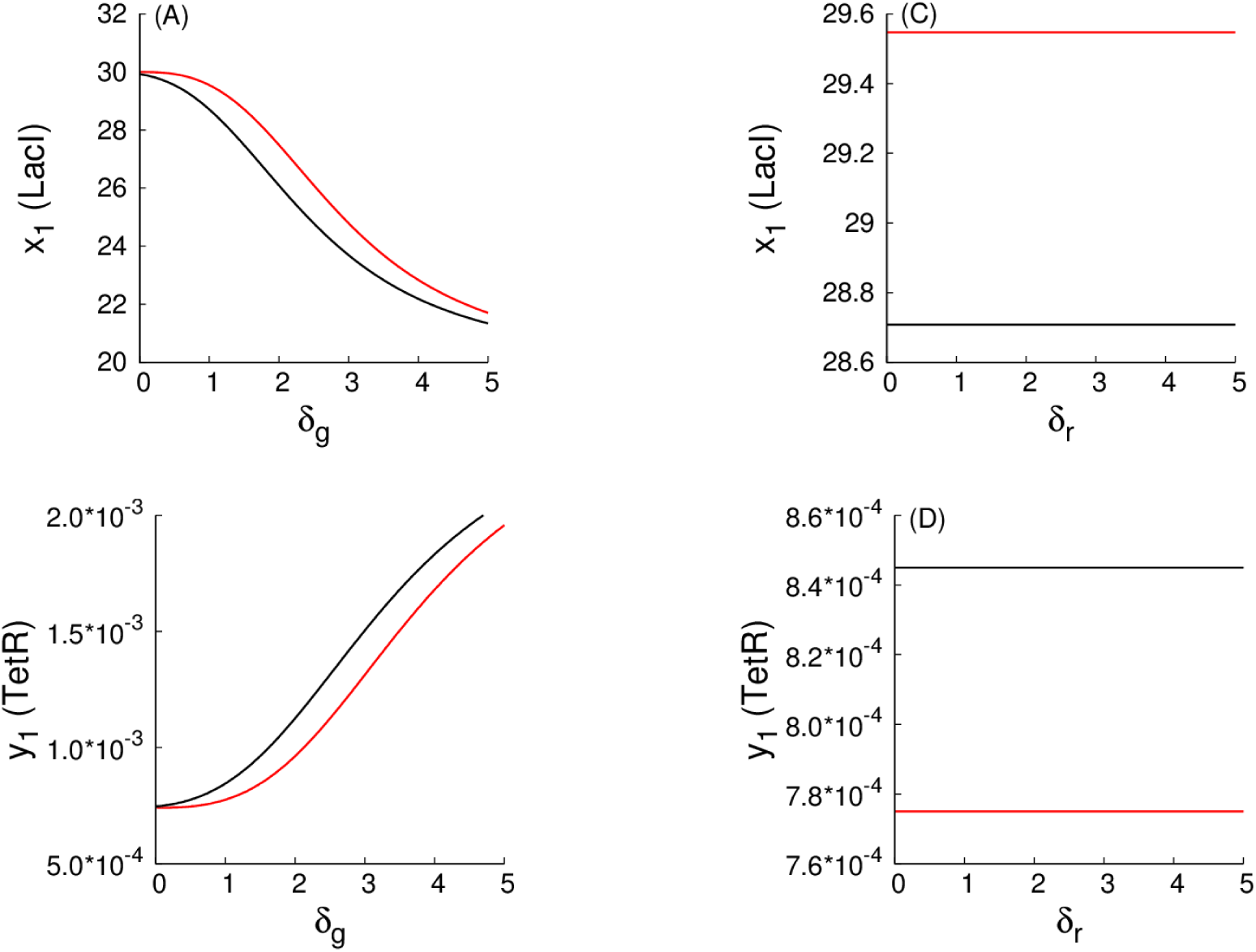
Examples of monotone parametric dependencies for the repressor-protein levels in the G-homogeneous state. Panels (A) and (B) correspond to *δ_g_*, while panels (C) and (D) correspond to *δ_r_*. Red solid curves correspond to a weak coupling among all toggles (*d* = 0.1), while black solid curves correspond to a strong coupling among all toggles (*d* = 10). The values of all other fixed parameters are given in (3), see Reference Values of Model Parameters.

**Figure 7.**
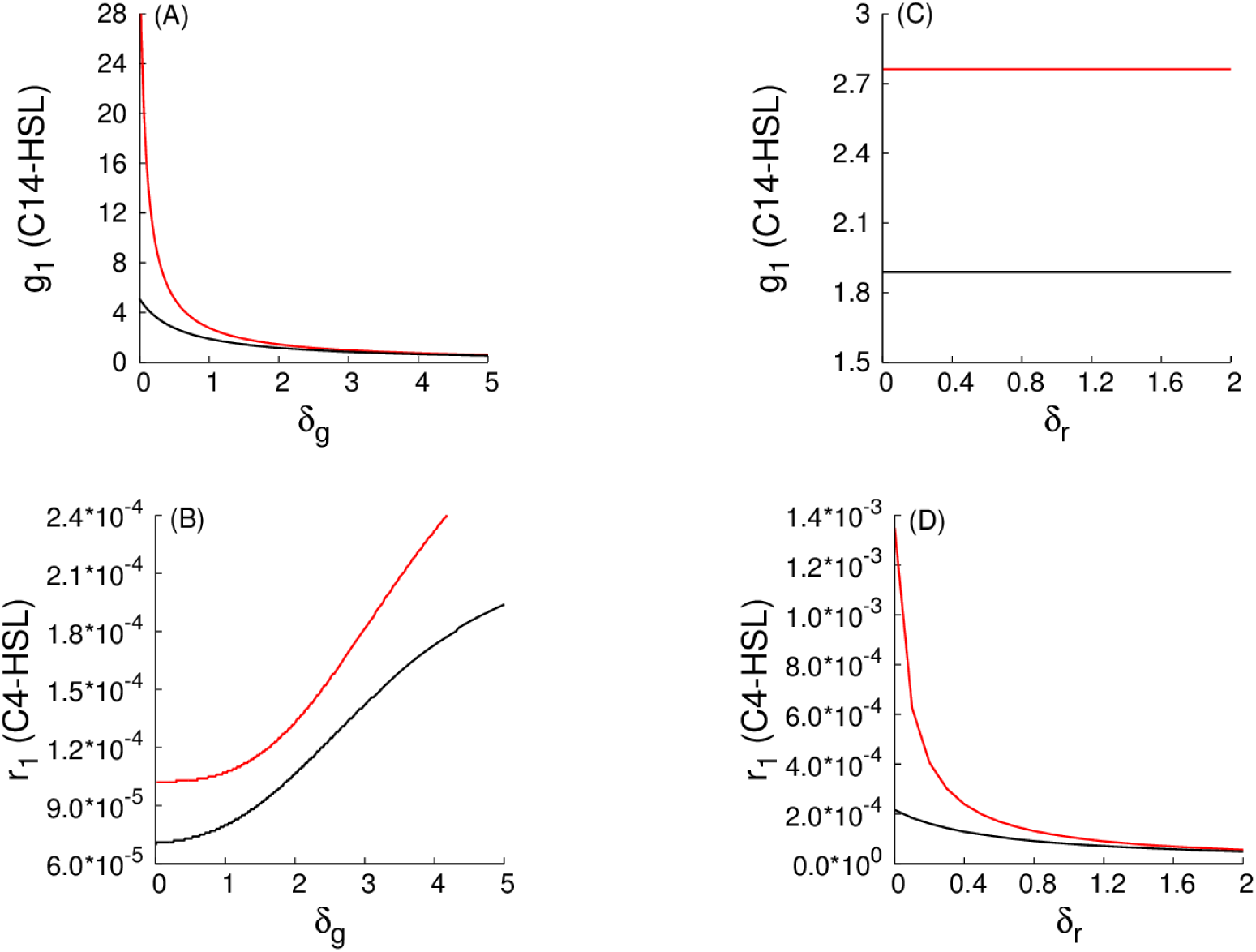
Examples of monotone parametric dependencies for the signaling species levels in the G-homogeneous state. Panels (A) and (B) correspond to *δ_g_*, while panels (C) and (D) correspond to *δ_r_*. Red solid curves correspond to a weak coupling among all toggles (*d* = 0.1), while black solid curves correspond to a strong coupling among all toggles (*d* = 10). The values of all other fixed parameters are given in (3), see Reference Values of Model Parameters.

**Figure 8.**
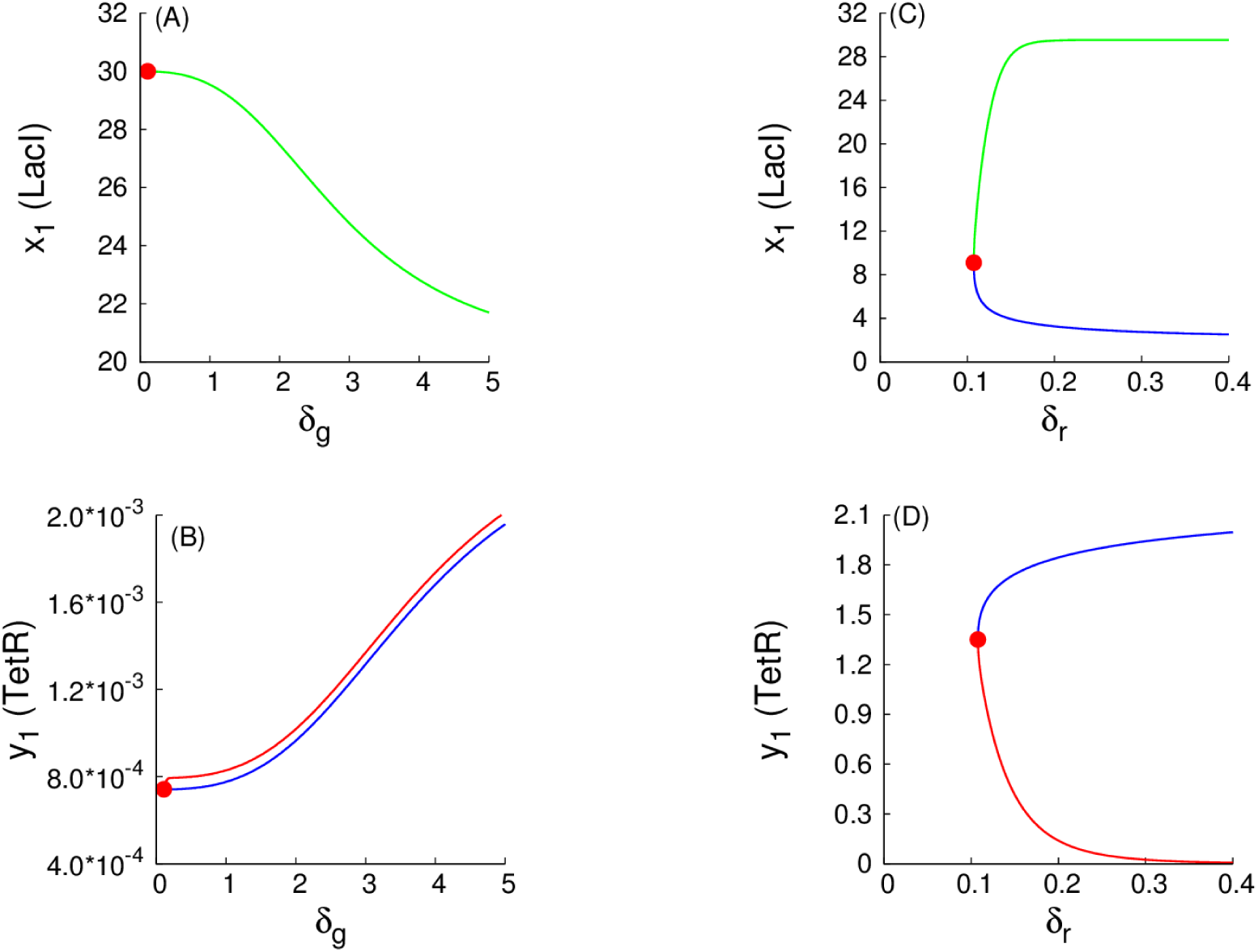
Examples of monotone parametric dependencies for the repressor-protein levels in the (1:1)-mixed state. As an example, the dependencies for the G-subpopulation are shown only, within which LacI is activated, while TetR is repressed. Panels (A) and (B) correspond to *δ_g_*, while panels (C) and (D) correspond to *δ_r_*. Green and red solid curves correspond to stable branches of (1:1)-equilibrium solutions, while all blue curves correspond to unstable branches of the solutions. Red filled circles correspond to an LP-bifurcation point. In panel (A), projections of stable and unstable branches coincide and, so, only the stable branch is shown.

**Figure 9.**
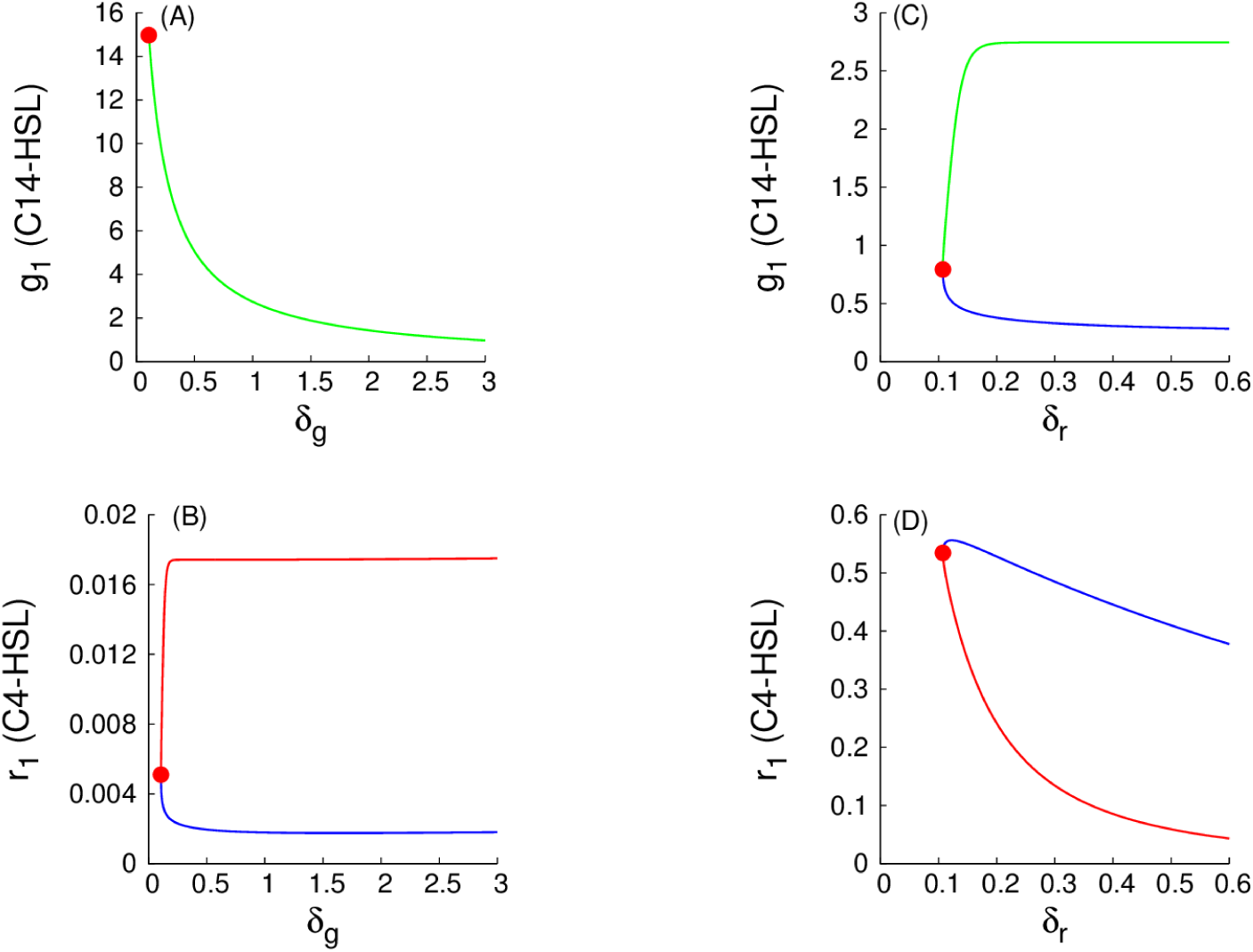
Examples of monotone parametric dependencies for the signaling species levels in the (1:1)-mixed state. Panels (A) and (B) correspond to *δ_g_*, while panels (C) and (D) correspond to *δ_r_*. All explanations are as in Fig. 8.

We note that the parametric dependencies for unstable solutions are not described by Fig. 4. To explain this observation, we recall that our proof of monotone dependence on parameters applies to stable solutions only, see above.

The monotone parametric dependencies for a (9:1)-mixed state corresponding to a spontaneous synchronization error are illustrated in Fig. 10 for a large G-subpopulation comprising 90% of all cells, and in Fig. 11 for a small R-subpopulation comprising 10% of all cells in the given (9:1)-mixed state.

We observe that LP-bifurcation points are present in both panels (A) and (B), and are absent from both panels (C) and (D) in Fig. 10 and Fig. 11. To explain this observation we have to recall the difference between parameters *δ_g_* and *δ_r_*. As discussed earlier, a decrease in the values of *δ_g_* can be interpreted in terms of the improved communication between the toggles within the large subpopulation, while a decrease in the values of *δ_r_* can be interpreted in terms of the improved communication between the toggles within the small subpopulation. We can thus conclude that by increasing the strength of interactions between the toggles from the large subpopulation, the spontaneous error can be eliminated, corresponding to the existence of the LP-points in panels (A) and (B) of Fig. 10 and Fig. 11. At the same time, increasing the strength of interactions between the toggles from the small population, the corresponding spontaneous error cannot be eliminated.

**Figure 10.**
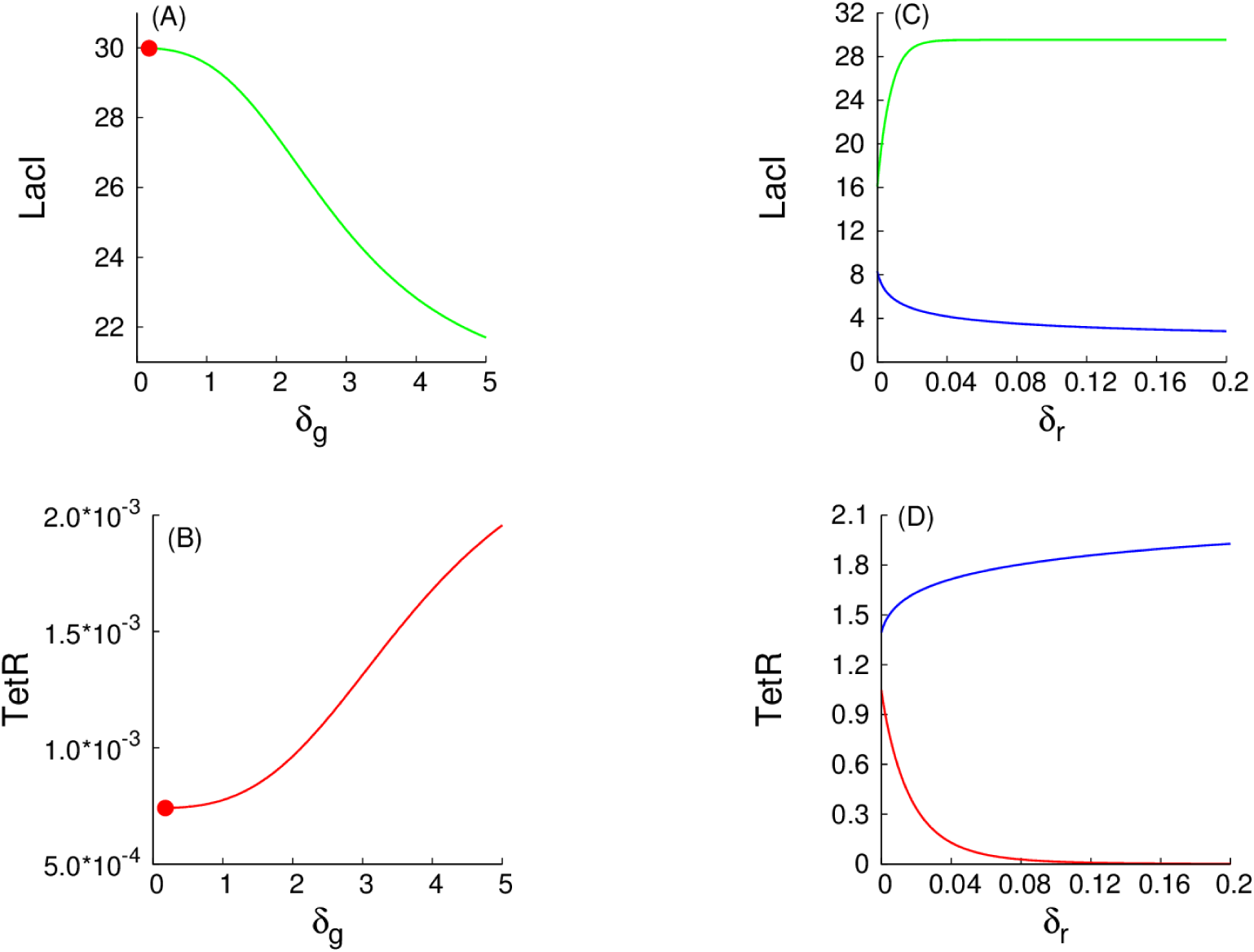
Examples of monotone parametric dependencies for the repressor-protein levels in the (9:1)-mixed state (a 90% large G-subpopulation.) The curves for the large G-subpopulation are shown in panels (A) (D), within which TetR is activated, while LacI is repressed. Panels (A) and (B) correspond to *δ_g_*. Panels (C) and (D) correspond to *δ_r_*. Green and red solid curves correspond to stable solution branches, while all blue curves correspond to unstable solution branches. Red filled circles correspond to an LP-bifurcation point. In panels (A) and (B), projections of stable and unstable solution branches coincide and, so, only the stable solution branches are shown.

**Figure 11.**
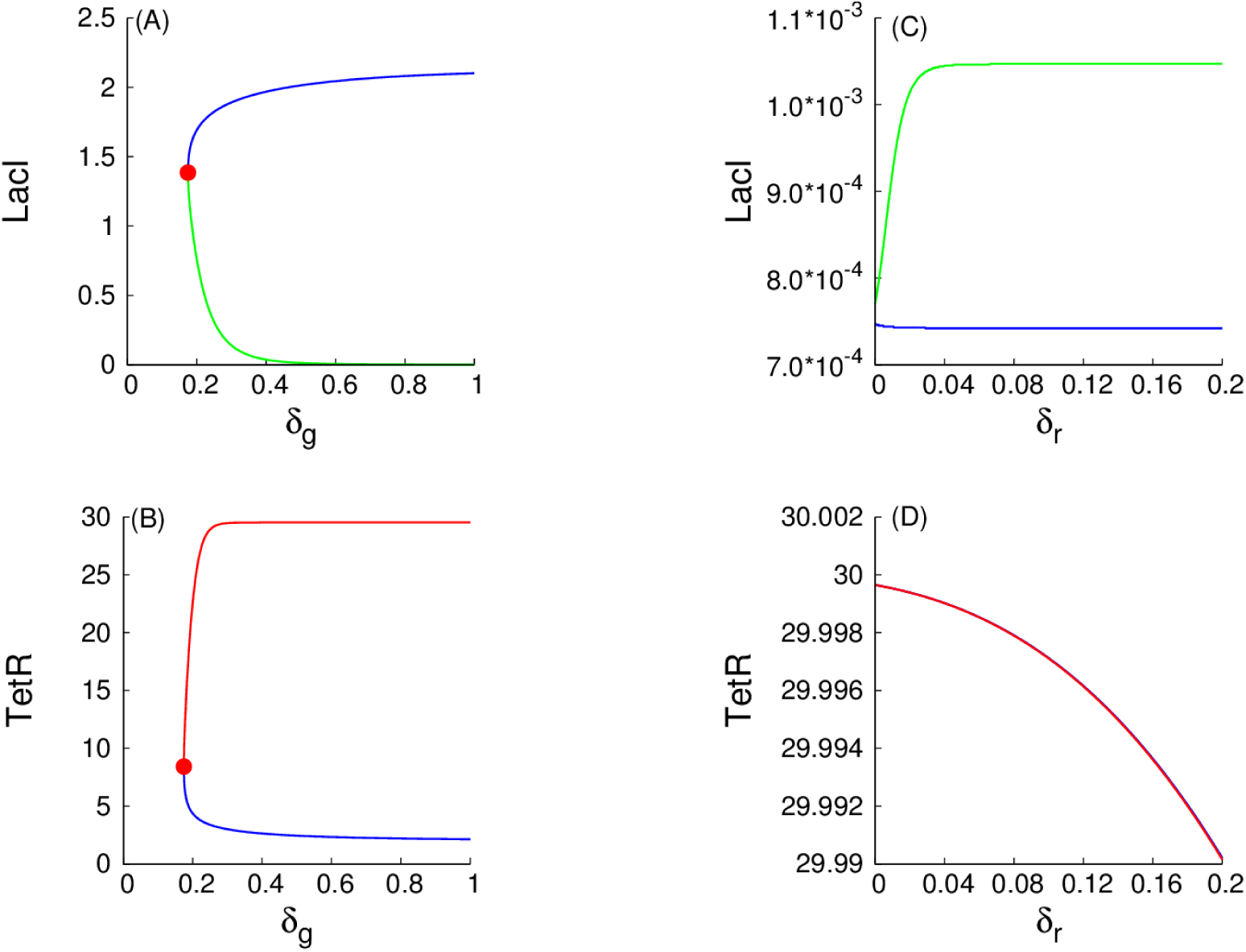
Examples of monotone parametric dependencies for the repressor-protein levels in the (9:1)-mixed state (a 10% small R-subpopulation.) The curves for the small R-subpopulation are shown in panels (A) (D), within which TetR is activated, while LacI is repressed. Panels (A) and (B) correspond to *δ_g_*. Panels (C) and (D) correspond to *δ_r_*. Green and red solid curves correspond to stable solution branches, while all blue curves correspond to unstable solution branches. In panel (D), projections of stable and unstable solution branches coincide. Red filled circles in panels (A) and (B) correspond to an LP-bifurcation point.

### Bistability in Single S-Design and A-Design Toggles Isolated from the Environment

Before comparing population properties of our S design to those of the A design, we remark that, even for isolated cells (when the diffusion constant *d* is zero), there is a larger range of bistability for the S design compared to the A design. Specifically, a bistability region for a single A toggle in the plane (*a*_1_, *a*_2_) at *d* = 0 is shown in Fig. 12. Similar regions were found in [1,13].

We observe that the entire quadrant, *a*_1_ *≥* 0 and *a*_2_ *≥* 0, spans a bistability region for the S-model at the fixed parameter values given in (3).

**Figure 12.**
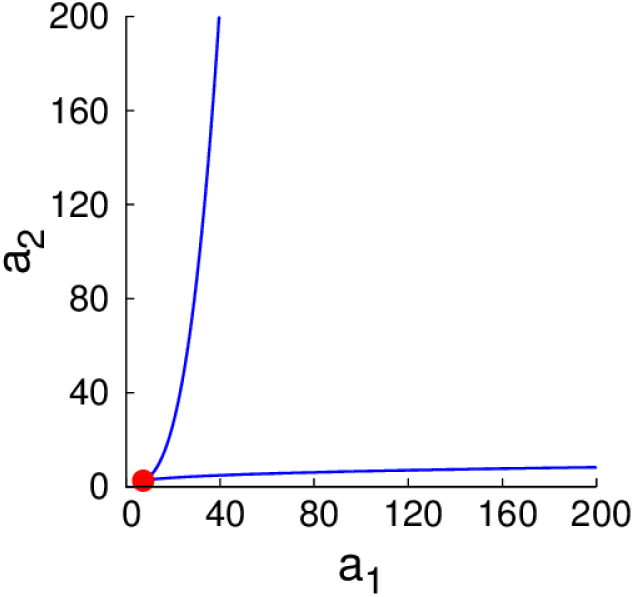
Bistability regions for single S toggle and A toggle models. The region between two blue color coded LP-bifurcation loci corresponds to a bistability region for the A toggle model (2) at *d* = 0. A red filled circle corresponds to a cusp point (CP). For the S toggle model, bistability exists for all parameter values *a*_1_ *≥* 0 and *a*_2_ *≥* 0 at *d* = 0. Other fixed parameter values are given in (3).

An important observation that follows immediately from Fig. 12 is that in the case of the S toggle, bistability exists at the origin of the non-negative quadrant in the plane (*a*_1_, *a*_2_), that is, at *a*_1_ = *a*_2_ = 0. This simply means that the genes *lacI* and *tetR* can be removed from the corresponding plasmids bearing promoters P_Y_ and P_X_, respectively (Fig. 2). In this case, it is enough to keep the genes on the plasmids bearing the corresponding promoters P_G_ and P_R_ (Fig. 2). We view the reduced S toggle as a minimal design that could be implemented experimentally. The fuller construct S is interesting too, in so far as it is based on the well-characterized and studied Cantor-Collins switch, coupled to quorum-sensing components [4]. We find that the full and reduced designs do not differ much in performance, and, so, we do not consider the minimal design in the rest of the paper.

### Bistable Homogeneous Populations Consisting of S toggles

Fig. 13 shows scaled levels of LacI and C14-HSL for a homogeneous population in the G-state, depending on the values of the parameter *d*. We observe that the intracellular and extracellular levels of the QS signaling molecule C14-HSL become asymptotically indistinguishable from one another as *d → ∞*.

The asymptotic behavior for large values of *d* can be analytically understood after introducing a small parameter *ε* = *d*^−1^ into the S-model (1) which becomes a singularly-perturbed problem [65].

**Figure 13.**
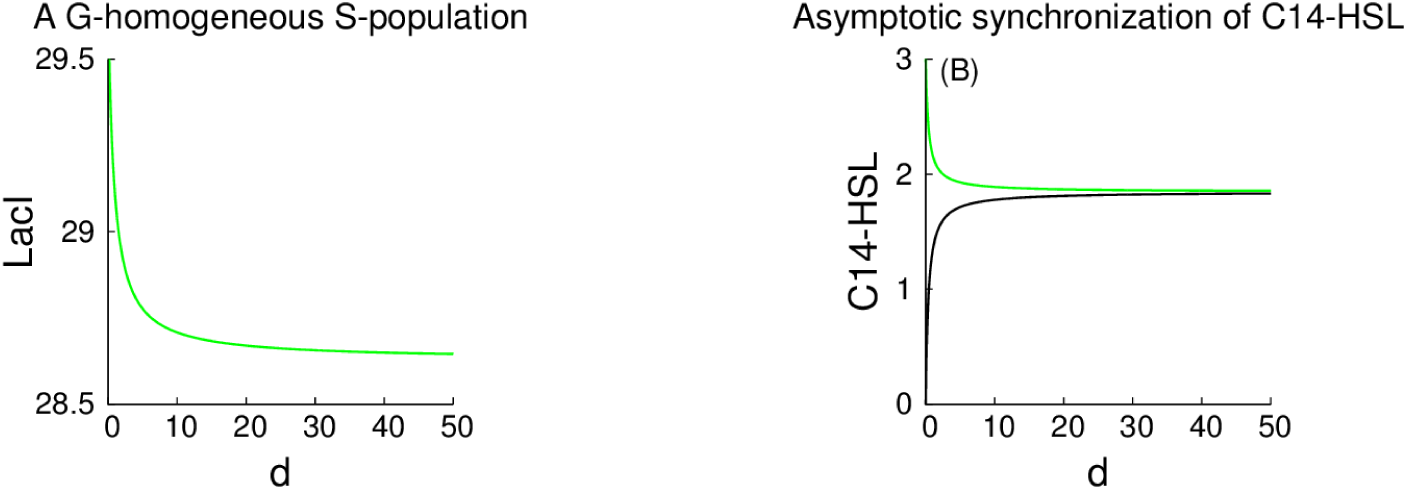
A G-homogeneous population of S toggles. Dependence of a G-homogeneous population on the values of the parameter *d* is shown. Panel (A) presents dimensionless levels of the activated LacI. Dimensionless levels of the repressed TetR are of order of magnitude about 10*^−^*^3^ and are not shown. Panel (B) presents dimensionless levels of C14-HSL. Here, the green curve corresponds to the intracellular levels, while the black plot corresponds to extracellular levels of C14-HSL in the medium, respectively.

Setting formally *ε* = 0 in the singularly-perturbed problem as required by the theory of singular perturbations [65], the differential equations (1c) and (1d) can be reduced to elementary algebraic equations *g* = *g*_*e*_ and *r* = *r*_*e*_, respectively.

### Bistable Homogeneous Populations Consisting of A toggles

Fig. 14 shows two stable homogeneous populations of A toggles which coexist while the parameter *d* is allowed to vary.

Because the A toggle design does not have typically any intrinsic symmetry, the levels of the activated repressor proteins, LacI for the G-homogeneous population shown in the Fig. 14(A,B) and TetR for the R-homogeneous population shown in Fig. 14(C,D), differ significantly from one another. Recall that the levels of LacI and TetR in the corresponding Gand R-homogeneous populations consisting of S toggles (Fig. 13) are identically the same due to the mirror (Z_2_) symmetry.

The combination of the analyses in Figs. 13 and 14 can be summarized by saying that under each one of the two designs, S and A, bistable homogeneous stable populations are possible, in either “Red” or “Green” consensus states, and with the same order of magnitude of expression. The difference between these designs is evident when we study heterogeneous (mixed) populations, as discussed next.

### A (1:1)-Mixed Population Consisting of S toggles

Fig. 15 shows a richness of dynamic effects (bifurcations) for a (1:1)-mixed population of S toggles. We see that as soon as the parameter *d* takes on larger values, the (1:1)-mixed state loses its stability via a Branch Point (BP) bifurcation (alternatively called “pitchfork” or “symmetry-breaking” bifurcation), giving rise to two stable (1:1)-mixed non-symmetric states at *d ≈* 1.43. The bifurcation has co-dimension one for all typical systems with Z_2_-symmetry [52,53].

To understand the symmetry-breaking phenomenon occurring at the BP-point shown in Fig. 15, we need first to define the symmetry of the original symmetric (1:1)-mixed state. We observe that the original symmetric (1:1)-mixed state is invariant with respect to transformation *g*,

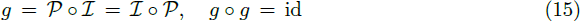

where id is the identity. Here, *𝒫* and *𝒤* are as defined in (9) and (10), respectively.

**Figure 14.**
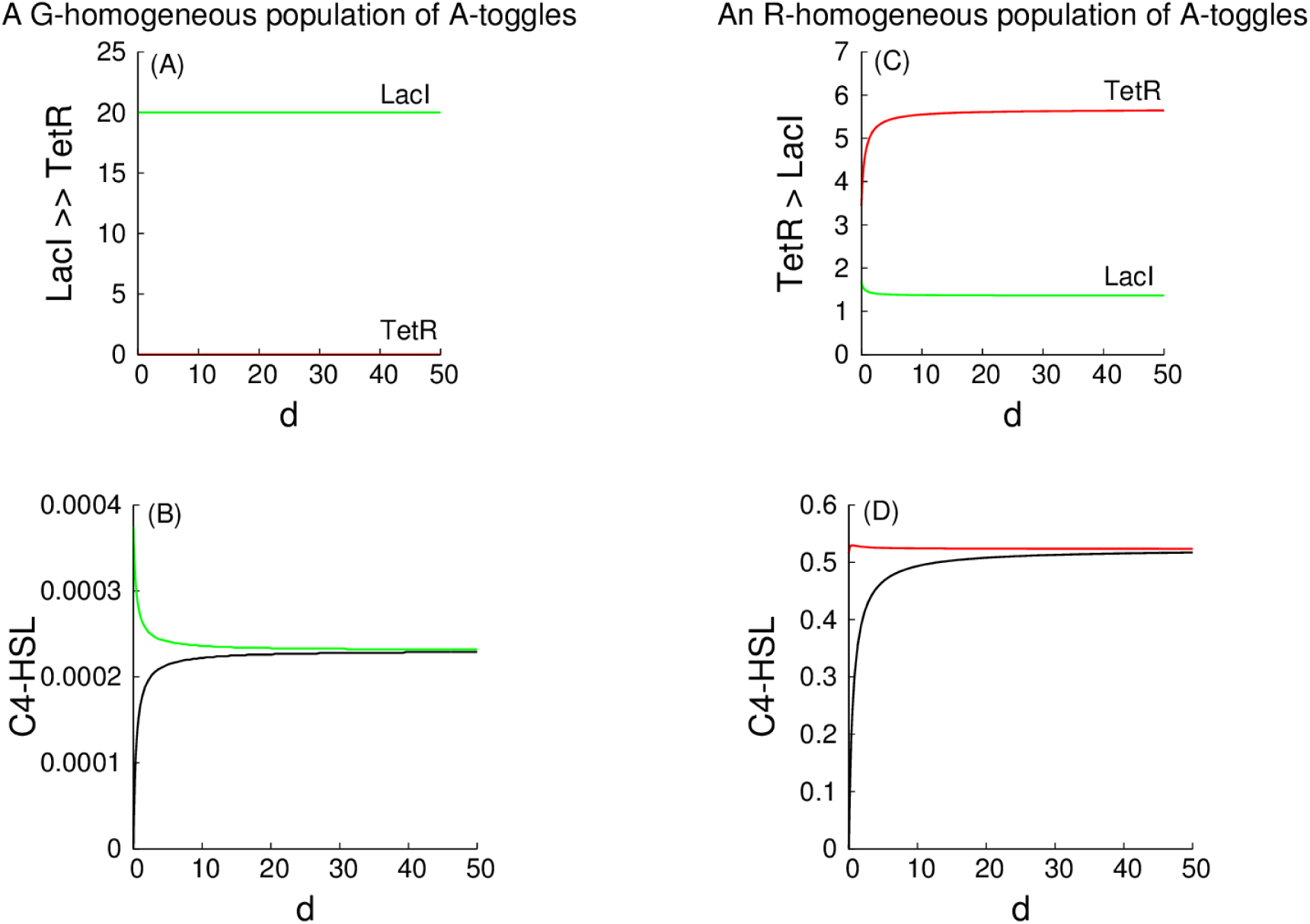
Gand R-homogeneous populations of A toggles. Panels (A) and (B) present dimensionless levels of the activated LacI and C4-HSL obtained for the A toggle settled at the G-state (LacI *>* TetR), while the other two panels (C) and (D) present dimensionless levels of the activated TetR and C4-HSL obtained for the A toggle settled at the R-state (TetR *>* LacI). Recall that C4-HSL is a unique QS signaling molecule facilitating communication between the A toggles, and this is why C4-HSL is present in both (B) and (D) panels.

**Figure 15.**
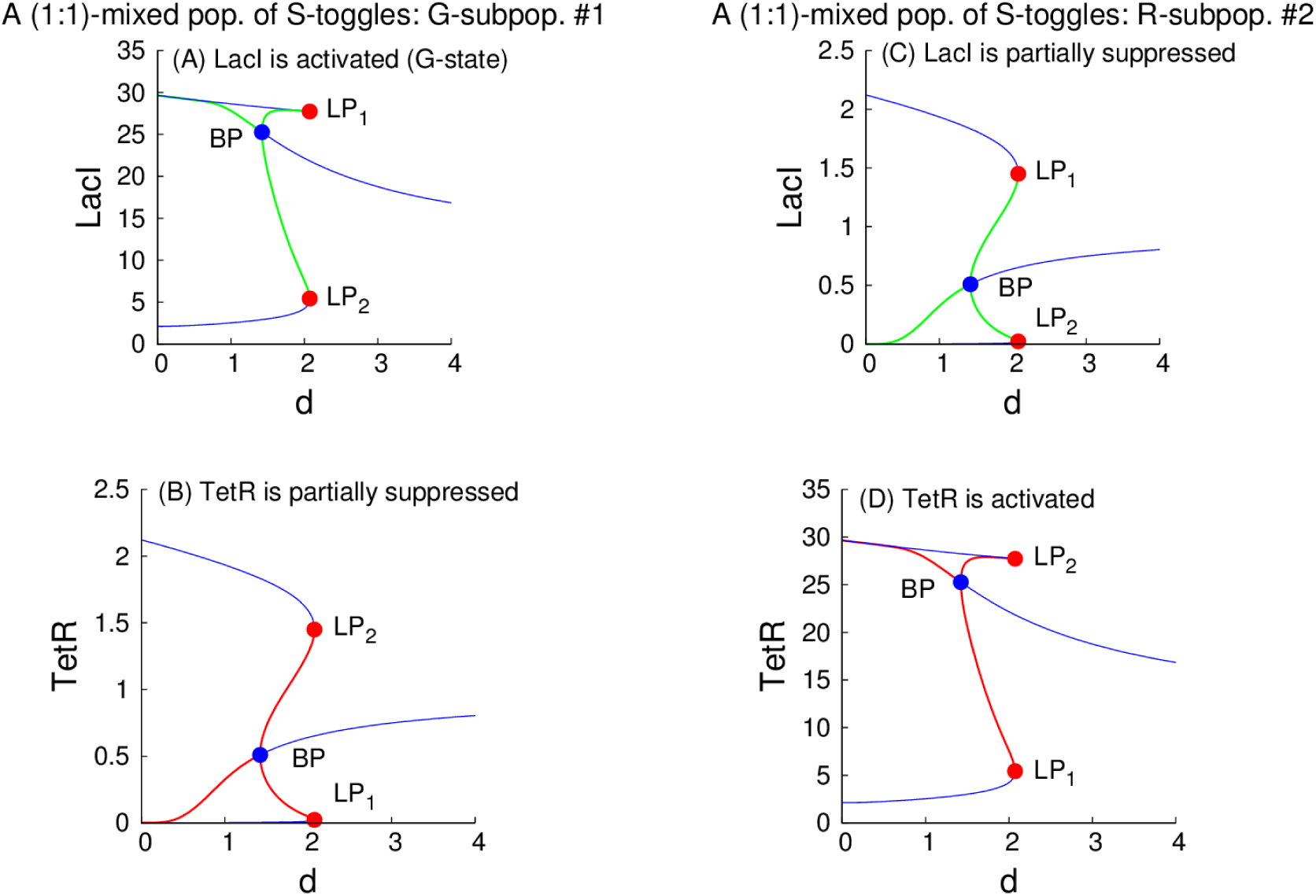
Symmetry breaking in a (1:1)-mixed population of S toggles. The given (1:1)-mixed population consists of two equal subpopulations, (1) a G-subpopulation and (2) an R-subpopulation, see the center panel in Fig. 3. Panels (A) and (B) show dimensionless levels of *activated* LacI (A) and *repressed* TetR (B), respectively, within the G-subpopulation. Panels (C) and (D) show dimensionless levels of *repressed* LacI (A) and *activated* TetR (B), respectively, within the R-subpopulation. Blue color-coded plots correspond to all *unstable* equilibrium solution branches, while green and red color-coded plots correspond to all *stable* equilibrium solution branches. All blue filled BP-labeled points correspond to *d ≈* 1.43. All red filled LP-labeled points correspond to *d ≈* 2.07. Both labels LP_1_ and LP_2_ correspond to the same bifurcation and are used for convenience of understanding the diagrams. Due to the *isotropy* subgroup Σ_(1:1)_ of the original (1:1)-mixed state, the isotropy involution *g* defined in (15) of the main text maps panel (A) to panel (D), and panel (B) to panel (C). We thus have, correspondingly, *g* : LP_1_ *→* LP_2_, *g* : LP_2_ *→* LP_1_, and *g* : BP *→* BP, see the coordinates of the three critical points, BP, LP_1_, and LP_2_, at the end of the caption. *Abbreviations:* pop., population; subpop., subpopulation. The coordinates of the critical points are given in the format (*x*_1_, *y*_1_, *x*_2_, *y*_2_; *d*), where index *i* corresponds to the subpopulation *i*, *i* = 1, 2, BP = (25.265, 0.510, 0.510, 25.265; 1.424), LP_1_ = (27.730, 0.023, 1.450, 5.427; 2.073), LP_2_ = (5.427, 1.450, 0.023, 27.730; 2.073).

The transformation (15) forms subgroup Σ(1:1) of the group Z_2_ *×* **S**_2_ (see Symmetry), which consists of two elements, that is, Σ_(1:1)_ = {id, *g*}. The subgroup Σ_(1:1)_ is called the *isotropy* subgroup [52] of the original (1:1)-mixed state. Because the two (1:1)-mixed states bifurcating from the original Σ_(1:1)_-symmetric (1:1)-mixed state at the BP-point (Fig. 15) are not invariant with respect to the isotropy subgroup Σ_(1:1)_, and, instead, are mapped to one another by the transformation (15), this motivates using the “symmetry-breaking” terminology [52] with respect to the loss of the isotropy symmetry by the (1:1)-mixed state at the BP-point.

As the value of the parameter *d* is further allowed to increase above the critical value *d ≈* 1.43 (Fig. 15), one branch of the pair of the stable non-symmetric (1:1)-states evolves along the locus connecting the points BP and LP_1_, while another branch of the same pair evolves along the locus connecting the points BP and LP_2_ (Fig. 15).

As the value of the parameter *d* is further increased, the (1:1)-mixed non-symmetric equilibria lose their stability and disappear via an LP-bifurcation at *d ≈* 2.07 (Fig. 15) simultaneously as relative states (see Symmetry). Indeed, we have *g* : LP_1_ *→* LP_2_, where *g* is defined in (15).

We can describe the above symmetry-breaking scenario intuitively as follows. Suppose that we start with a mixed population in which 50% of the cells are in “green” state and 50% of the cells are in “red” state, and the nondimensional diffusion coefficient *d* (which, as we saw, in fact incorporates many of the kinetic parameters in the original system) has a low value. Suppose that we now slowly increase the value of *d*, and ask what happens to the (1:1)-mixed state. The first event that is observed, at *d ≈* 1.43, is that this “pure 50-50 mixed state” loses its stability. A new mixed state arises, in which there are *two* subpopulations, one in which green gene-expression dominates (but with different expression levels of LacI in each of them), and another one which red gene-expression dominates (also with different TetR levels).

Furthermore, as *d* is increased a bit more (past *d ≈* 2.07), even these mixed states disappear. Thus, even with moderate diffusion, heterogeneous populations cannot be sustained, emphasizing the consensus-forming character of the S design. This is in marked contrast to the A design, as shown next.

The loss of stability by the (1:1)-mixed state increases the robustness of the S toggle design towards its self-synchronization by reducing the number of alternative stable states to which the toggle state can settle.

### A (1:1)-Mixed Population Consisting of A toggles

In contrast to (1:1)-mixed populations of S toggles, we observe from Fig. 16, that the original (1:1)-mixed A-population cannot be eliminated (made unstable) by increasing the values of the parameter *d* within a very large parameter interval. In other words, increasing the strength of interactions between the cells does not help to establish synchronization across the given population of identical A toggles.

### A (9:1)-Mixed Population Consisting of S-Toggles

Next, we consider bistable (9:1)-mixed populations, which as discussed in the introduction, we think of as arising from random synchronization errors. We observe that populations quickly become unstable as a function of the nondimensional diffusion parameter d (Fig. 17).

### (9:1)-and (1:9)-Mixed Populations Consisting of A toggles

In contrast to the S design, in the A design, the mixed (9:1)and (1:9)-heterogeneous populations that might arise from random state switching cannot be eliminated by changes in the values of the parameter *d* (Fig 18).

**Figure 16.**
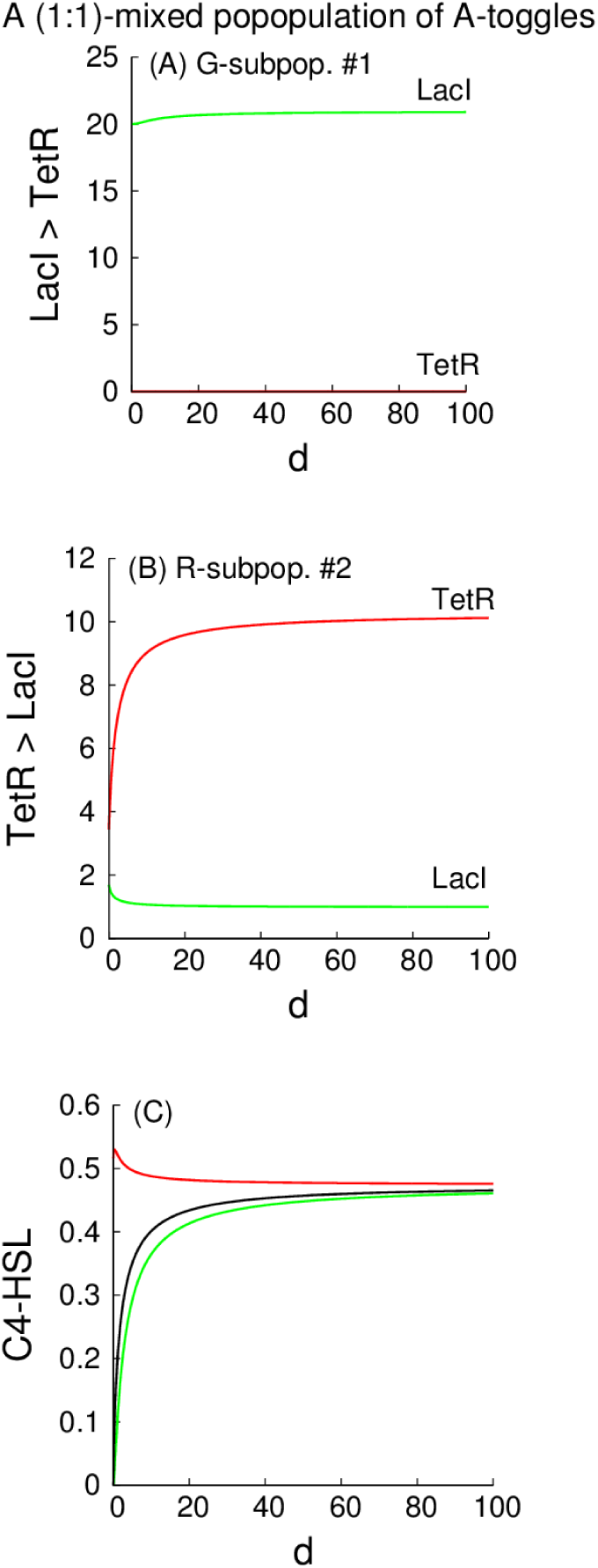
A (1:1)-mixed population of A toggles. Green and red color coded curves in panels (A) and (B) correspond to the intracellular concentrations of LacI and TetR, respectively, while a black color-coded curve in panel (C) corresponds to the extracellular concentration of C4-HSL. In panel (C), the green color-coded curve corresponds to the concentration of C4-HSL within the G-subpopulation, that is, LacI *>* TetR as in panel (A), while the red color-coded curve corresponds to the R-subpopulation, that is, TetR *>* LacI as in panel (B).

**Figure 17.**
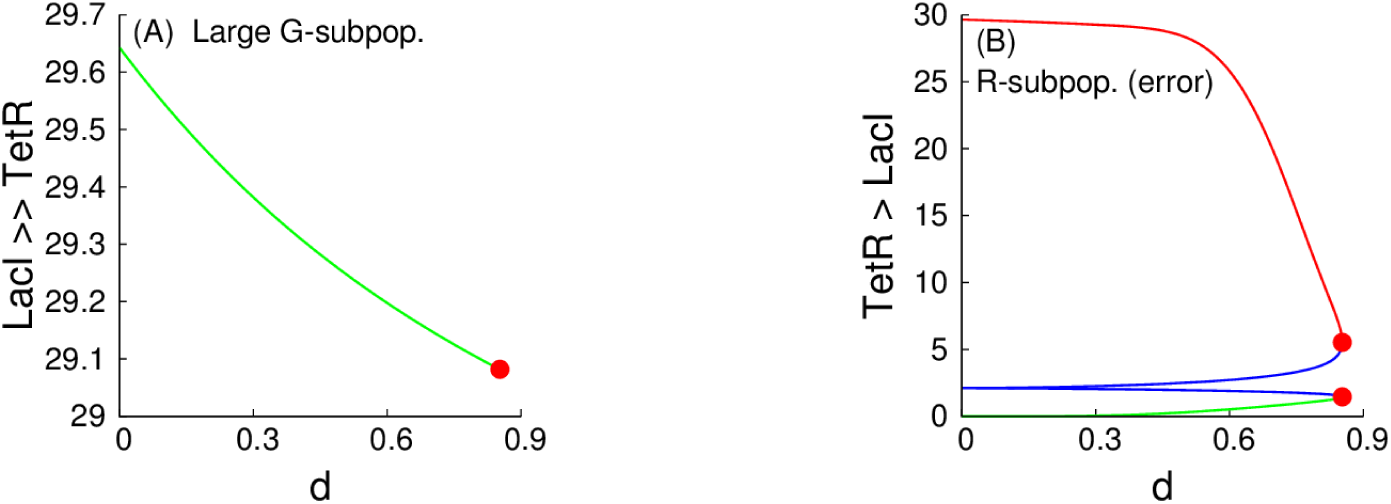
A (9:1)-mixed population of S toggles. Here, all notations and color-coding schemes are as in Fig. 15. Red filled circles correspond to the same LP-bifurcation point. In panel (A), projections of stable and unstable solution branches overlap. Because TetR is totally suppressed in the large (90%) subpopulation (G-subpopulation in the left panel), the levels of TetR are not shown. Instead, we zoom-in to a small range of the levels for LacI to obtain a better graphical visualization of the evolution of the LacI levels as *d* increases. Contrarily to panel (A), both TetR and LacI levels are plotted in panel (B) since LacI is only moderately suppressed in the small (10%) R-subpopulation.

**Figure 18.**
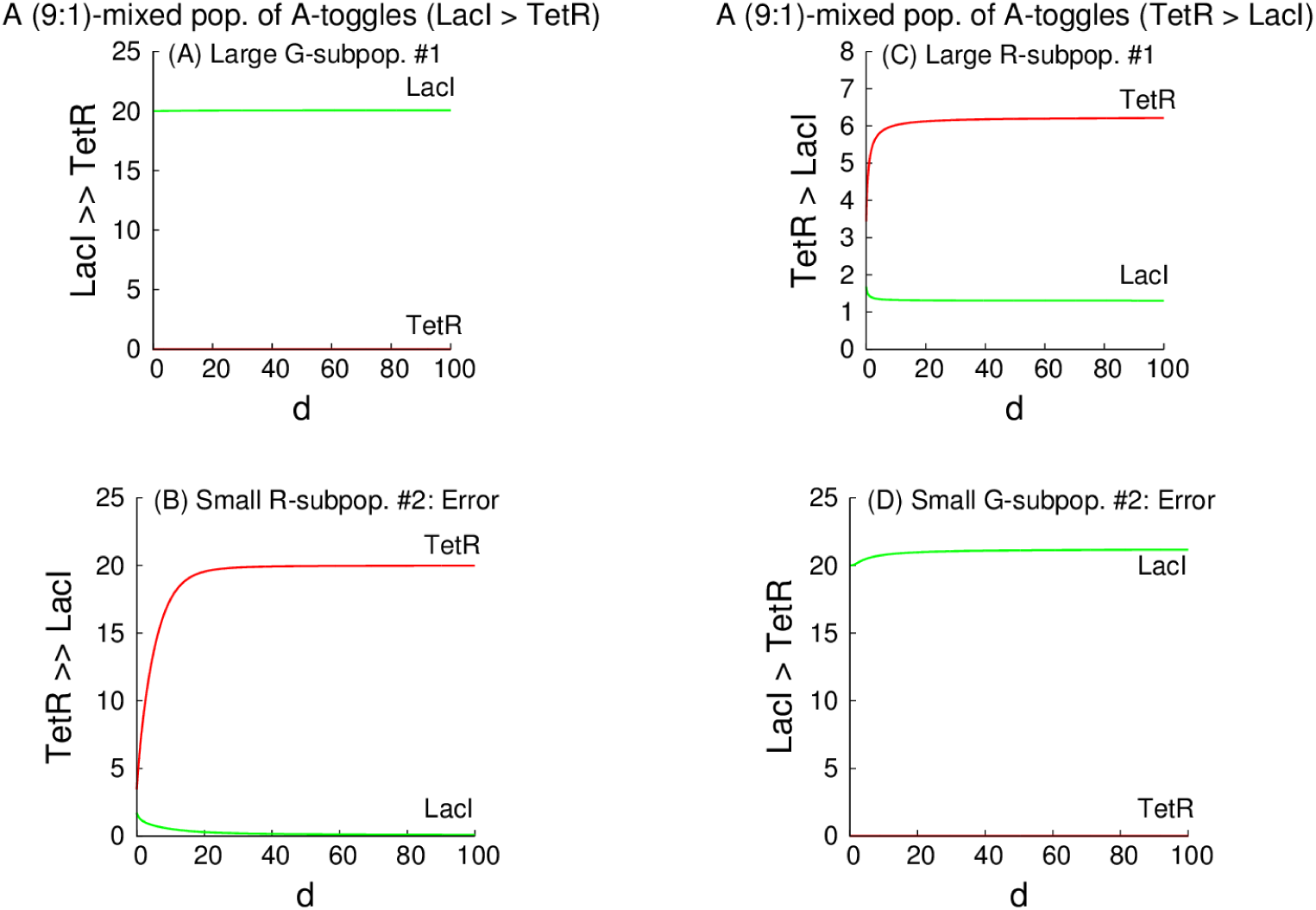
A (9:1)and (1:9)-mixed population of A toggles. Here, all notations and color-coding schemes are as in Fig. 17. Panels (A) and (B) correspond to the (9:1)-mixed population, within which the transcription signature LacI *≫* TetR dominates in proportion 9:1 (*i.e*., with 90% of green cells and 10% of red cells), while panels (C) and (D) correspond to the (1:9)-mixed population, within which the opposite transcription signature TetR *≫* LacI as well dominates in proportion 9:1 (*i.e*., with 90% of red cells and 10% of green cells.)

### Probing Capabilities of S- and A-Toggle Designs for Self-Correction of Spontaneous Synchronization Errors

To probe and compare capabilities of the S toggle and A toggle designs to correct “spontaneous synchronization errors” caused by a random flip of one toggle (or a small fraction of toggles) from a homogeneous population to the state opposite to the transcription signature adopted by the majority of the cells, we have performed simple random tests which will be discussed below. In mathematical and computational terms, these random tests can be interpreted as an elementary numerical procedure to evaluate the size of the basin of attraction for the corresponding equilibrium solutions by sampling the corresponding small neighborhoods of the solutions, using random initial conditions.

Our numerical procedure is organized as follows. For each of the parameter values in the following set: *d* ∈ {0.01, 10, 100}, two stable Gand R-homogeneous states have been found numerically, where the stability is computed for any value of *N* > 1, see S3 Exponential Stability of Cellular Populations. Due to the mirror symmetry of the S design, the analysis of the G-homogeneous state is sufficient as all trajectories are mapped to the corresponding trajectories for the alternative R-homogeneous state under the action of the involution defined in (7). In contrast, the property of error self-correction should be analyzed for two different A-homogeneous stable equilibria (Fig. 14) separately.

The random initial conditions at *t* = 0 have been selected as follows. First, we take *Z* = 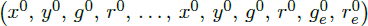 to be a (10:0)-homogeneous solution (G-state with *x*^0^ *> y*^0^). Then, the perturbed initial conditions for 9 cells were set to the G-state. The initial condition for the last, 10th cell was set at the R-state taken from the same the (10:0)homogeneous solution *Z*, using the involution *x*_10_(0) *← y*^0^, *y*_10_(0) *← x*^0^, *g*_10_(0) *← r*^0^, and *r*_10_(0) ← *g*^0^, that is, we obtained *x*_10_(0) *< y*_10_(0) (R-state). We also used 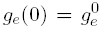 and 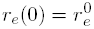. Each initial condition was then perturbed by adding a random term with a magnitude of 10% of the initial condition value for the corresponding state variable.

For the A design, two different sets of initial conditions were used as follows. First, let 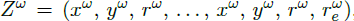, where *ω* ∈ {G, R}. Then, to form the fist set of initial conditions, the initial conditions for 9 cells were set at the G-state taken from the (10:0)-homogeneous solution, *Z*^G^, that is, we used *x*_*i*_(0) = *x*^G^, *y*_*i*_(0) = *y*^G^, *r*_*i*_(0) = *r*^G^ (*i* = 1, *… ,* 9), and 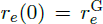, while the initial condition for the last, 10th cell was set at the R-state taken from the alternative, stable (0:10)-homogeneous solution, *Z*^R^, that is, we had *x*_10_(0) = *x*^R^, *y*_10_(0) = *y*^R^, and *r*_10_(0) = *r*^R^. The second set of initial conditions was formed, using similar steps, resulting in 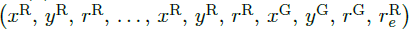. Finally, each initial condition was perturbed by adding a random term with a magnitude of 10% of the initial condition value for the corresponding state variable.

We can conclude from Fig. 19 and Fig. 20 that the A toggle does not have any capability for self-correction of spontaneous errors for all tested values of the parameter *d* (Fig. 20). The S toggle can self-correct spontaneous synchronization errors for medium and large values of *d* (Fig. 19, that is, for all parameters values for which the mixed state becomes unstable, see Fig. 17 (A (9:1)-Mixed Population Consisting of S-Toggles.)

**Figure 19.**
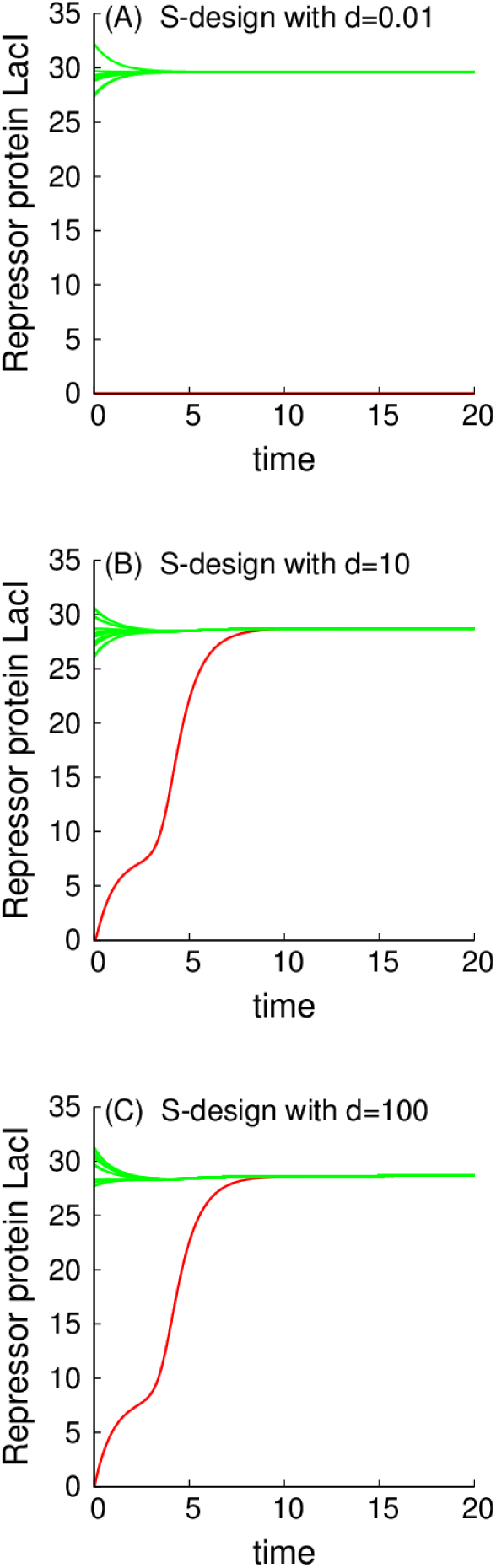
Probing self-correction capabilities of spontaneous errors by S toggles. All explanations are given in the main text.

**Figure 20.**
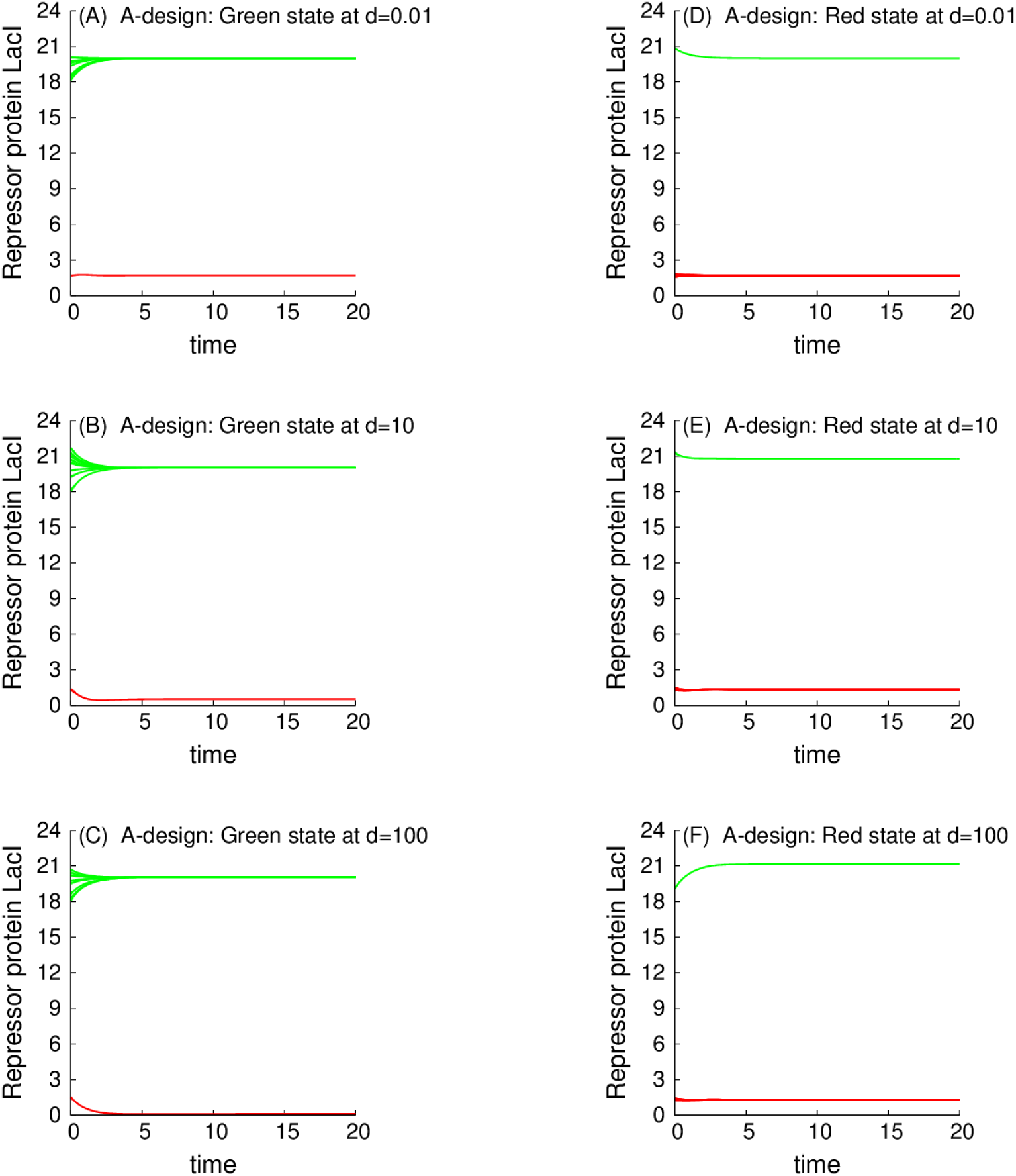
Lack of any self-correction capability for spontaneous errors by A toggles. The left panels correspond to the reference (10:0)-state (G-state), while the right panels correspond to the reference (0:10)-state (R-state). The initial conditions in the left panels correspond to nine “green” cells and one “red” cell. The initial conditions in the right panels correspond to one “green” cell and nine “red” cells.

## Conclusions

In this study, we have shown how synthetic bistable circuits (toggles), and hosting them, programmable cellular populations, can be designed so as to solve a robust molecular task, the maintenance of a coordinated state, and a “majority-vote” auto-correction of deviations, of a binary switch. Our design was guided by concepts from monotone systems theory [15–21]. Specifically, we have shown how this concept can be used for the design of a new class of monotone synthetic biological toggles, including predictive capabilities describing both dynamic state variables and monotone parametric tendencies caused by parameter perturbations.

To benchmark the new toggle design, termed the S design, and the monotone systems approach, we have compared the S design with the known (and non-monotone) asymmetric B2-strain from [4], termed the asymmetric or A design in this work. The B2-strain has been previously studied both experimentally [4] and theoretically [13,14]. Despite a number of remarkable properties of the B2-strain (A design), the A toggle multifunctionality suggests that the design must be tightly controlled to avoid spontaneous switching not only between different expression states, but, as well, between different functions such as a bistable memory and an oscillatory phenotype.

In this respect, modern gene therapy interventions are currently limited to transfected genes to be either in an “on” or “off” state, when the expression of the transfected gene needs to be regulated tightly for the effective treatment of many diseases. To address this challenge, the monotone S toggle design completely excludes any unpredictable chaotic behaviors, as well as undesired stable oscillations. This conclusion is valid for all parameter values, and provides a strong theoretical guarantee missing from other synthetic biology designs.

To achieve an in-depth understanding of dynamic properties of the S toggle design, we have developed biochemically-detailed and biologically-relevant mathematical models to test predictions of monotone systems theory by employing computational bifurcation analysis. To have all results biologically grounded, concrete molecular entities have been used, though the results are general and independent of any specific details.

Additionally, to investigate the effect of a spontaneous toggle switching within cellular populations, leading to bimodal distributions, we have formalized a concept of spontaneous synchronization errors and tested the toggle design capabilities to self-correct spontaneous synchronization errors by sampling the basin of attraction of the corresponding equilibrium solutions. We found that the S toggle design was able to self-correct (or, auto-correct) synchronization errors, while the non-monotone A toggle design was not.

Finally, because the number of cells in populations is *a priori* unknown, all the above results and conclusions can make sense only if they are made independently of the population size. To justify the above assertion, we have proved a few general theorems on the exponential stability of the equilibrium solutions corresponding to both homogeneous and mixed populations. The simple exponential stability results are independent of the number of cells in the populations and are based on basic first principles of stability analysis resulting from the Schur’s formula [66], which allow the characteristic polynomials for the corresponding model linearizations to be computed explicitly.

## Supporting Information

### S1 Model Derivation

Here, we describe the main assumptions and steps used to derive mass-balance equations for the Sand A-models formulated in the main text.

#### S1.1 Mass-Balance Equations

The derivation of the Sand A-models includes the development of two modules:

I. A transcription-translation module describing biosyntheis of repressor proteins.

II. A metabolic module describing biosynthesis of autoinducers.

A general and systematic discussion of both modules can be found in [67,68]. The derivation of the first module for the A-model, relevant to our work, is given in [13]. Because one of our modeling objectives is to ultimately describe how the analysis of the mathematical models can be mechanistically interpreted in terms of tuning synthetic toggle “dials” by implementable experimental interventions as reviewed in [6], including modifications of ribosome-binding sites (RBS), carboxy-terminal tags, *etc*., [1,2,51], we will derive mass-balance equations at the level of molecular detail sufficient to suggest plausible modeling predictions.

A transcription-translation module can be described by a basic two-stage model [13,67,68],

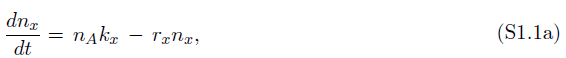

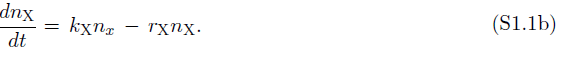

Here, *n*_*x*_ is the number of mRNA transcripts per cell for gene *x*, and *n*_X_ is the number of protein molecules per cell; *n*_*A*_ is the number of active promoters from which the mRNA of gene *x* is transcribed at an average rate *k*_*x*_; *k*_X_ is the averaged translation rate; *r*_*x*_ and *r*_X_ are the effective first-order rate constants associated with degradation of the mRNA and proteins, respectively.

Since mRNA molecules are usually degraded rapidly compared to other cellular processes, a quasi-steady state for the equation (S1.1a) can often be assumed [13], yielding

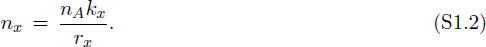

Using (S1.2) in the right-hand side of the equation (S1.1b), we obtain

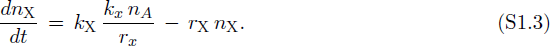

The ratio *b*_*x*_ = *k*_X_*/r_x_* in equation (S1.3) is called a burst parameter of the protein X [13]. Using *b*_*x*_ in (S1.3) yields

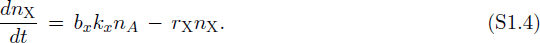

Assuming log-phase growth of *E. coli*, the volume *V* (*t*) of the growing bacterium can be approximated by the expression *V* (*t*) = *V*_0_ exp(*λt*), and equation (S1.4) can be rewritten in a concentration form,

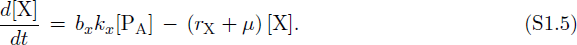

Here, [X](*t*) = *n*_X_(*t*)*/V* (*t*) and [P_A_](*t*) = *n*_*A*_(*t*)*/V* (*t*).

The concentration of activate promoters, [P_A_], can be computed, using an appropriate Hill function [68]. For example, we use

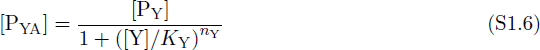

for the repressor protein Y binding to the promoter P_Y_ with the dissociation constant 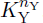. In (S1.6), [P_Y_] is the total concentration of all promoters P_Y_, while [P_YA_] is the concentration of active promoters not bound with the repressor protein Y. Recall that the cooperativity described by the Hill exponent *n*_Y_ can arise from [1,67,68]:

i. Multimerization of represssor proteins;
ii. Cooperative binding of repressor multimers to multiple operator sites in the promoter.

Analogously, we use a Hill-function

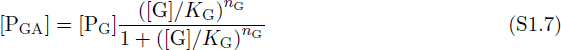

for the autoinducer G binding to the promoter P_G_ with the dissociation constant 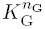 and the Hill exponent *n*_G_. The concentration of all active promoters P_A_ can now be obtained from (S1.6) and (S1.7) as

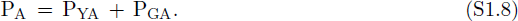

Using expression (S1.8), the equation (S1.5) can finally be updated as

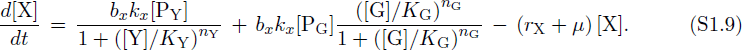

Here, all parameters are described below in Table S2.2.

Similar mass balanced equations can be derived for the repressor protein Y, and synthases U and W. For example, the mass balance equation for the synthase U is

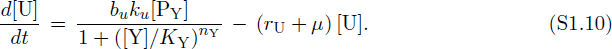

Analogously, we can write down a mass balance equation for the autoinducer concentration [G], that is, [C14-HSL], governed by the synthase U (CinI),

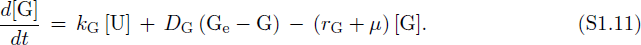

Here, *k*_G_ is the maximum production rate of C14-HSL by CinI (Table S2.4), *D*_G_ is the export rate of C14-HSL (Table S2.4), and [G_e_] is the extracellular concentration of C14-HSL.

Assuming that the concentration of the enzyme U reaches its quasi-steady state rapidly [69], one can obtain from (S1.10) that

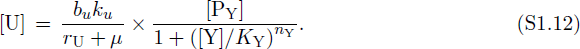

Using (S1.12) in the equation (S1.11) yields

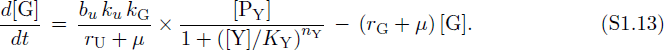

Here, the definitions and the values of all parameters are given in Table S2.4. A similar mass balanced equation can be derived for the second autoinducer R (C4-HSL), and we omit the details.

#### S1.2 Nondimensionalization (Scaling)

To nondimensionalize mass balance equations, as for example, the mass balance equations (S1.9) and (S1.13), we use the following dimensionless state variables, which are similar to those introduced in [13],

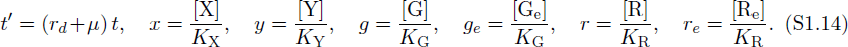

Here, we assume that all protein degradation rates can be set experimentally so that the following equalities can be obtained *approximately* [2],

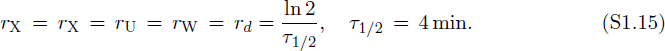

The procedure of setting all protein degradation rates or, equivalently, all protein half-lives approximately equal to a prescribed value close to mRNA half-lives [2] is required to balance the toggle [1]. We discuss the balancing procedure and relevant experimental interventions in Sect. S1.3.

Using the dimensionless variables (S1.14), all original modeling mass balances can be nondimensionalized, yielding the Sand A-models formulated in the main text, where the prime is dropped from dimensionless time *t*′. In this case, dimensional and dimensionless parameters are related to one another as:

1. For the dimensionless rates, we obtain:

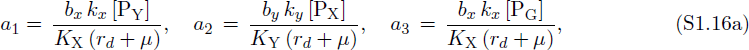

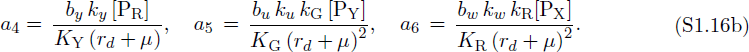

2. For dimensionless diffusion and degradation parameters, we obtain:

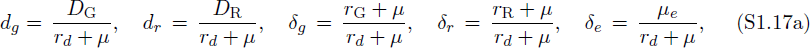

Molecular and biophysical parameter values used in the expressions (4) (5) will be estimated in Sect. S1.3, while dimensionless parameters will be estimated in Sect. S1.3 In this section, we only mention that due to [2], we can set

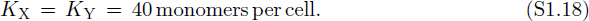

We could not find any estimation of values for the two parameters *K*_G_ for C14-HSL and *K*_R_ for C4-HSL in the literature despite the fact that more and more precise measurements of kinetic parameters become available [27]. We estimate the order of magnitude of *K*_G_ and *K*_R_ as follows.

In the detailed experimental results on the C4-HSL-mediated quorum sensing regulatory system of the opportunistic Gram-negative bacterium *Aeromonas hydrophila*, the concentration of C4-HSL was found to be of order of magnitude equal to 10 *μ*M [70]. In *E.coli* biology, it is convenient to use nM units [38], because relative to the effective *E. coli* volume [43], the value of 1 nM corresponds to one molecule per cell. This fact is widely used in the literature [2]. Therefore, the above estimate of 10 *μ*M corresponds to 10^4^ C4-HSL signaling molecules per cell.

Another *ad-hoc* rule of *E.coli* biology used in a number of studies with the Cornell *E.coli* computer model [37,44,71,72], resulting in a number of relevant predictions such as ribosomal-protein limitations, *lac*-control, plasmid stability, and *etc*. [30,39–43,45], is that, the coarse-grained estimation for the dissociation equilibrium constant to be used in the Hill function can be calculated as 25% of the intracellular modifier (reference) concentration. In our case, this yields a coarse-grade estimate of 0.25 *×* 10^4^ C4-HSL signaling molecules per cell,

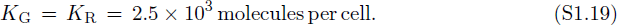

The values for other parameters will be estimated below.

#### S1.3 Toggle Balancing (Symmetrization)

As it was observed experimentally [1,2], synthetic circuits can operate and deliver the engineered traits only if special molecular constrains are fulfilled,

*(i)* repressor protein half-lives are close to mRNA halflifes [2];

*(ii)* repressor protein half-lives are approximately equal [1,2].

Constrain *(ii)* is required for “balancing” the given circuit [1]. Moreover, both works suggest experimental interventions to fulfill the above constraints [1,2]. Such and similar interventions are termed a “tuning dials” in the review [6].

It is mathematically convenient for us to generalize the above balancing procedure by the procedure of “symmetrization” of two antagonistic, mutually repressing toggle subsystems by selecting synthetic (tuned) parameter values that would make two antagonist subsystems symmetric to one another. In other words, we assume that an *ideal* S toggle has a mirror ℤ_2_-symmetry group of permutations between the two antagonistic subsystems. Symmetry usually helps with analytical analysis of nonlinear mathematical models.

Specifically, we “symmetrize” (balance) biosynthesis kinetic rates, using constraints

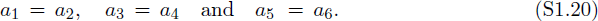

Appropriate molecular interventions, which can be used to set the relationships (S1.20) approximately under certain experimental conditions, are reviewed in [6]. Similarly, we symmertize “diffusion” parameters,

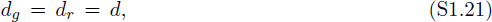

and the autoinducer “degradation” or “utilization” (“load”) parameters,

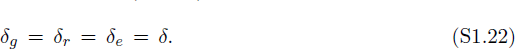

### S2 Estimation of Parameter Values

#### Toggle Parameters

In our estimations, we use general biochemical calculations [73,74]. First, we collect generic prokaryotic and specific *E.coli* parameter values in Table S2.1.

**Table S2.1.** Generic prokaryotic and *E. coli* specific values of model parameters

| Name | Description | Value | Unit | Reference |
| --- | --- | --- | --- | --- |
| $r_{\text{mRNA}}$ | Rate of transcription by RNA polymerase in prokaryotes | 80 | bp/sec | [46] |
| $\tau_{\text{mRNA}}$ | Typical half-life time for 80% of genes in *E. coli* | 3 - 8 | min | [75] |
| $r_{\text{aa}}$ | Rate of translation by the ribosome in prokaryotes | 20 | aa/sec | [46]<sup>a</sup> |
| $k_{\text{P}}$ | Rate of translation by the ribosome in prokaryotes | 1.71 | sec<sup>-1</sup> | estimated in (S2.3) |
| $T$ | *E. coli* replication period under specific nutrition conditions | 25 | min | [43] |
| $\mu$ | Intracellular specific dilution rate due to *E. coli* cell growth | $\ln 2/T$ | min<sup>-1</sup> | [43] |
| $\mu_e$ | Extracellular dilution rate due to flow | 0.1 | min<sup>-1</sup> | [76] |
| $\rho$ | Total volume fraction of cells in chamber | 0.8 | — | [77] |
| $N$ | Number of *E. coli* cells in an overnight population culture | $10^9$ (OD600 = 1) | cells/ml | [43] |

<sup>a</sup>This estimate is smaller than the estimate 33 aa/sec used in [27].

**A general rate of translation of protein P in prokaryotes** (*k*_P_). Suppose that an mRNA transcript of protein P contains *n*_aa_ amino acids. Then, for one ribosome to transcribe P from its mRNA transcript, assuming a translation rate of 20 amino acids per second (Table S2.1), it will take time

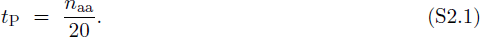

The above estimates yields the rate per ribosome which is

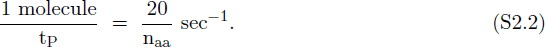

Given that the coding region of protein P is *n*_aa_ *×* 3 nucleotides long, and that a ribosome can attach every 35 nucleotides, we can estimate that *n*_aa_ *×* 3*/*35 ribosomes can be attached per mRNA molecule. We, thus, obtain

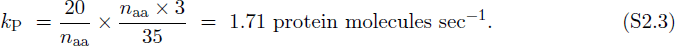

To illustrate our parameter estimation procedure, we derive parameter values for the Lac-repressor subsystem only. Parameter values for all other subsystems can be derived similarly.

*k*_*x*_ : **Fully induced strength of promoters** P_Y_ (P_tet_) **and** P_G_. One *lacI* mRNA transcript is 1204 bases long (Table S2.2). To transcribe one molecule of *lacI* mRNA from one gene with a rate of 80 bases per second (Table S2.2) takes

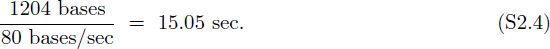

**Table S2.2.** Parameter values of the LacI-repressor subsystem.

| Name | Description | Value | Unit | Reference |
| --- | --- | --- | --- | --- |
| <u>Transcription:</u> |  |  |  |  |
| $L_{lacI}$ | Size of *lacI* gene | 1204 | bp | [78] |
| $\Gamma_x$ | Repressed strength of promoter $P_{tet}$ | $5.0 \times 10^{-4}$ | molecules/cell/sec | [2] |
| $k_x$ | Fully induced transcription rate<sup>a</sup> | $6.65 \times 10^{-2}$ | species/cell/sec | estimated in (S2.5) |
| $K_Y$ | The number of TetR to repress<sup>b</sup> $P_{tet}$ | 40 | monomers/cell | estimated in (S1.18) |
| $K_G$ | The number of C14-HSL to activate<sup>c</sup> $P_{cin}$ | $2.5 \times 10^3$ | molecules/cell | estimated in (S1.19) |
| $\tau_{1/2, lacI}$ | Half-life of *lacI* mRNA | 3.8 | min | [35] |
| $r_x$ | Rate of *lacI* mRNA degradation | $3.04 \times 10^{-3}$ | sec<sup>-1</sup> | estimated in (S2.7) |
| $n_Y$ | The number of subunits in TetR | 2 | | [34] |
| $n_G$ | Hill coefficient of C14-HSL | 4 | | [1] |
| <u>Translation:</u> |  |  |  |  |
| $L_{LacI}$ | Size of one subunit in tetrameric LacI | 360 | aa/subunit | [33] |
| $k_X$ | Rate of LacI translation | 1.71 | molecules/cell/sec | estimated in (S2.3) |
| $\tau_{1/2, LacI}$ | Half-life of LacI protein | 4 | min | [2] |
| $r_X$ | Rate of LacI degradation | $1.16 \times 10^{-3}$ | sec<sup>-1</sup> | estimated in (S2.7) |

<sup>a</sup>Fully induced strength of promoters $P_{tet}$ and $P_{cin}$ ,
<sup>b</sup>For the sake of brevity, “to repress” means “to half-maximally repress.”
<sup>c</sup>For the sake of brevity, “to activate” means “to half-maximally activate.”

Then, per gene the estimate (S2.4) yields

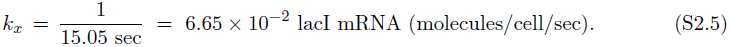

The estimate (S2.4) is one order of magnitude less than the estimate 0.5 lacI mRNA (molecules*/*cell*/*sec) provided in [2].

*r*_*x*_: **Rate of *lacI* mRNA degradation.** The calculation of degradation rates for proteins is based on the known protein half-lives,

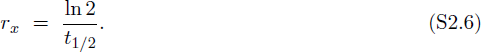

We obtain (Table S2.2),

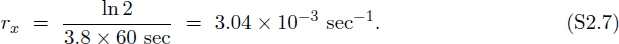

*r*_X_ : **Rate of LacI (X) degradation.** We obtain (Table S2.2),

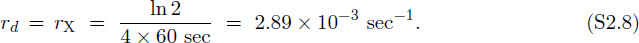

We use the estimate (S2.8) for all proteins in the model.

**Dimensionless parameter values**

Using data from Table S2.1 and the estimate (S2.8), we obtain

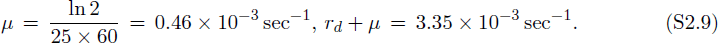

**Table S2.3.** Parameter values of the TetA-repressor subsystem.

| Name | Description | Value | Unit | Reference |
| --- | --- | --- | --- | --- |
| <u>Transcription:</u> |  |  |  |  |
| $L_{tetR}$ | Size of *tetR* gene | 905 | bp | [79] |
| $\Gamma_y$ | Repressed strength of promoter $P_{lac}$ | $5 \times 10^{-4}$ | molecules/cell/sec | estimated in Table S2.2 |
| $k_y$ | Fully induced transcription rate<sup>a</sup> | $8.87 \times 10^{-2}$ | molecules/cell/sec | estimated |
| $K_X$ | The number of LacI to repress<sup>b</sup> $P_{lac}$ | 40 | monomers/cell | estimated in Table S2.2<sup>d</sup> |
| $K_R$ | The number of C4-HSL to activate<sup>c</sup> $P_{rhl}$ | $5 \times 10^4$ | monomers/cell | estimated in Table S2.2 |
| $\tau_{1/2,tetR}$ | Half-life of *tetR* mRNA | 0.5 | min | [32] |
| $r_y$ | Rate of *tetR* mRNA degradation | $2.31 \times 10^{-2}$ | sec<sup>-1</sup> | estimated |
| $n_X$ | The number of subunits in LacI | 2 | | [34] |
| $n_R$ | Hill coefficient of C4-HSL | 4 | | estimated in Table S2.2 |
| <u>Translation:</u> |  |  |  |  |
| $L_{TetR}$ | Size of one subunit in tetrameric TetR | 207 | aa/subunit | [79] |
| $k_Y$ | Rate of TetR translation | 1.71 | molecules/cell/sec | estimated in (S2.3) |
| $\tau_{1/2,TetR}$ | Half-life of TetR protein | 4 | min | estimated Table S2.2 |
| $r_Y$ | Rate of TetR degradation | $1.16 \times 10^{-3}$ | sec<sup>-1</sup> | estimated |

<sup>a</sup>Fully induced strength of promoters $P_{lac}$ and $P_{tet}$ ,
<sup>b</sup>For the sake of brevity, “to repress” means “to half-maximally repress.”
<sup>c</sup>For the sake of brevity, “to activate” means “to half-maximally activate.”
<sup>d</sup> Equilibrium dissociation constant for LacI is $7.7 \times 10^{-8}$ M [80].

**Table S2.4.** Parameter values of the 3-OH-C14-HSL/CinI-signaling subsystem.

| Name | Description | Value | Unit | Reference |
| --- | --- | --- | --- | --- |
| <u>Transcription:</u> |  |  |  |  |
| $L_{cinI}$ | Size of *cinI* gene<sup>a</sup> | 663 | bp | estimated |
| $\Gamma_u$ | Repressed strength of promoter $P_{lac}$ | $5 \times 10^{-4}$ | molecules/cell/sec | estimated in Table S2.2 |
| $k_u$ | Fully induced strength of promoter $P_{tet}$ | $13.27 \times 10^{-2}$ | molecules/cell/sec | estimated |
| $\tau_{1/2,cinI}$ | Half-life of *cinI* mRNA | 6.6 | min | arbitrary<sup>b</sup> |
| $r_u$ | Rate of *cinI* mRNA degradation | $1.75 \times 10^{-3}$ | sec<sup>-1</sup> | estimated |
| <u>Translation:</u> |  |  |  |  |
| $L_{CinI}$ | Size of CinI aa-sequence | 221 | aa | http://string-db.org |
| $k_U$ | Rate of CinI translation | 1.71 | molecules/cell/sec | estimated in Table S2.2 |
| $\tau_{1/2,CinI}$ | Half-life<sup>a</sup> of CinI protein | 4 | min | [2] |
| $r_U$ | Rate of protein CinI degradation | $1.16 \times 10^{-3}$ | sec<sup>-1</sup> | estimated in Table S2.2 |
| <u>Signaling:</u> |  |  |  |  |
| $k_G$ | Maximal production rate of CinI | 2 | min<sup>-1</sup> | [81] |
| $r_G$ | Degradation rate of C14-HSL | 0.002 | hr<sup>-1</sup> | [82]<sup>c</sup> |
| $D_G$ | Export rate of C14-HSL | 2.1 | min<sup>-1</sup> | [83] |

<sup>a</sup>The coding region of the gene has been estimated from its protein sequence size provided in the same table as $221 \times 3 = 663$ .
<sup>b</sup>This estimate corresponds to a general (or typical) pattern for mRNA half-lives in *E. coli* [84]. Note that [2] use a generic half-life parameter value of 2 min.
<sup>c</sup>Data for 3-OH-C12-HSL is used.

**Table S2.5.** Parameter values of the C4-HSL/RhlI-signaling subsystem.

| Name | Description | Value | Unit | Reference |
| --- | --- | --- | --- | --- |
| <u>Transcription:</u> |  |  |  |  |
| $L_{rhII}$ | Length of *rhII* gene | 603 | bp | [85] |
| $\Gamma_w$ | Repressed strength of promoter $P_{lac}$ | $5 \times 10^{-4}$ | molecules/cell/sec | estimated in Table S2.2 |
| $k_w$ | Fully induced strength of promoter $P_{lac}$ | $13.27 \times 10^{-2}$ | molecules/cell/sec | estimated |
| $\tau_{1/2, rhII}$ | Half-life of *rhII* mRNA | 6.6 | min | [86]<sup>a</sup> |
| $r_w$ | Rate of *lacI* mRNA degradation | $1.75 \times 10^{-3}$ | sec<sup>-1</sup> | estimated |
| <u>Translation:</u> |  |  |  |  |
| $L_{LacI}$ | Length of RhlI protein aa-sequence | 196 | aa | [87] |
| $k_W$ | Rate of RhlI translation | 1.71 | molecules/cell/sec | estimated in Table S2.2 |
| $\tau_{1/2, RhlI}$ | Half-life of RhlI protein | 4 | min | estimated in Table S2.2 |
| $r_X$ | Rate of LacI degradation | $1.16 \times 10^{-3}$ | sec<sup>-1</sup> | estimated |
| <u>Signaling:</u> |  |  |  |  |
| $k_R$ | Maximal production rate of C4-HSL by RhlI | 16 | min<sup>-1</sup> | [81] |
| $r_R$ | Degradation rate of C4-HSL | 0.02 | hr<sup>-1</sup> | [82] |
| $D_R$ | Export rate of C4-HSL | 3.0 | min<sup>-1</sup> | [83] |

<sup>a</sup>The half-life data for *lasI* mRNA is used because the degradation of *rhII* is positively regulated by *LasI* [86] and, so, could have a longer half-life. This estimate is in line with a general (or typical) pattern for mRNA half-lives in *E. coli* [84]. Note that [2] use a genetic half-life parameter value of 2 min (Table S2.4).

To estimate rates *a*_*i*_, *i* = 1 *…* 6, defined in (4), we assume that the equalities *b*_*x*_ = *b*_*y*_ = *b*_*u*_ = *b*_*w*_ = 10 can be approximately set by using RBS-related interventions [6]. Also, to avoid competition for ribosomes, only a few plasmids bearing promoters P_X_, P_Y_, P_G_, and P_R_ can be used. By selecting [P_X_] = [P_Y_] = [P_G_] = [P_R_] = 1 copies per cell, we obtain

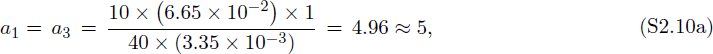

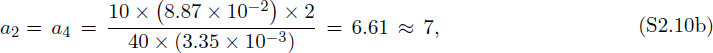

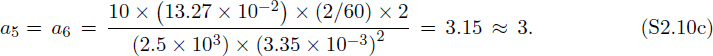

Next, from (5), we obtain

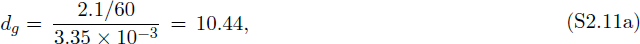

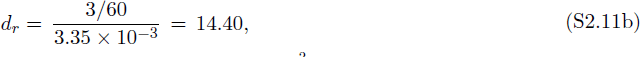

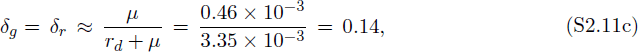

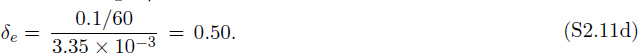

We find the estimated values of the parameters to be of the same order of magnitude as the corresponding parameter values estimated and used in [1,2,13,14]. Not enough is yet known about molecular interactions inside host cells to obtain highly precise descriptions [6]; it is common to computationally evaluate the effect of different values for rate parameters and even for Hill exponents [1,2,13,14]. Following [1,2,13,14], where genetic circuits built from similar elements have been studied, we have explored sets of parameter values which are close to the estimates given in (S2.10) and (S2.11), which ensure bistability in both Sand A-models, see the main text.

### S3 Exponential Stability of Cellular Populations

A systematic analysis of dynamical mathematical models begins with finding equilibrium solutions followed by the analysis of their exponential stability [24]. The next step is often to carry out (local) bifurcation analysis of the equilibrium solutions, allowing for the exploration of “stability boundaries” in the parameter space [53]. Both stability and bifurcation analyses rely on the computation of the eigenvalues from the corresponding model linearizations [53].

A nontrivial specificity of the computation of eigenvalues for the stability and bifurcation analyses of the Aand S-population models is that both models with *N* > 1 are invariant with respect to the action or the given linear representation of the *symmetric* group **S**_*N*_ of permutations among *N* -cells [52]. It is known that irreducible representations of groups enforce *multiple* eigenvalues of matrices that commute with their linear representations, a well-known fact following from Schur’s Lemma in the representation theory of Lie groups [52].

To take into account the necessity to deal with multiple eigenvalues in the situations when the value of *N* is *a priori* unknown, we have developed a general approach to the analysis of exponential stability [24] in arbitrary populations of identical cells, independently of *N*, as described below.

#### A General Population Model of Identical Cells

In this SI, we use Schur’s formula [66] to compute explicitly the characteristic polynomials for the corresponding model linearizations. The most important implication of Schur’s formula is that it can be easily seen that the values of the eigenvalues are independent of *N* ≥ 2. A conceptually similar result on the exponential orbital stability of limit cycles in a system of both identical and slightly different oscillators coupled via a medium was obtained by E. E. Shnol [88] in 1987. In his work, an averaging technique over the entire cellular population was used for both homogeneous and heterogeneous populations. Some results obtained in [88] were independently rediscovered in 2008 by G. Katriel [89] for homogeneous populations only, using Floquet Theory.

To describe the general exponential stability analysis, we first introduce an appropriate notation as follows. Let *S* and *z* be “generalized” global (extracellular) and local (intracellular) state variables, respectively, dim *S* = *m ≥* 1 and dim *z* = *k ≥* 2. Using the generalized variables, both the S-model (1) and the A-model (2) can then be rewritten in the following general form, which we call a G-model,

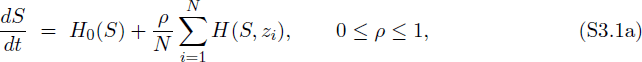

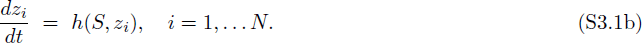

The G-model (S3.1) includes *m* + *N k* equations.

#### A Homogeneous Population

In the case of a homogeneous population of identical cells, we have *z*_*i*_(*t*) *≡ z*(*t*). As a result, the G-model (S3.1) reduces to a system of (*m* + *k*)-differential equations,

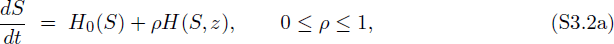

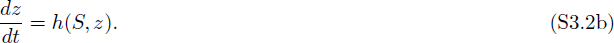

Observe that the model (S3.2) describes a *single* cell placed in a “free”, non-constant medium.

#### Definition 1.

*Let* (*S*_0_, *z*_0_) *be an equilibrium solution of the model* (S3.2). *Then,* (*S*_0_, *z*_0_) *corresponds to a homogeneous population equilibrium solution,*

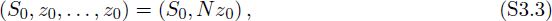

*of the full G-model* (S3.1) *for any N ≥* 2. *Notation N z*_0_ *means that z*_0_ *is repeated N -times in* (*S*_0_, *z*_0_, *…, z*_0_).

Although the model (S3.2) is sufficient to study the *existence* of homogeneous population equilibrium solutions (S3.3), it is not enough to establish the exponential *stability* of the corresponding solutions (S3.3). Let (*S*_0_, *N z*_0_) be a homogeneous population equilibrium solution of the G-model (S3.1) with any fixed *N* ≥ 2. To analyze the exponential stability of (*S*_0_, *N z*_0_) in the “full” G-model (S3.1), we need to compute the eigenvalues of the corresponding Jacobian matrix **J**_*N*_,

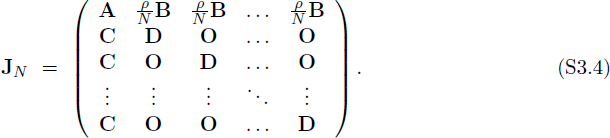

In (S3.4), each of three matrices, **B**, **C**, and **D**, is repeated *N* -times; **A** and **D** are square matrices of sizes *m* and *k*, respectively; **B** and **C** are rectangular matrices of sizes *m × k* and *k × m*, respectively,

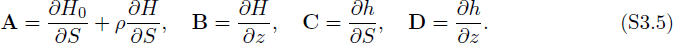

All partial derivatives in the expressions (S3.5) are evaluated at (*S*_0_, *z*_0_) which depends on all G-model parameters with the one important exception that they are *independent* of *N* because (*S*_0_, *z*_0_) is obtained using (S3.2). Notation **O** corresponds to zero submatrices of appropriate sizes.

We call a square matrix *stable* if all its eigenvalues have strictly negative real parts. The following theorem holds for J_*N*_.

#### Theorem 2.

(I). *Statements* (a), (b), *and* (c) *are equivalent*.

(a). *The matrix* **J***_N_ is stable for all N ≥* 2.

(b). *The matrix* **J**_1_ *and its submatrix* **D** *are both stable*.

(c). *The matrix* **J**_2_ *is stable*.

(II). *The matrix* **J***_N_ has typically k different eigenvalues, each of multiplicity N* 1 *in the following sense. Let {λ*_1_, *…, λ*_*m*+*k*_} *be the set of eigenvalues of matrix* **J**_1_, *and let {μ*_1_, *…, μ_k_} be the set of eigenvalues of its submatrix* **D**, *Then,*

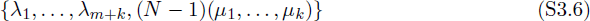

*is the set of all eigenvalues of matrix* **J***_N_ for any N ≥* 2, *where {μ*_1_, *…, μ_k_} is repeated* (*N* − 1)*-times*.

*Proof*. Let *λ* be a complex number, *λ* ∈ ℂ. Consider a new matrix **M***_λ_* = **J***_N_ λ***I**_*m*+*Nk*_, where **I**_*m*+*Nk*_ is the identity matrix of size *m* + *nk*. To find eigenvalues of **J**_*N*_, we need to write down the corresponding characteristic equation *P* (*λ*) = 0, *P* (*λ*) = det **M***_λ_*. Let us represent matrix **M***_λ_* in the form

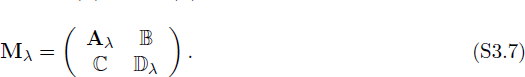

Here, matrices **A***_λ_* = **A***-λ***I**_*m*_, 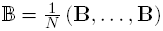, ℂ = (**C**, *… ,* **C**)^T^, and 𝔻*_λ_* = diag (**D***_λ_, … ,* **D***_λ_*) with **D***_λ_* = **D** *λ***I**_*k*_. Next, assume for a moment that 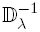s exists. Then, Schur’s formula can be used to compute det **M***_λ_* [66],

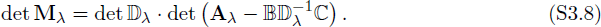

Next, we compute

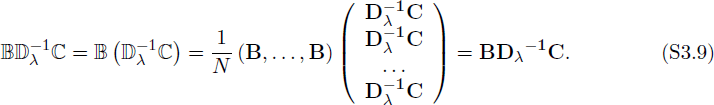

For the determinant of the block diagonal 𝔻*_λ_*, we obtain det 𝔻*_λ_* = (det **D***_λ_*)*^N^*. Substituting (S3.9) into (S3.8) yields

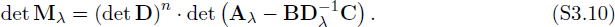

Using the Schur’s formula for the product det 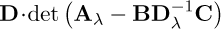 in the “backward” direction, we can rewrite (S3.10) in the following equivalent form

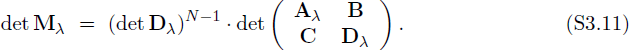

The expression (S3.11) can now be rewritten simply as

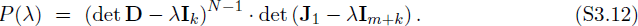

Recall that the expression (S3.12) has been proven under a restrictive condition det 𝔻*_λ_ ≠* 0, see above, which means that *λ* is not an eigenvalue of the matrix **D**. This restriction can be removed, for example, as follows. Let *λ*_0_ be an eigenvalue of the matrix **D**. Then, we obtain for the polynomial *P* (*λ*) by continuity

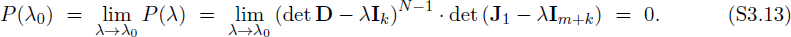

It follows from (S3.13) that (S3.12) holds for all *λ* ∈ ℂ.

Finally, we observe from (S3.12) that to compute all eigenvalues of the Jacobian matrix **J**_*N*_ for any *N* ≥ 2, it is sufficient to compute the eigenvalues of either two smaller matrices, **D** and **J**_1_, or one matrix **J**_2_. The latter may be practically slightly easier than computing the eigenvalues for **D** and **J**_1_ separately. The proof of the theorem follows.

Consider a differential equation

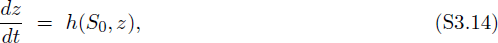

where *S*_0_ is a fixed parameter corresponding to the equilibrium (*S*_0_, *N × z*_0_) of the full Gmodel (S3.1). In contrast to equation (S3.2), equation (S3.14) describes a single cell placed into a constant environment, which can be interpreted as an environment shaped by the large population of cells and which does not “sense” any changes in a single cell. Additionally, consider a cascade model

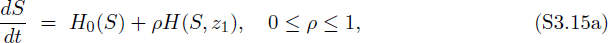

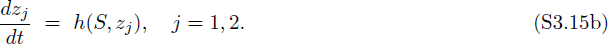

Observe that the variable *z*_2_ is absent from the first equation (S3.15a) and, hence, (S3.15) cannot be obtained from (S3.1) by simply setting *N* = 2.

Then, using the definition of exponential stability [24], the first statement of Theorem 2 can be reformulated as the following corollary which admits an intuitive interpretation of the fact why the case of *N* = 2 is sufficient to study the exponential stability of homogeneous population solutions.

#### Corollary 1.

*Let* (*S*_0_, *N × z*_0_) *be an equilibrium solution of the G-model* (S3.1). *Then, statements* (*a*) (*d*) *are equivalent*.

(a). (*S*_0_, *N × z*_0_) *is exponentially stable in the G-model* (S3.1) *for any N ≥* 2.

(b). (*S*_0_, *z*_0_) *is exponentially stable in the reduced model* (S3.2), *and z*_0_ *is exponentially stable in the single-cell model* (S3.14).

(c). (*S*_0_, *z*_0_, *z*_0_) *is exponentially stable in the G-model* (S3.1) *at N* = 2.

(*d*). (*S*_0_, *z*_0_, *z*_0_) *is exponentially stable in the cascade model* (S3.15).

A comparison of Statements *(a)* and *(b)* of Corollary 1 leads to a conclusion that the given population consisting of identical cells is stable with respect to any small perturbation if and only if *(i)* the population is stable with respect to any small *uniform* perturbation of the entire population described by system (S3.1) and, simultaneously, *(ii)* a majority of *unperturbed* cells forces a single *slightly perturbed* cell to re-join back the unperturbed majority.

Indeed, system (S3.14) used in Statement *(b)* means that the entire population does not sense small perturbations in a single cell because *S*_0_ is fixed in (S3.14).

Note that both conditions in Statement *(b)* can be reformulated, using the cascade model (S3.15) from statement (*d*). Finally, because the stability property is independent of the number *N* of identical cells in the population, the simple case of *N* = 2 can be used as given by statement *(c)*.

### A Mixed Population Split into Two Subpopulations

Suppose now that the given population consisting of *N*, *N* ≥ 4, identical cells is split into two different subpopulations of sizes *N*_1_ *≥* 2 and *N*_2_ *≥* 2, respectively, where *N* = *N*_1_ + *N*_2_. We always assume that each subpopulation consists of at least two cells. Then, the two different homogeneous subpopulations can be described by two state variables *z*_1_ and *z*_2_, respectively, where *z*_1_ ≠ *z*_2_, that is, *z*_*i*_*p*__ (*t*) *≡ z*_1_(*t*) for some subset of indexes *i*_*p*_, *p* = 1, *…, N*_1_, and *z*_*i*_*q*__ (*t*) *≡ z*_2_(*t*), for another subset of indexes *i*_*q*_, *q* = 1, *…, N*_2_. It follows that the equation (S3.1a) from the G-model (S3.1) simplifies as follows

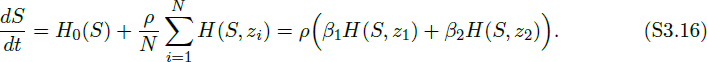

In (S3.16), *β_j_* is the fraction of the *j*-th subpopulation, *β_j_* = *N*_*j*_/*N*, *j* = 1, 2, *β*_1_ + *β*_2_ = 1. In this case, the entire G-model (S3.1) reduces to the following three equations

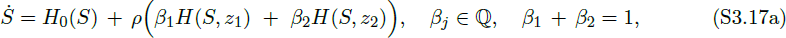

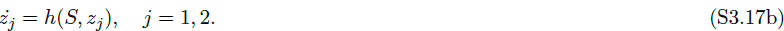

#### Definition 2.

*Let* (*S*_0_, *z*_10_, *z*_20_), *z*_10_ ≠ *z*_20_, *be a non-uniform equilibrium solution of the reduced system* (S3.17). *Then,* (*S*_0_, *z*_10_, *z*_20_), *z*_10_ *≠ z*_20_ *corresponds to a mixed population equilibrium solution,*

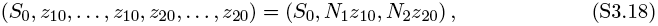

*of the full G-model* (S3.1). *The solution* (S3.18) *describes a mixed population of N identical cells, split into two* (*non-identical*) *subpopulations of sizes N*_1_ *>* 0 *and N*_2_ *>* 0, *respectively, N*_1_ + *N*_2_ = *N. Notation N*_*j*_*z*_*j*0_ *means that z*_*j*0_ *is repeated N_j_-times in* (*S*_0_, *z*_10_, *…, z*_10_, *z*_20_, *…, z*_20_), *j* = 1, 2.

Due to the condition *β*_1_ + *β*_2_ = 1 used in (S3.17a), there formally exists a *continuum* of different fractions *β*_1_ : *β*_2_, *β*_1_ *∈* ℝ and *β*_2_ *∈* ℝ. Of course, in the *biological* sense, only rational values *β*_1_ *∈* ℚ and *β*_2_ *∈* ℚ are allowable, leading to infinitely many fractional (*β*_1_ : *β*_2_)-configurations in the subdivision of the original population into two different subpopulations. Simple examples of such situations can be easily presented (Fig. S3.1).

**Figure S3.1.**
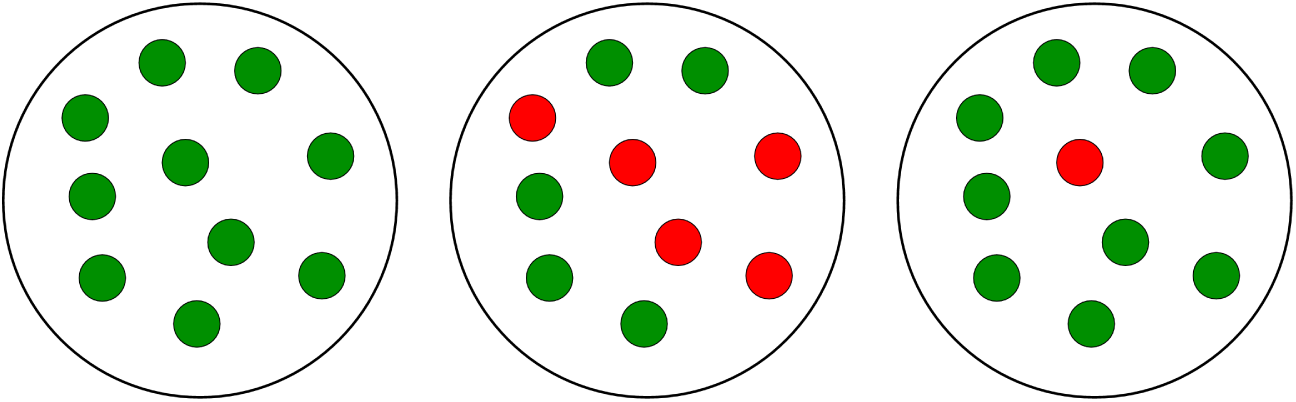
Examples of (*p* : *q*)-populations. The left panel corresponds to the case of a (10:0)-homogeneous population; the middle panel corresponds to the case of a (5:5)or, equivalently, (1:1)-mixed population, and the right panel corresponds to a (9:1)-mixed population.

For the sake simplicity of the exponential stability analysis, we will always assume that both *β*_1_ and *β*_2_ are real numbers, that is, *β_j_* ∈ ℝ, *j* = 1, 2.

Let (*S*_0_, *N*_1_*z*_10_, *N*_2_*z*_20_) be a mixed population equilibrium solution of the G-model (S3.1) with any fixed *N* ≥ 4, see (S3.3). To analyze the exponential stability of (*S*_0_, *N*_1_*z*_10_, *N*_2_*z*_20_), we need to compute the eigenvalues of the corresponding Jacobian matrix **J**_*N*_ obtained from the G-model

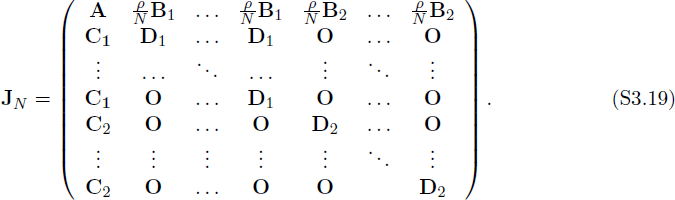

In the matrix (S3.19), submatrices **B**_*i*_, **C**_*i*_, and **D**_*i*_, are repeated *N*_*j*_-times; **A** and **D**_*i*_ are square matrices of sizes *m*, and *k*, respectively; **B**_*i*_ and **C**_*i*_ are rectangular matrices of sizes *m × k* and

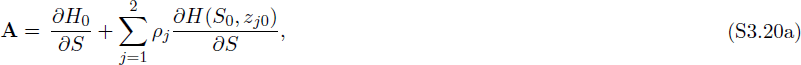

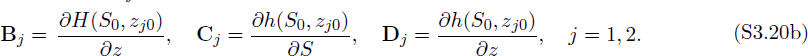

Consider the Jacobian matrix **Q**_2_ of size *m*+2*k* for the system (S3.17), computed at (*S*_0_, *z*_10_, *z*_20_),

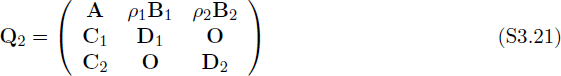

#### Theorem 3.

(I). *Statements* (a) *and* (b) *are equivalent*.

(a). *The matrix* **J***_N_ is stable for all N ≥* 4, *and with any N*_1_ *≥* 2 *and N*_2_ *≥* 2 *such that N*_1_ + *N*_2_ = *N*.

(b). *Matrix* **Q**_2_, *and its two submatrices,* **D**_1_ *and* **D**_2_, *are stable*.

(II). *Matrix* **J**_*N*_ has typically 2*k different multiple eigenvalues in the following sense. Let* {*λ*_1_, …, λ_*m*+2*k*_} *be the set of eigenvalues of* **Q**_2_, *let* {μ_1_, *…, μ*_*k*_} *be the set of eigenvalues of* **D**_1_, *and let {σ*_1_, *…, σ*_*k*_} *be the set of eigenvalues of* **D**_2_. *Then,*

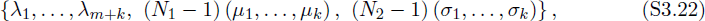

*is the set of all eigenvalues of the matrix* **J***_N_ for any N ≥* 4. *In* (S3.22), *the set {μ*_1_, *…, μ_k_} is repeated* (*N*_1_ − 1)*-times, and the set* {*σ*_1_, *…, σ_k_} is repeated* (*N*_2_ − 1)*-times. To have a nonzero value of multiplicity N_j_* − 1 *in* (S3.22), *condition N_j_ ≥* 2 *and, hence, N ≥* 4, *are natural requirements, j* = 1, 2. *In other words, the latter two conditions guarantee that both matrices* D_1_ *and* D_2_ *exist. Otherwise, Theorem 3 does not make any sense*.

*Proof*. The proof Theorem 3 can be carried out, using a simple modification of the proof of Theorem 2. For this reason, we only provide a brief sketch of the proof for Theorem 3. Similarly to the proof of Theorem 2, we need to write down a characteristic equation *P* (*λ*) = 0 Here *P* (*λ*) = det **M***_λ_*, and matrix **M***_λ_* can be defined as in (S3.7), using appropriate submatrices,

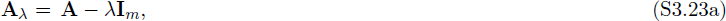

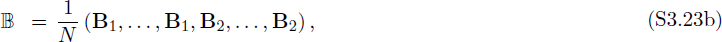

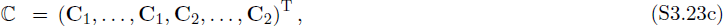

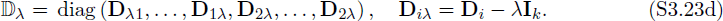

In the above submatrix definitions, the matrices with index *j* are repeated *N*_*j*_-times, *i, j* = 1, 2. In this case, det **M***_λ_* can also be computed using Schur’s formula, see (S3.8). However, (S3.9) should be replaced by

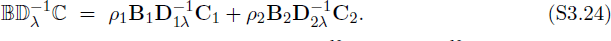

For the block diagonal matrix 𝔻*_λ_*, we obtain det 𝔻*_λ_* = (det **D**_1*λ*_)*^N^*^_1_^. (det **D**_2*λ*_)*^N^*^_2_^. Now, similarly to (S3.10), we will have

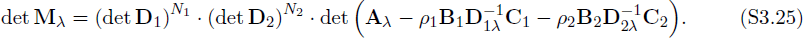

Using the Schur’s formula in the “backward” direction, we will then have

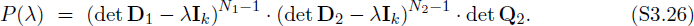

The rest can be proved as in the proof for Theorem 2. The proof of Theorem 3 follows.

Consider the following cascade model

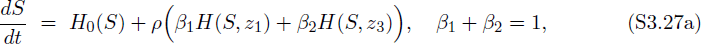

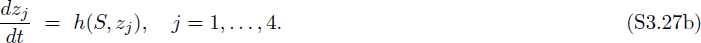

Variables *z*_2_ and *z*_4_ are absent from the first equation (S3.27a) and, hence, the cascade system (S3.27) cannot be obtained from the G-model (S3.1) by simply setting *N* = 4. Now, Theorem 3 can be reformulated in terms its Corollary 2 as follows.

#### Corollary 2.

*Let* (*S*_0_, *N*_1_ *× z*_10_, *N*_2_ *× z*_20_) *be a mixed population equilibrium solution of the G-model* (S3.1). *Then, Statements* (a) (c) *are equivalent*.

(a). (*S*_0_, *N*_1_ *× z*_10_, *N*_2_ *× z*_20_) *is exponentially stable in the G-model* (S3.1) *for any N ≥* 4, *and with any N*_1_ *≥* 2 *and N*_2_ *≥* 2 *such that N*_1_ + *N*_2_ = *N*.

(b). (*S*_0_, *z*_10_, *z*_20_) *is exponentially stable in the reduced model* (S3.17), *and each z*_*j*0_ *is exponentially stable in the single-cell model* (S3.14), *j* = 1, 2.

(c). (*S*_0_, *z*_10_, *z*_10_, *z*_20_, *z*_20_) *is exponentially stable in the cascade model* (S3.27).

### A Mixed Population Split into *L* Subpopulations

The case of a mixed population split into two subpopulations with densities *ρ*_1_ = *β*_1_*ρ* and *ρ*_2_ = *β*_2_*ρ* can be generalized to the case of a mixed population split into *L*-different subpopulations with densities *ρ*_1_, *…, ρ_L_*, where *ρ*_1_ + *…* + *ρ_L_* = *ρ*, *L ≥* 3 as follows.

Let subpopulation *j* consist of *N*_*j*_ cells, and let subpopulation *j* correspond to variable *z*_*j*_, that is, we have *z_j_′* (*t*) *≡ z_j_*(*t*), where *j′* ∈ {*i*_1_, *i*_2_, *…, i_N_j__ } ⊂ {*1, 2, *… N }*, *j* = 1, *…, L*. In this case, the G-model (S3.1) reduces to the following equations

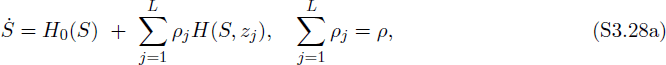

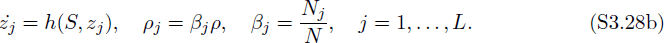

#### Definition 3.

*Let* (*S*_0_, *z*_10_, *…, z*_*L*0_) *be a non-uniform equilibrium solution of the system* (S3.28), *where z*_*j*0_ *≠ z*_*j*_*′*0 *for all j ≠ j*′. *Then,* (*S*_0_, *z*_10_, …, *z*_*L*0_), *corresponds to a mixed population equilibrium solution,*

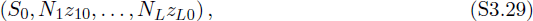

*of the full G-model* (S3.1). *The solution* (S3.29) *describes a mixed population of N identical cells, which is split into L subpopulations of the corresponding sizes N_j_ ≥* 2, *N*_1_ + *…* + *N*_*L*_ = *N*. Notation *N*_*j*_*z*_*j*0_ *means that z*_*j*0_ *is repeated N*_*j*_*-times in the vector-form solution of the the full G-model* (S3.1), *N*_*j*_ ≥ 2, *j* = 1, *…, L*.

Consider the Jacobian matrix **Q**_*L*_ for the reduced system (S3.28), computed at (*S*_0_, *z*_10_, …, *z*_*L*0_),

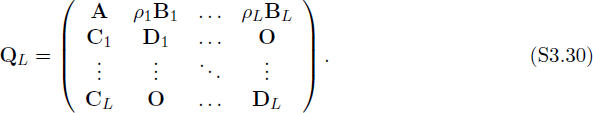

In (S3.30), all submatrices are defined as in (S3.20), where *j* = 1, 2 should be replaced by *j* = 1, *…, L*. Below, we formulate Theorem 4 and Corollary 3 without any proof because they are similar to Theorem 3 and Corollary 1, respectively.

#### Theorem 4.

(I). *Statements* (a) *and* (b) *are equivalent*.

(a). *The Jacobian* **J***_N_ computed for the G-model at the given equilibrium* (S3.29) *is stable for all N ≥* 2*L, and with any N_j_ ≥* 2, *j* = 1, *…, L, such that N*_1_ + *… N_L_* = *N*.

(b). *Matrix* **Q***_L_ and its submatrices* **D***_j_, j* = 1, *…, L, are stable*.

(II). *Matrix* **J***_N_ has typically kL different multiple eigenvalues in the following sense. Let* {*λ*_1_, …, *λ*_*m*+*kL*_} *be the set of eigenvalues of* **Q**_*L*_, *and let 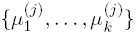 be the set of eigenvalues of* **D**_*j*_, *j* = 1, …, *L. Then,*

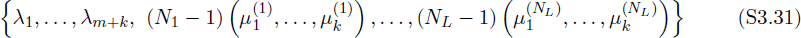

*is the set of all eigenvalues of the matrix* **J***_N_ for any N ≥* 2*L. In* (S3.31), *each set 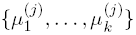 is repeated* (*N*_*j*_ − 1)*-times with all N_j_ ≥* 2, *j* = 1, *…, K*.

Consider the following cascade model

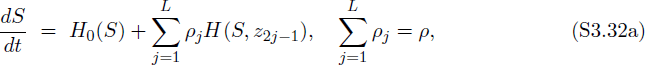

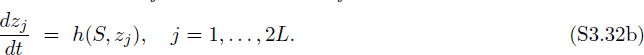

State variables *z*_2*j*_ with even indeces are absent from the first equation (S3.32a) of the cascade model (S3.32).

#### Corollary 3.

*Let* (*S*_0_, *N*_1_*z*_10_, …, *N*_*L*_*z*_*L*0_) *be a mixed equilibrium solution of the G-model* (S3.1), where *N*_*j*0_ *≠ N*_*j*_*′*0 for all *j* ≠ *j*′. *Then, Statements* (a) - (c) *are equivalent*.

(a). (*S*_0_, *N*_1_*z*_10_, …, *N*_*L*_*z*_*L*0_) *is exponentially stable in the G-model* (S3.1) *for any N ≥* 2*L, and with any N_j_ ≥* 2, *j* = 1, *…, L, such that N*_1_ + *… N_L_* = *N*.

(b). (*S*_0_, *z*_10_, …, *z*_*L*0_) *is exponentially stable in the reduced model* (S3.28), and each *z*_*j*0_ *is exponentially stable in the single-cell model* (S3.14), *j* = 1, *…, L*.

(c). (*S*_0_, *z*_10_, *z*_10_, *z*_20_, *z*_20_, *…, z*_*L*0_, *z*_*L*0_) *is exponentially stable in the cascade model* (S3.32).

## Acknowledgments

The work was supported in part by grants AFOSR FA9550-14-1-0060 and ONR 5710003367.

## References

1. Gardner TS, Cantor CR, Collins JJ. Construction of a genetic toggle switch *In Escherichia, coli*. Nature. 2000;403(6767):339–342.

2. Elowitz MB, Leibler S. A synthetic oscillatory network of transcriptional regulators. Nature. 2000;403(6767):335–338.

3. Hasty J, Pradines J, Dolnik M, Collins JJ. Noise-based switches and amplifiers for gene expression. Proceedings of the National Academy of Sciences. 2000;97(5):2075–2080.

4. Kobayashi H, Kærn M, Araki M, Chung K, Gardner TS, Cantor CR, et al. Programmable cells: interfacing natural and engineered gene networks. Proceedings of the National Academy of Sciences of the United States of America. 2004;101(22):8414–8419.

5. Purnick PE, Weiss R. The second wave of synthetic biology: from modules to systems. Nature reviews Molecular cell biology. 2009;10(6):410–422.

6. Arpino JA, Hancock EJ, Anderson J, Barahona M, Stan GBV, Papachristodoulou A, et al. Tuning the dials of synthetic biology. Microbiology. 2013;159(Pt 7):1236–1253.

7. Nadell CD, Bassler BL, Levin SA. Observing bacteria through the lens of social evolution. Journal of biology. 2008;7:27.

8. Miller MB, Bassler BL. Quorum sensing in bacteria. Annual Reviews in Microbiology. 2001;55(1):165–199.

9. Pesci EC, Pearson JP, Seed PC, Iglewski BH. Regulation of *las* and *rhl* quorum sensing in Pseudomonas aeruginosa. Journal of bacteriology. 1997;179(10):3127–3132.

10. Lithgow JK, Wilkinson A, Hardman A, Rodelas B, Wisniewski-Dye F, Williams P, et al. The regulatory locus cinRI *in Rhizobium leguminosarum* controls a network of quorum-sensing loci. Molecular microbiology. 2000;37(1):81–97.

11. McAnulla C, Edwards A, Sanchez-Contreras M, Sawers RG, Downie JA. Quorum-sensing-regulated transcriptional initiation of plasmid transfer and replication genes *in Rhizobium leguminosarum biovar viciae*. Microbiology. 2007;153(7):2074–2082.

12. Garcia-Ojalvo J, Elowitz MB, Strogatz SH. Modeling a synthetic multicellular clock: repressilators coupled by quorum sensing. Proceedings of the National Academy of Sciences of the United States of America. 2004;101(30):10955–10960.

13. Kuznetsov A, Kærn M, Kopell N. Synchrony in a population of hysteresis-based genetic oscillators. SIAM Journal on Applied Mathematics. 2004;65(2):392–425.

14. Wang J, Zhang J, Yuan Z, Zhou T. Noise-induced switches in network systems of the genetic toggle switch. BMC systems biology. 2007;1(1):50.

15. Smale S. On the differential equations of species in competition. Journal of Mathematical Biology. 1976;3:5–7.

16. Hirsch MW. The dynamical systems approach to differential equations. Bull AMS. 1984;11:1–64.

17. Smith HL. Monotone Dynamical Systems, Vol. 41. Mathematical Surveys and Monographs. 1995;.

18. Angeli D, Sontag ED. Monotone control systems. IEEE Trans Automat Control. 2003;48(10):1684–1698.

19. Hirsch M, Smith H. Monotone dynamical systems. Handbook of differential equations: ordinary differential equations. 2005;2:239–357.

20. Sontag ED. Monotone and near-monotone biochemical networks. Systems and Synthetic Biology. 2007;1:59–87.

21. Smith HL. Monotone dynamical systems: an introduction to the theory of competitive and cooperative systems. vol. 41. American Mathematical Soc.; 2008.

22. Parsek MR, Greenberg EP. Acyl-homoserine lactone quorum sensing in gram-negative bacteria: a signaling mechanism involved in associations with higher organisms. Proceedings of the National Academy of Sciences. 2000;97(16):8789–8793.

23. Resmi V, Ambika G, Amritkar R. Synchronized states in chaotic systems coupled indirectly through a dynamic environment. Physical Review E. 2010;81(4):046216.

24. Sontag ED. Mathematical control theory: deterministic finite dimensional systems. vol. 6. Springer; 1998.

25. Parsek MR, Val DL, Hanzelka BL, Cronan JE, Greenberg E. Acyl homoserine-lactone quorum-sensing signal generation. Proceedings of the National Academy of Sciences. 1999;96(8):4360–4365.

26. Hooshangi S, Thiberge S, Weiss R. Ultrasensitivity and noise propagation in a synthetic transcriptional cascade. Proceedings of the National Academy of Sciences of the United States of America. 2005;102(10):3581–3586.

27. Tuttle LM, Salis H, Tomshine J, Kaznessis YN. Model-driven designs of an oscillating gene network. Biophysical journal. 2005;89(6):3873–3883.

28. Stricker J, Cookson S, Bennett MR, Mather WH, Tsimring LS, Hasty J. A fast, robust and tunable synthetic gene oscillator. Nature. 2008;456(7221):516–519.

29. Hussain F, Gupta C, Hirning AJ, Ott W, Matthews KS, Josić K, et al. Engineered temperature compensation in a synthetic genetic clock. Proceedings of the National Academy of Sciences. 2014;111(3):972–977.

30. Laffend L, Shuler M. Structured model of genetic control via the *lac* promoter *in Escherichia coli*. Biotechnology and bioengineering. 1994;43(5):399–410.

31. Lutz R, Bujard H. Independent and tight regulation of transcriptional units *in Escherichia coli* via the LacR/O, the TetR/O and AraC/I1-I2 regulatory elements. Nucleic Acids research. 1997;25(6):1203–1210.

32. Baumeister R, Flache P, Melefors Ö, von Gabain A, Hillen W. Lack of a 5’non-coding region in Tn 1721 encoded *tetR* mRNA is associated with a low efficiency of translation and a short half-life *In Escherichia coil*. Nucleic acids research. 1991;19(17):4595–4600.

33. Markiewicz P, Kleina LG, Cruz C, Ehret S, Miller JH. Genetic studies of the lac repressor. XIV. Analysis of 4000 altered *Escherichia coli* lac repressors reveals essential and nonessential residues, as well as “spacers” which do not require a specific sequence. Journal of molecular biology. 1994;240(5):421–433.

34. Ramos JL, Martínez-Bueno M, Molina-Henares AJ, Terán W, Watanabe K, Zhang X, et al. The TetR family of transcriptional repressors. Microbiology and Molecular Biology Reviews. 2005;69(2):326–356.

35. Semsey S, Jauffred L, Csiszovszki Z, Erdőssy J, Stéger V, Hansen S, et al. The effect of LacI autoregulation on the performance of the lactose utilization system *in Escherichia coli*. Nucleic acids research. 2013;p. gkt351.

36. Gray KM, Garey JR. The evolution of bacterial LuxI and LuxR quorum sensing regulators. Microbiology. 2001;147(8):2379–2387.

37. Shuler M, Leung S, Dick C. A mathematical model for the growth of a single bacterial cell. Annals of the New York Academy of Sciences. 1979;326(1):35–52.

38. Neidhardt FC, Ingraham JL, Schaechter M. Physiology of the Bacterial Cell: A Molecular Approach. Sinauer Associates Sunderland, MA; 1990.

39. Kim BG, Shuler M. A structured, segregated model for genetically modified *Escherichia coli* cells and its use for prediction of plasmid stability. Biotechnology and bioengineering. 1990;36(6):581–592.

40. Shu J, Shuler M. Prediction of effects of amino acid supplementation on growth of *E. coli* B/r. Biotechnology and bioengineering. 1991;37(8):708–715.

41. Laffend L, Shuler M. Ribosomal protein limitations *in Escherichia coli* under conditions of high translational activity. Biotechnology and bioengineering. 1994;43(5):388–398.

42. Bailey JE. Mathematical modeling and analysis in biochemical engineering: past accomplishments and future opportunities. Biotechnology progress. 1998;14(1):8–20.

43. Domach M, Leung S, Cahn R, Cocks G, Shuler M. Computer model for glucose-limited growth of a single cell of *Escherichia coli* B/r-A. Biotechnology and bioengineering. 2000;67(6):827–840.

44. Nikolaev EV, Atlas JC, Shuler ML. Computer models of bacterial cells: from generalized coarsegrained to genome-specific modular models. In: Journal of Physics: Conference Series. vol. 46. IOP Publishing; 2006. p. 322.

45. Atlas J, Nikolaev E, Browning S, Shuler M. Incorporating genome-wide DNA sequence information into a dynamic whole-cell model of *Escherichia coli*: Application to DNA replication. Systems Biology, IET. 2008;2(5):369–382.

46. Lodish H, Berk A, Kaiser CA, Krieger M, Bretscher A, Ploegh H, et al. Molecular cell biology. Macmillan; 2012.

47. Dockery JD, Keener JP. A mathematical model for quorum sensing *in Pseudomonas aeruginosa*. Bulletin of mathematical biology. 2001;63(1):95–116.

48. Bressloff PC. Stochastic processes in cell biology. vol. 41. Springer; 2014.

49. Dong YH, Wang LH, Xu JL, Zhang HB, Zhang XF, Zhang LH. Quenching quorum-sensing-dependent bacterial infection by an N-acyl homoserine lactonase. Nature. 2001;411(6839):813–817.

50. Waldherr S, Allgöwer F. Network-level dynamics of diffusively coupled cells. 51st IEEE Conference on Decision and Control December 10-13, Maui, Hawaii, USA. 2012;.

51. Farrell CM, Baker TA, Sauer RT. Altered specificity of a AAA+ protease. Molecular cell. 2007;25(1):161–166.

52. Golubitsky M, Schaeffer DG, Stewart I. Singularities and groups in bifurcation theory. vol. 2. Springer New York; 1988.

53. Kuznetsov YA. Elements of applied bifurcation theory. vol. 112. Springer Science & Business Media; 2013.

54. Krupa M. Bifurcations of relative equilibria. SIAM journal on mathematical analysis. 1990;21(6):1453–1486.

55. Hirsch M. Differential equations and convergence almost everywhere in strongly monotone flows. Contemporary Mathematics. 1983;17:267–285.

56. Hirsch M. Systems of differential equations that are competitive or cooperative II: Convergence almost everywhere. SIAM J Mathematical Analysis. 1985;16:423–439.

57. Smith H. Monotone dynamical systems: An introduction to the theory of competitive and cooperative systems, Mathematical Surveys and Monographs, vol. 41. Providence, RI: AMS; 1995.

58. Hadeler K, Glas D. Quasimonotone systems and convergence to equilibrium in a population genetics model. J Math Anal Appl. 1983;95:297–303.

59. Hirsch M, Smith H, et al. Monotone dynamical systems. Handbook of differential equations: ordinary differential equations. 2005;2:239–357.

60. Zaslavsky T. Bibliography of signed and gain graphs. Electronic Journal of Combinatorics. 1998;DS8.

61. Harary F. On the notion of balance of a signed graph. Michigan Mathematical Journal. 1953;2:143–146.

62. Dhooge A, Govaerts W, Kuznetsov YA. MATCONT: a MATLAB package for numerical bifurcation analysis of ODEs. ACM Transactions on Mathematical Software (TOMS). 2003;29(2):141–164.

63. Govaerts W, Kuznetsov YA, Dhooge A. Numerical continuation of bifurcations of limit cycles in MATLAB. SIAM journal on scientific computing. 2005;27(1):231–252.

64. Khibnik AI, Kuznetsov YA, Levitin VV, Nikolaev EV. Continuation techniques and interactive software for bifurcation analysis of ODEs and iterated maps. Physica D: Nonlinear Phenomena. 1993;62(1):360–371.

65. Vasil’eva AB, Butuzov VF, Kalachev LV. The boundary function method for singular perturbation problems. SIAM; 1995.

66. Gantmakher FR. The theory of matrices. vol. 131. American Mathematical Soc.; 1959.

67. Bolouri H. Computational modeling of gene regulatory networks: a primer. World Scientific; 2008.

68. Sauro HM. Enzyme kinetics for systems biology. Future Skill Software; 2011.

69. Cornish-Bowden A. Fundamentals of enzyme kinetics. John Wiley & Sons; 2013.

70. Garde C, Bjarnsholt T, Givskov M, Jakobsen TH, Hentzer M, Claussen A, et al. Quorum sensing regulation *in Aeromonas hydrophila*. Journal of molecular biology. 2010;396(4):849–857.

71. Shuler M. Single-cell models: promise and limitations. Journal of biotechnology. 1999;71(1):225–228.

72. Castellanos M, Wilson DB, Shuler ML. A modular minimal cell model: purine and pyrimidine transport and metabolism. Proceedings of the National Academy of Sciences of the United States of America. 2004;101(17):6681–6686.

73. Segel IH. Biochemical Calculations: How To Solve Mathematical Problems In General Biochemistry. Wiley; 1976.

74. Bhattacharya BS, Sweby PK, Minihane AM, Jackson KG, Tindall MJ. A mathematical model of the sterol regulatory element binding protein 2 cholesterol biosynthesis pathway. Journal of Theoretical Biology. 2014;p. 150–162.

75. Bernstein JA, Khodursky AB, Lin PH, Lin-Chao S, Cohen SN. Global analysis of mRNA decay and abundance in Escherichia coli at single-gene resolution using two-color fluorescent DNA microarrays. Proceedings of the National Academy of Sciences. 2002;99(15):9697–9702.

76. Prindle A, Selimkhanov J, Li H, Razinkov I, Tsimring LS, Hasty J. Rapid and tunable post-translational coupling of genetic circuits. Nature. 2014;.

77. Thomas PW, Stone EM, Costello AL, Tierney DL, Fast W. The quorum-quenching lactonase from *Bacillus thuringiensis* is a metalloprotein. Biochemistry. 2005;44(20):7559–7569.

78. Raj M. PCR Amplification, and Sequence Comparison of lacI gene in WT *E. coli* C29 cells and a presumptive *lacI* Knockout *E. coli* C29 cells to Determine the Difference in the Basal Expression Level of *lacZ* in Lac Operon. J Exp Microbiol Immunol. 2004;6:13–19.

79. Postle K, Nguyen TT, Bertrand KP. Nucleotide sequence of the repressor gene of the TN10 tetracycline resistance determinant. Nucleic acids research. 1984;12(12):4849–4863.

80. Wang Y, Tegenfeldt JO, Reisner W, Riehn R, Guan XJ, Guo L, et al. Single-molecule studies of repressor-DNA interactions show long-range interactions. Proceedings of the National Academy of Sciences of the United States of America. 2005;102(28):9796–9801.

81. Pai A, You L. Optimal tuning of bacterial sensing potential. Molecular systems biology. 2009;5(1).

82. Chen CC, Riadi L, Suh SJ, Ohman DE, Ju LK. Degradation and synthesis kinetics of quorum-sensing autoinducer *in Pseudomonas aeruginosa* cultivation. Journal of biotechnology. 2005;117(1):1–10.

83. Flynn JM, Levchenko I, Seidel M, Wickner SH, Sauer RT, Baker TA. Overlapping recognition determinants within the ssrA degradation tag allow modulation of proteolysis. Proceedings of the National Academy of Sciences. 2001;98(19):10584–10589.

84. Selinger DW, Saxena RM, Cheung KJ, Church GM, Rosenow C. Global RNA half-life analysis *in Escherichia coli* reveals positional patterns of transcript degradation. Genome research. 2003;13(2):216–223.

85. Ochsner UA, Reiser J. Autoinducer-mediated regulation of rhamnolipid biosurfactant synthesis *in Pseudomonas aeruginosa*. Proceedings of the National Academy of Sciences. 1995;92(14):6424–6428.

86. Takaya A, Tabuchi F, Tsuchiya H, Isogai E, Yamamoto T. Negative regulation of quorum-sensing systems *in Pseudomonas aeruginosa* by ATP-dependent Lon protease. Journal of bacteriology. 2008;190(12):4181–4188.

87. Brint JM, Ohman DE. Synthesis of multiple exoproducts *in Pseudomonas aeruginosa* is under the control of RhlR-RhlI, another set of regulators in strain PAO1 with homology to the autoinducer-responsive LuxR-LuxI family. Journal of Bacteriology. 1995;177(24):7155–7163.

88. Shnol E. The synchronization of oscillators which interact via a medium. Journal of Applied Mathematics and Mechanics. 1987;51(1):9–13.

89. Katriel G. Synchronization of oscillators coupled through an environment. Physica D: Nonlinear Phenomena. 2008;237(22):2933–2944.

